# Small molecule G-quadruplex ligands are antibacterial candidates for Gram-negative bacteria

**DOI:** 10.1101/2022.09.01.506212

**Authors:** Yuiko Takebayashi, Javier Ramos-Soriano, Y. Jennifer Jiang, Jennifer Samphire, Efres Belmonte-Reche, Michael P. O’Hagan, Catherine Gurr, Kate J. Heesom, Philip A. Lewis, Thanadon Samernate, Poochit Nonejuie, James Spencer, M. Carmen Galan

**Affiliations:** School of Cellular & Molecular Medicine, Biomedical Sciences Building, BS8 1TD, University of Bristol; School of Chemistry, Cantock’s Close, BS8 1TS, University of Bristol; Centre for Genomics and Oncological Research (GENYO), Avenida de la Ilustración 114, 18016 Granada, Spain; Department of Biochemistry and Molecular Biology II, Faculty of Pharmacy, University of Granada, Granada, Spain; Instituto de Investigación Biosanitaria ibs.GRANADA, Hospital Virgen de las Nieves, Granada, Spain; Proteomics Facility, Biomedical Sciences Building, BS8 1TD, University of Bristol; Institute of Molecular Biosciences, Mahidol University, Nakhon Pathom, Thailand

**Keywords:** G-quadruplex DNA, G-quadruplex ligand, antimicrobial, therapeutics, G4 supramolecular targeting

## Abstract

There is great need for novel strategies to tackle antimicrobial resistance, in particular in Gram-negative species such as *Escherichia coli* that cause opportunistic infections of already compromised patients. Here we demonstrate, following a screen of G-quadruplex (G4) ligand candidates, that a novel pyridinium-functionalized azobenzene **L9** shows promising antibacterial activity (MIC values ≤ 4 μg/mL) against multi-drug resistant *E. coli*. Tandem Mass Tag (TMT) proteomics of *E. coli* treated with sub-lethal concentrations of **L9**, identified that, consistent with its superior antibacterial activity, **L9** treatment influences expression levels of more G4-associated proteins than the analogous ligands **L5** (stiff-stilbene) or pyridostatin (**PDS)**, and upregulates multiple essential proteins involved in translation. Biophysical analysis showed **L9** binds potential target G4-containing sequences, identified from proteomic experiments and by bioinformatics, with variable affinity, in contrast to the two comparator G4 ligands (**L5, PDS**) that better stabilize G4 structures but have lower antimicrobial activity. Fluorescence microscopy-based Bacterial Cytological Profiling (BCP) suggests that the **L9** mechanism of action is distinct from other antibiotic classes. These findings support strategies discovering potential G4 ligands as antibacterial candidates for priority targets such as multi-drug resistant *E. coli*, warranting their further exploration as potential novel therapeutic leads with G4-mediated modes of action.

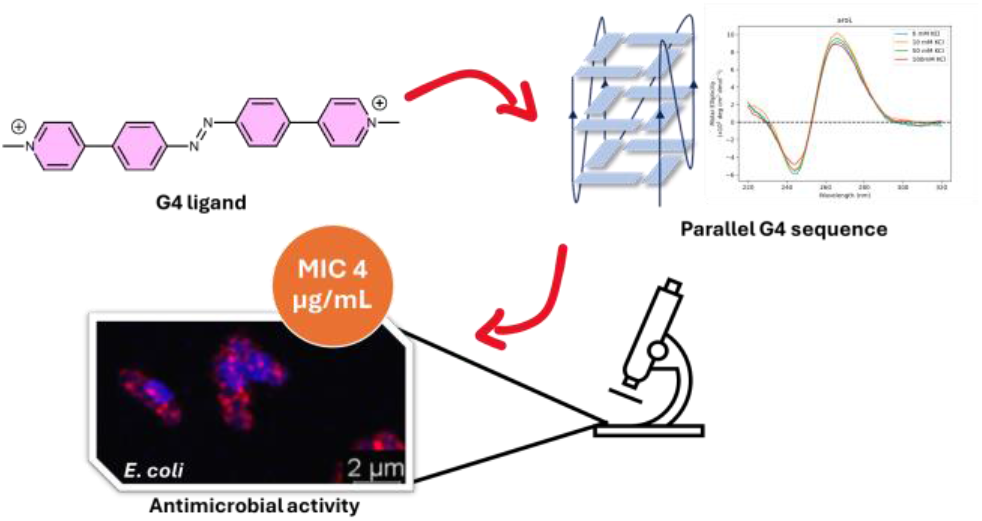

---

Antimicrobial resistance (AMR) is a 21^st^ century global public health emergency. With the continuing weakness of the antibacterial pipeline and decades of empirical antibiotic overuse^1^, once readily treatable infections now cause increasing morbidity and mortality, making the search for alternative agents urgent. Amongst bacterial pathogens, the World Health Organization (WHO) has identified carbapenem-resistant and extended spectrum beta-lactamase (ESBL)-producing *Enterobacterales* (Gram-negative bacteria including *Escherichia* and *Klebsiella* species) as critical, highest priority pathogens on which research and development of new antibiotics should be focused.^2^

G-quadruplexes (G4s) are secondary structures that form in guanine-rich regions of DNA and RNA in both eukaryotes and prokaryotes.^3, 4^ G4s form within both genes and regulatory regions of DNA and mRNA; including telomeres, promoter sites, transcriptional start sites (TSS) and 5’ and 3’ untranslated regions (UTRs), and consequently influence gene regulation and expression.^5, 6^ The prevalence of G4s, along with the growing realization of their importance in human disease progression (e.g. cancer, diabetes and neurogenerative diseases), and involvement in cellular processes in a wider range of organisms including plants, parasites, fungi, protozoa, bacteria and viruses; has excited considerable interest in their potential as therapeutic targets.^7^

Compared to mammalian systems, G4s in bacteria remain understudied. A 2006 *in silico* study of 18 prokaryotes, including *Escherichia coli, Pseudomonas aeruginosa* and *Mycobacterium bovis*, observed enrichment of G4s in regulatory regions within 200 bp upstream of genes involved in secondary metabolite biosynthesis, transcription and signal transduction.^8^ More recent whole genome sequencing studies have identified hundreds of G4 sites in the DNA and RNA of *E. coli*.^9, 10^ Moreover, in *Mycobacterium tuberculosis* (Mtb), G4 sites were identified in the coding regions of genes responsible for virulence in host cells (*espK, espB*) and essential to survival by maintenance of membrane fluidity (*cyp51*).^11^ The well characterized G4 ligand TMPyP4 ((5,10,15,20-Tetrakis-(*N*-methyl-4-pyridyl)porphine,^12^ an inhibitor of telomerase activity in cancer, stabilizes these secondary structures and downregulates expression of these three genes, leading to inhibition of Mtb growth (IC_50_ = 6.25 µM). In *Streptococcus pneumoniae*, essential genes involved in DNA repair (*recD*), drug efflux (*pmrA*) and virulence phase regulation (*hsdS*) all harbored intragenic G4 sites. TMPyP4 both stabilized these sequences *in vitro* and inhibited expression of a fluorescent reporter protein in the human HEK 293 cell line when the same sequences were incorporated in the upstream promoter.^13^ In multi-drug resistant *Klebsiella pneumoniae*, G4 sites in the promoters of five essential genes were stabilized by another well-known anti-viral G4 ligand BRACO-19, which decreased expression of their respective RNAs at a concentration of 5.38 µM and inhibited growth with an IC_50_ of 10.77 µM.^14^ These studies, although limited to a few species, highlight the potential of G4 motifs as targets for candidate antibacterials able to disrupt cellular function and viability. However, to date G4 ligands with potent antimicrobial activity that act upon specific microbial G4 sequences have not been identified and validated. To investigate the potential antibacterial activity of small molecules that bind to G4 sequences, we used the agar disc susceptibility method to screen a selection of new and known candidate G4 ligands (**L1** - **L11**,^15-18^ **Figure S1**, see SI for synthetic details), and the comparators TMPyP4,^19^ BRACO-19^20^ and pyridostatin (**PDS**), against *E. coli* and *Staphylococcus aureus* as exemplar Gram-negative and Gram-positive target pathogens (**Table S1**). Of the tested ligands, two (**L5, L9, Figure 1**) exhibited activity against *E. coli* and 6 (**L3, L5**-**L9**) against *S. aureus*. Minimal inhibitory concentrations (MICs) were then determined for the *S. aureus* Newman and *E. coli* ATCC 25922, UTI 808 and IR60 strains by broth microdilution. MICs ranged from 1 – >512 µg/mL for the 6 ligands against *S. aureus*, and 2 – 128 µg/mL for the two ligands active against *E. coli* (**Table S2**). Of the tested ligands, **L9** was selected for further study due to its activity against the multidrug resistant (MDR) clinical *E. coli* isolates UTI 808 and IR60. **L5** was selected as a less potent comparator due to its structural similarity to **L9**.

**Figure 1.**
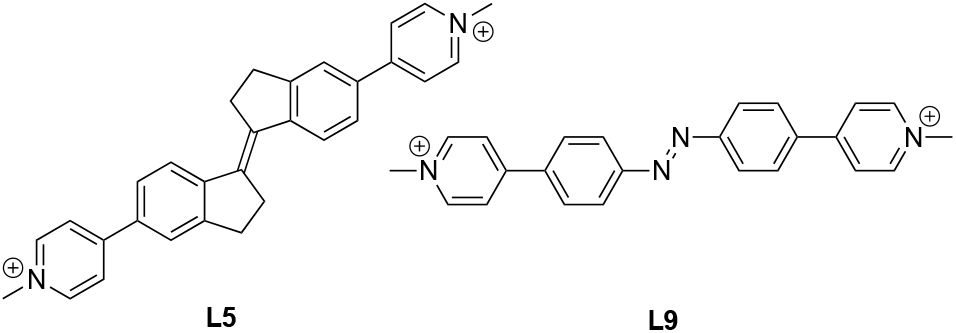
Structures of Candidate G-Quadruplex Ligands L5 and L9. (See SI for structures of all ligands tested).

**L5, L9** and **PDS** were screened against a wider range of Gram-negative organisms, representing both laboratory and MDR clinical strains (**Table 1**). **L9** showed selective activity against *E. coli* compared to *K. pneumoniae, P. aeruginosa* and *Acinetobacter baumannii*. To assess whether membrane permeability was a factor influencing the specific activity of **L9** against *E. coli*, MIC assays were repeated with addition of the membrane disrupting agents, polymyxin B nonapeptide (PMBN) and colistin, the former having no antibacterial activity against Enterobacterales.^21^ No significant decrease in MIC values (defined by MIC fold change >2) of **L5, L9** and **PDS** towards the *E. coli* strains was observed on addition of PMBN, with the exception of **L5** against IR60. Notable decreases in MIC values of **L5** and **L9** were observed for *K. pneumoniae* NCTC 5055 and *P. aeruginosa* PAO1 and 4623, suggesting that these compounds less readily penetrate the envelopes of these species compared to *E. coli*. Changes in MIC values of **L5** and **L9** were not significant for *A. baumannii* strains. PMBN did not affect the MIC of **PDS** against all tested organisms. However, colistin addition decreased the MIC of all three ligands by 8 – 32-fold in *E. coli*, highlighting the possibility that these G4 ligands may be potentiated by existing antibiotics.

**Table 1.** Minimal Inhibitory Concentration (MIC) of L5, L9 and pyridostatin (PDS) ± polymyxin B nonapeptide (PMBN) or colistin (COL) against selected Gram-negative bacteria.

|  | MIC (µg/mL) |  |  |  |  |  |
| --- | --- | --- | --- | --- | --- | --- |
|  | **L5** |  | **L9** |  | **PDS** |  |
|  |  | +PMBN<br>(+COL) |  | +PMBN<br>(+COL) |  | +PMBN<br>(+COL) |
| *E. coli*<br>ATCC 25922 | 64 | 32<br>(8) | 4 | 2<br>(0.25) | 64 | 128<br>(4) |
| *E. coli*<br>UTI 808 | 128 | 64<br>(8) | 4 | 4<br>(0.5) | 64 | 64<br>(4) |
| *E. coli*<br>IR60 | 128 | 32<br>(8) | 2 | 2<br>(1) | 64 | 64<br>(2) |
| *K. pneumoniae*<br>NCTC 5055 | 256 | 64<br>(16) | >256 | 64<br>(2-32)* | 64 | 64<br>(64) |
| *K. pneumoniae*<br>KP11 | >512 | 256<br>(32) | >256 | >256<br>(256) | 64 | 32<br>(64) |
| *P. aeruginosa*<br>PAO1 | >512 | 128<br>(64) | >256 | 32<br>(128) | 64 | 64<br>(64) |
| *P. aeruginosa*<br>4623 | >512 | 128<br>(64) | >256 | 32<br>(32) | 64 | 64<br>(64) |
| *A. baumannii* | 128 | 64 | 64 | 32 | 32 | 64 |

| ATCC 19606 |  | (32) |  | (64) |  | (32) |
| --- | --- | --- | --- | --- | --- | --- |
| *A. baumannii* | 128 | 128 |  | >256 |  | 64 |
| 6490A |  | (64) | >256 | (128) | 32 | (32) |

\* Reported as a range due to highly variable MIC values (n=6).

Next, the growth of *E. coli* ATCC 25922 was tracked spectrophotometrically for 13 h with the addition of **L5, L9** and **PDS** (at 1 x MIC) in the exponential phase (5 h). All three ligands exerted bacteriostatic activity, as evidenced by a plateau, rather than reduction, in absorbance after administration (**Figure S2**). To further investigate effects on cell physiology, *E. coli* treated with **L5, L9** and **PDS** (2 h at 1 x MIC) were visualized by scanning electron microscopy (SEM) (**Figure S3**). **L5** and **PDS** appeared bacteriolytic, as evidenced by visible cell debris and cell wall damage, while **L9** did not exhibit this effect.

Next-generation sequencing-based techniques have recently been applied to experimentally identify G4 motifs.^9, 10^ Marsico *et al*^*10*^ identified 47 observed G4 sequences (OQs) in the sense and antisense DNA strands of *E. coli* K-12 that were stabilized by addition of K^+^ (a known stabilizer of G4 structures); and 560 G4 OQs that were stabilized by the addition of K^+^ + PDS. Shao *et al*^*9*^ identified 168 mRNA sequences stabilized by K^+^. Given this large number of potential G4 ligand binding sites in the *E. coli* genome/transcriptome, we explored a proteomic approach to identify candidate G4 sequences as potential targets for, and to further investigate the mode of action of, **L5** and **L9**. We hypothesized that ligand-induced stabilization of G4 motifs would alter protein expression from genes containing these sequences (in either upstream regulatory or coding regions) compared to untreated controls.

*E. coli* ATCC 25922 cultures were individually treated with **L5, L9** and **PDS** (64, 1 and 64 µg/mL, respectively 7 h) (**Figure S4**), and their total cell proteomes, together with untreated controls, analyzed by tandem mass tagging (TMT)-labelled LC-MS/MS (**Figure S5A**). A “significance” ruleset was applied to the datasets where only statistically significant (p ≤ 0.05) differences of at least 2-fold in protein levels (-1 ≤ Log2FC ≥ 1) for treated vs. untreated cultures were considered. In all cases significant differences in expression levels were observed for large numbers of proteins (**Figure 2A**); these were classified in PANTHER^22^ and mapped in STRING^23, 24^ to compare protein-protein interactions (PPI) and enrichments. Despite their differing antibacterial activities, **L9** and **PDS** both upregulated proteins in similar categories (e.g. RNA metabolism, translation, metabolite interconversion) clustering in the biological process of translation (**Figure S6**), while **L5** upregulated proteins involved in transcriptional regulation, transport and metabolite interconversion. All three ligands downregulated gene-specific transcriptional regulators, transporters and metabolite interconversion proteins. The datasets were then searched for proteins associated with the OQs identified by Marsico *et al*^9, 10^, with the results showing that **L9** up-(32) or down-(46) regulated expression of more G4-associated proteins than either **L5** (25 and 43) or **PDS** (18 and 34) **(Figure 2B)**, consistent with the possibility of a G4-mediated mechanism of antibacterial action.

**Figure 2.**
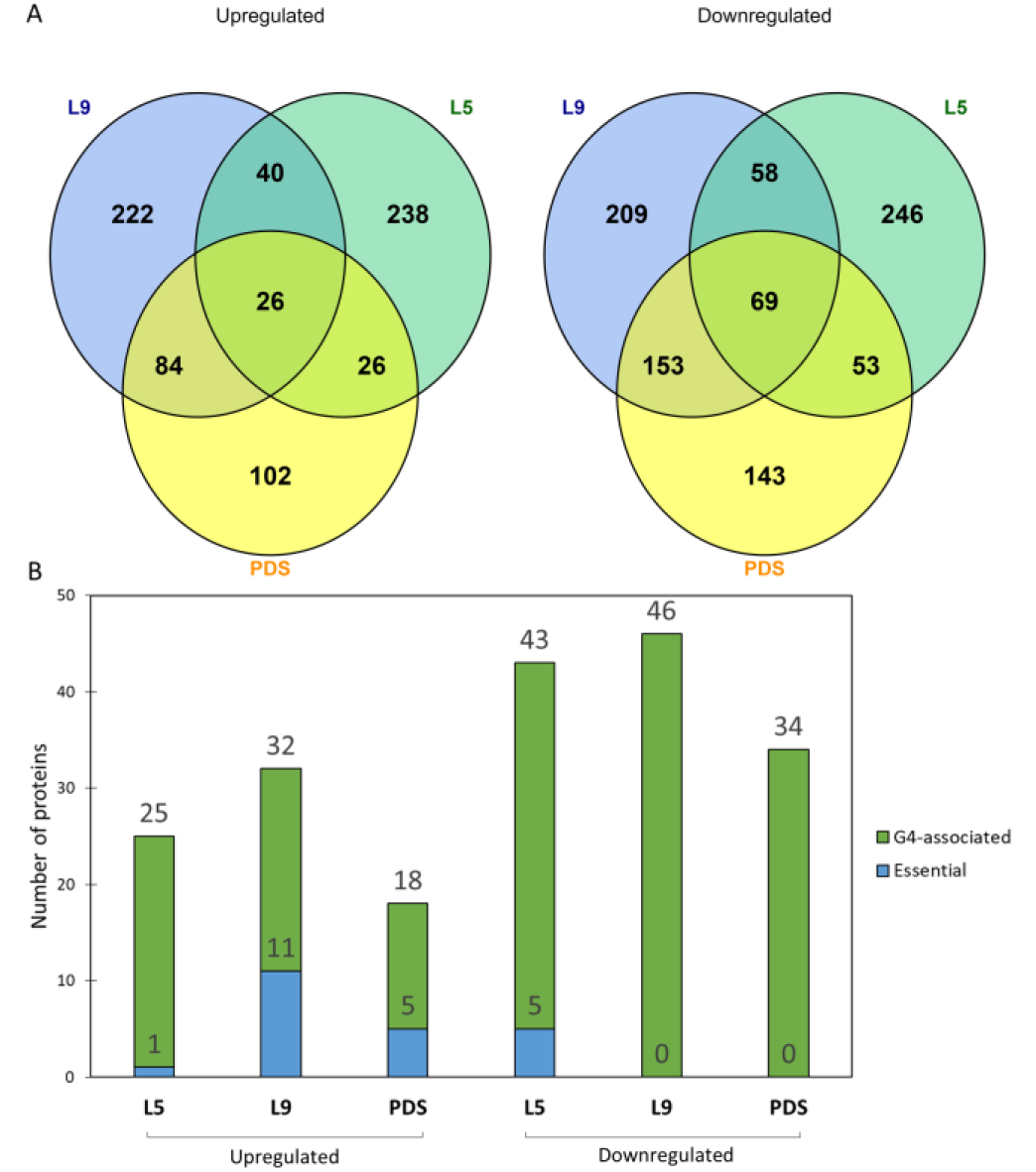
Effect of G-Quadruplex Ligands on *E. coli* Proteome. **(A)** Numbers of proteins with expression up/downregulated by **L5, L9** or **PDS**, compared to untreated controls. **(B)** Numbers of G4-associated (green) and essential G4-associated (blue) proteins with expression up/downregulated by L5, L9 or PDS, compared to untreated controls. (Quoted numbers are p ≤ 0.05, -1 ≤ Log2FC ≥ 1 in all cases).

Next, the overall effect of **L5, L9** and **PDS** exposure on expression of essential genes was examined, as this would be expected to influence *E. coli* viability. Previous reports observed G4 ligands to affect expression of essential genes in processes such as DNA repair, drug efflux etc., in various species (Mtb, *S. pneumoniae, K. pneumoniae*)^13, 14, 25^ Accordingly, the abundances of proteins encoded by 302 genes listed as essential for growth on the Profiling of *E. coli* Chromosome (PEC) database^31^ were identified in the proteomic datasets.^26^ Changes in levels of these proteins are summarized in **Figure S5B. L9** upregulated 69 proteins, compared to 16 and 57 for **L5** and **PDS**, respectively. In contrast, only 2 proteins were downregulated by **L9**, compared to 31 and 4 for **L5** and **PDS**. Thus, treatment of *E. coli* by candidate G4 ligands leads predominantly to upregulation of a subset of essential genes. For the 43 previously identified G4-containing genes^9, 10^ present in the PEC database (**Table S3**), 11 were upregulated by **L9**, compared to 1 and 5 in the **L5** and **PDS**-treated datasets respectively (**Figure 2B**). While the numbers of G4-associated essential genes regulated by **L5, L9** and **PDS** were small, these data nevertheless identify **L9** as more strongly affecting expression of G4-associated essential genes than either **L5** or **PDS**, consistent with its greater antibacterial activity.

Bioinformatic tools were next applied to the OQ-associated proteins of interest (**Figure 2B**), to shortlist candidate G4 sequences for biophysical investigation with **L5** and **L9**. All OQs were screened using G4Hunter^27^ to assess their propensity to form G4 structures *in vitro*, and seven were chosen for further investigation. Ten further OQs and putative G4 sequences (PQS) were chosen following *in silico* analysis of the complete genome of *E. coli* K-12 MG1655 using the R-package G4-iM Grinder^28^, selecting those with significant changes in protein expression in **L9**-treated cells, with the exception of *ybiO* (selected as a negative control for which little change in expression level was observed **(Table S4** and SI for details)).

All synthetic oligonucleotides corresponding to these 17 G4 DNA sequences formed G4 structures in solution^30^, as evidenced by circular dichroism (CD) spectroscopy (**Figure S7**), with some undergoing conformational transitions in the presence of K^+^. Preliminary screening using Fluorescence Resonance Energy Transfer (FRET) melting assays in the presence of 100 mM K^+^ and 5 µM ligand (**L5, L9** and **PDS, Table S7)** revealed that **L5** and **PDS** stabilized all of the sequences (ΔT_m_ values from 9.4 - 39 °C; whereas **L9** appears more selective towards 7 sequences (i.e. *glnD, pdxA, thrA, yjbD, aroL, cobS*, and *mtgA*) whose ΔT_m_ values range from 9.4-16.7 °C. Importantly, negligible duplex DNA (F10T) stabilization was observed for either **L5** (ΔT_m_ = 3.0 °C at 5 µM) or **L9** (ΔT_m_= 0.4 °C at 5 µM), further confirming these to be selective G4 ligands as opposed to non-specific DNA intercalators.

To further investigate binding of **L5** and **L9** to the six most promising bacterial G4 sequences, UV-Vis titrations were performed^31^ (**Figures 3, S8**). All ligands bind these sequences with similar affinities (*K*_D_ = 0.34 - 0.61 μM). G4 binding was further confirmed by CD spectroscopy titrations (**Figure S9**) which showed the three ligands significantly perturb the CD spectra of all sequences. An attenuation of ellipticity between 240 - 280 nm, with an increase at 290 nm, was observed for the parallel *pdxA, thrA* and hybrid *yjbD* sequences, while an increase towards a more parallel topology was shown for the mixed hybrid-parallel *cobS* and parallel *aroL* sequences and a shift towards hybrid for *mtgA*. Both **L5** and **L9** can bind G4 sequences *in vitro*, although stabilization of a wider range of G4 sequences (**Table S6**) does not appear to correlate with anti-bacterial activity; with **L9** exhibiting 16-fold better activity against *E. coli* (MIC 4 µg/mL) compared to **L5** and **PDS** (MIC 64 µg/mL in each case).

**Figure 3.**
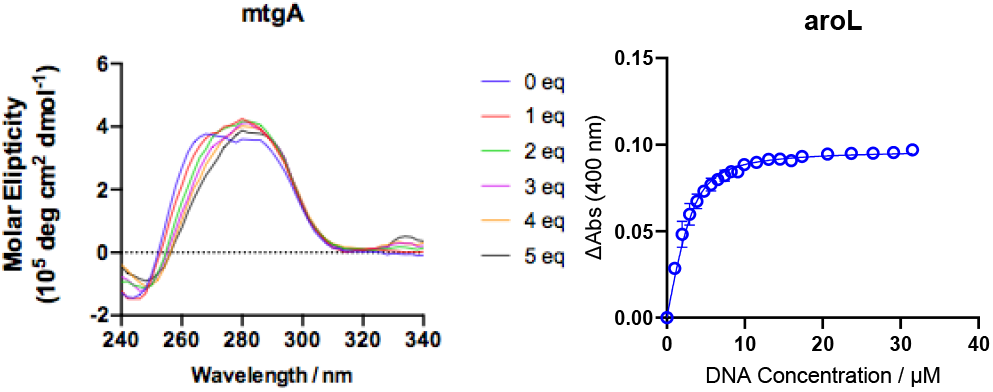
L9 binds G-quadruplex sequences *in vitro*. (**Left**) Representative CD spectra of parallel *mtgA* oligonucleotid G4 structure with varying concentrations of **L9**. (**Right**) UV-Vis titration of **L9** into *aroL* G4 DNA at 100 mM K^+^sequence monitored by absorbance spectroscopy. Fitted line is to one-site binding equation giving Ka = 0.52 mM (N=3).

We then employed the fluorescence microscopy-based Bacterial Cytological Profiling (BCP) technique, as previously applied to *E. coli*, ^32, 33^ to further investigate the mechanism of action of **L9**. The morphological changes induced by **L9** were compared with those of antibiotics targeting cellular pathways that interfere with the flow of genetic information, including a DNA replication inhibitor (mitomycin C; MMC), an RNA transcription inhibitor (rifampicin; RIF), and a protein translation inhibitor (chloramphenicol; CAM), together with PDS as a known G4 ligand control. BCP analysis (**Figure4**) showed that the effects of **L9** treatment differ from those induced by the other three antibiotics, with DNA (blue) appearing condensed in **L9**-treated cells while punctate staining was evident in the membrane (red, **Figure 4F**). Image analysis profiles of **L9**-treated cells clustered separately from those of both untreated controls and cells exposed to comparator antibiotics (**Figure 4G**). In contrast, PDS-treated cells clustered closer to those exposed to chloramphenicol. Similar to previous studies where distinct profiles were observed,^32^ it is possible that **L9** inhibits pathways distinct from those targeted by the comparator antibiotics. Interestingly, comparison of the **L9** profile with data from the original *E. coli* BCP study, which described 18 distinct morphological changes, did not reveal resemblance to any previously reported profiles. ^32^ It is also possible that **L9** exhibits simultaneous inhibitory activity against multiple cellular pathways, resulting in a combinatorial effect on bacterial morphology, as observed previously when bacteria were treated with two antibiotics. ^34, 35^

**Figure 4.**
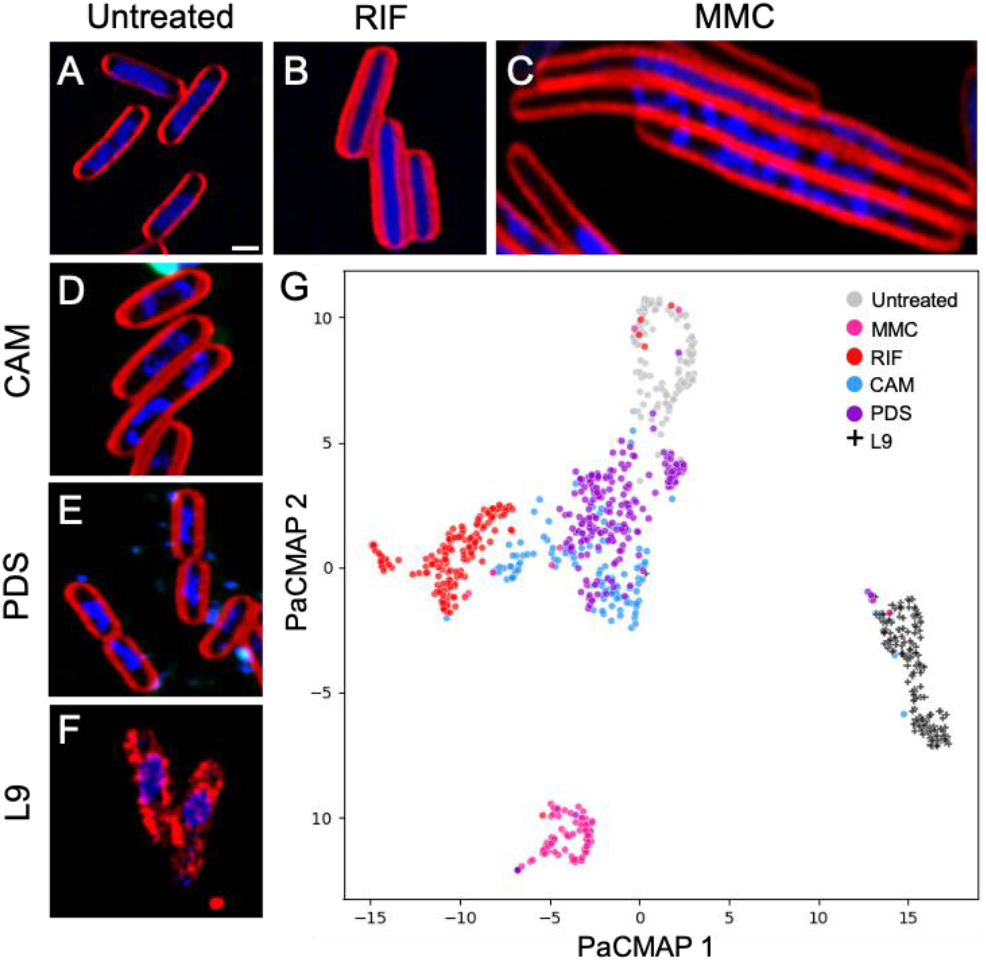
L9-treated *E. coli* Exhibit Distinct Morphological Changes. (**A – F)** Representative laser confocal images (blue, DAPI; red, FM-4-64) of *E. coli* treated with comparator antibiotics rifampicin (RIF), mitomycin C (MMC) and chloramphenicol (CHL, **B - D**); plus PDS (**E**), untreated control (**A**) and **L9** (**F**). (**G)** Representative Pairwise Controlled Manifold Approximation (PaCMAP) analysis of morphological parameters. Note that **L9**-treated cells (points represent individual cells) form a cluster distinct from those generated by treatment with comparator agents.

The data presented here identify the azobenzene G4 ligand **L9** as possessing bacteriostatic antibacterial activity against reference and clinical multi-drug resistant *E. coli*. Electron and laser confocal microscopy, the latter supported by BCP analysis, show **L9** treatment to induce a phenotype differing from those conferred by exposure to comparator antibiotics or G4 ligands, identifying **L9** as an anti-Gram-negative agent with a distinct mode of action. Proteomic analysis reveals **L9** exposure to induce expression changes in multiple proteins, both G4-associated and non-associated. When considered alongside our hypothesis that the effects of **L9** upon bacterial cells arise from interaction with G4 sequences, these data implicate bacterial G4 sequences as potentially involved in a wide range of cellular processes, including translation. Compared to the G4 ligands **L5** (stiff-stilbene) and **PDS**, the response to **L9** differs in that expression levels of greater numbers of G4-associated, and G4-associated and essential, proteins are affected. This is striking in the context of our biophysical data showing **L9** to be more selective with respect to G4-containing oligonucleotides than either comparator. Importantly, therefore, antibacterial activity is then not solely a result of a high affinity of **L9** for G4 sequences, but likely arises from interactions with specific sequences or sequence types in the (bacterial) cellular environment, suggesting scope for optimizing antibacterial activity of G4 ligands over that towards other cell types. The selectivity of **L9** for *E. coli* over other Gram-negative species, that extends to multi-resistant *E. coli* strains and persists even in the presence of permeabilizing agents, is then consistent with targeting of a specific subset of cellular G4 targets, some of which may be associated with essential proteins. The possibility of multi-targeting, i.e. that **L9** binding to multiple G4 sequences contributes to overall activity, is a further attractive aspect of G4-associated antibacterials as accumulation of changes at multiple sites would then be required to generate mutational resistance.

Taken together, these data support bacterial G4 structures as viable targets against which to pursue antibacterials development, with potential for developing both anti-Gram-negative and species-specific activity. Our findings justify exploration of small-molecule G4 ligands as scaffolds for urgently-needed antibacterials against some of the most important healthcare-associated pathogens.

## ASSOCIATED CONTENT

### Supporting Information

Synthetic procedures and spectroscopic data, supporting figures and tables, and full experimental details are available in the supporting information.

## AUTHOR INFORMATION

### Author Contributions

The manuscript was written through contributions of all authors. All authors have given approval to the final version of the manuscript.

## ACKNOWLEDGMENTS

This research was supported by ERC-COG: 648239 (MCG), EPSRC EP/G036764/1 (JSamphire) and EPSRC EP/S026215/1 (MCG, JS). EP/R043361/1 (TS/R014329/1) (MCG, JS) and GCRF EP/T020288/1 (MCG, JS). EPSRC EP/L015366/1 (MPO) and the European Research Council (ERC-COG: 648239) and J. R.-S. thanks MSCA fellowship (project 843720-BioNanoProbes). E.B.-R. was supported by a Maria Zambrano fellowship, financed by NextGenerationEU/European Union. Data processing and analysis for BCP was funded by National Research Council of Thailand (NRCT) and Mahidol University: N42A650368. We also thank Nina Allen for HPLC analysis assistance and Dominic Alibhai of the Wolfson Bioimaging Facility for their help with the confocal microscopy work.

## ABBREVIATIONS

TMPyP4: 5,10,15,20-Tetrakis-(N-methyl-4-pyridyl)porphine
PDS: pyridostatin
PMBN: polymyxin B nonapeptide
TMT: tandem mass tag
FRET: Fluorescence resonance energy transfer
CD: Circular Dichroism

