## Supplementary Data for "Small molecule G-quadruplex ligands are antibacterial candidates for Gram-negative bacteria"

**Supplementary Information Contents**

1. **Synthesis and characterization for L1-L11**
2. **Antimicrobial screening**
3. **Proteomic analysis of *E. coli* treated with L5, L9 and PDS**
4. **Bioinformatic analysis of OQ-associated proteins of interest to shortlist candidate G4 sequences for biophysical investigation**
5. **Circular Dichroism (CD) analysis of G4 DNA candidate sequences identified by bioinformatics**
6. **FRET melting assays**
7. **UV-visible spectroscopy**
8. **Circular Dichroism Titrations of L5, L9 and pyridostatin to *pdxA, thrA, yjbD, aroL, cobS*, and *mtgA* oligonucleotide sequences**
9. **Bacterial Cytological Profiling (BCP) image acquisition and analysis**
10. **References**

### Materials and Methods

#### 1. Synthesis and characterization for L1-L11

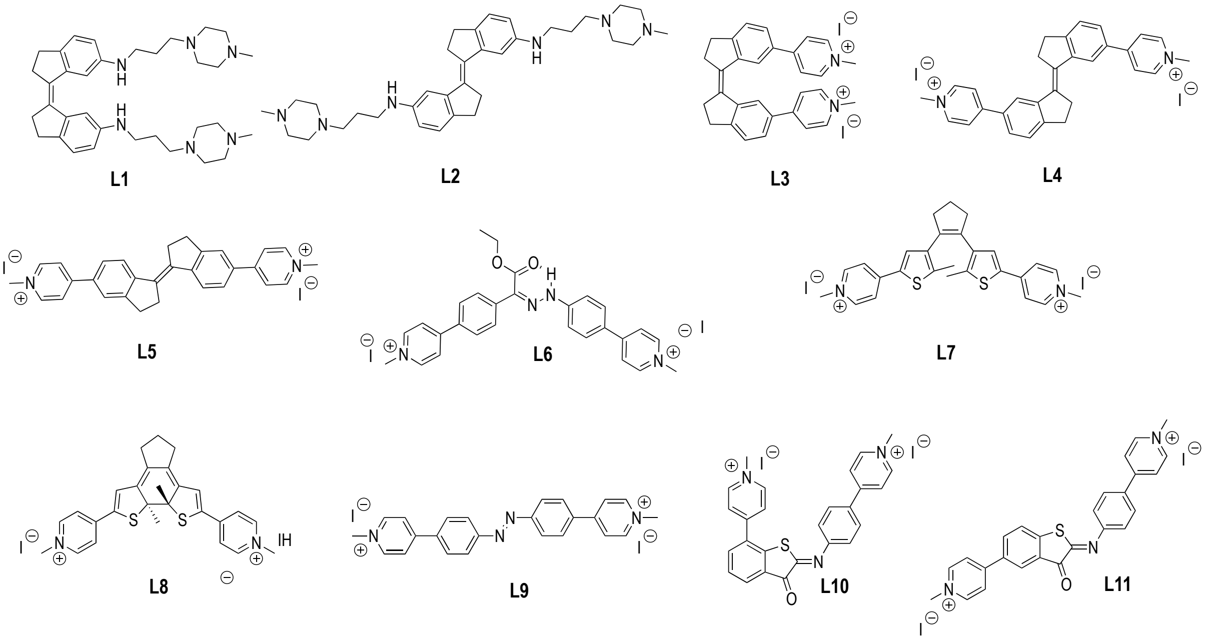

**Figure S1.** List of Ligand library selected for this study.

**General.** Reagents and solvents were purchased as reagent grade and used without further purification. Reactions requiring anhydrous conditions were performed under $N_2$; glassware and needles were either flame dried immediately prior to use, or placed in an oven (150 °C) for at least 2 h and allowed to cool in a desiccator or under reduced pressure. Compounds **L1-L4**[1, 2], **L5**[3] and DTE ligands **L7-L8**[4] were prepared following previous reported procedures. For column chromatography, silica gel 60 (230-400 mesh, 0.040-0.063 mm) was purchased from E. Merck. Thin Layer Chromatography (TLC) was performed on aluminium sheets coated with silica gel 60 $F_{254}$ purchased from E. Merck, visualization by UV light. NMR spectra were recorded on a Bruker AC 400 or AC500 with solvent peaks as reference. $^1H$ and $^{13}C$ NMR spectra were obtained for solutions in $CDCl_3$ and DMSO-$d_6$. All the assignments were confirmed by one- and two-dimensional NMR experiments (DEPT, COSY, HSQC and HMBC). Mass spectra were obtained by the University of Bristol mass spectrometry service by electrospray ionisation (ESI).

#### Synthesis of L6

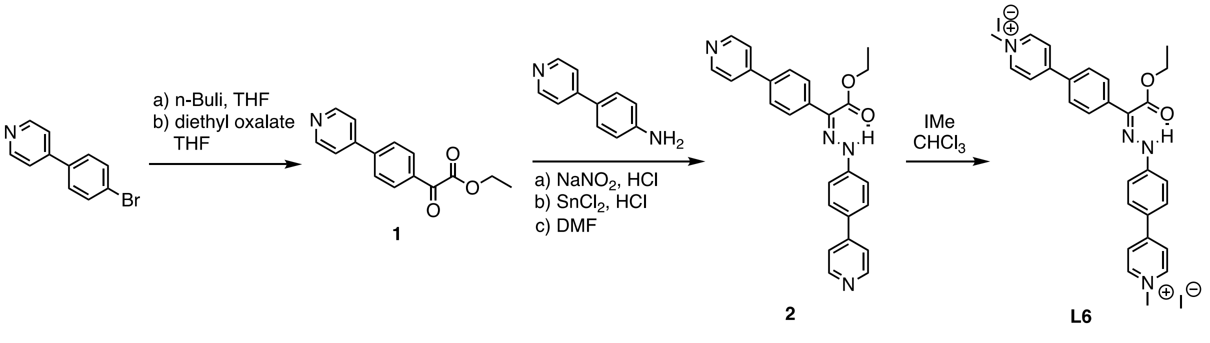


#### Ethyl 2-oxo-2-(4-(pyridin-4-yl)phenyl)acetate (1)

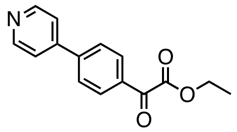

To a solution of 4-(4-bromophenyl)pyridine (500 mg, 2.04 mmol) in dry THF (4 mL), under atmosphere of $N_2$ and cooled to -78 °C, *n*-BuLi (2.5 M in hexane, 896 µL, 2.24 mmol) was added dropwise over 15 min. The reaction mixture was stirred at -78 °C for an additional 15 min. The resulting suspension was then transferred dropwise via canula at -78 °C to a separate round-bottom flask containing a solution of diethyl oxalate (339 µL, 3.26 mmol) in dry THF (3 mL) at -78 °C. The reaction mixture was stirred at -78 °C for 20 min and the allowed to warm to 0 °C. Water (8 mL) was slowly added to quench the reaction. The mixture was then extracted with $Et_2O$ (3 times). The combined organic layers were dried over anh. $MgSO_4$, filtered and concentrated. The crude was purified by silica gel chromatography column (EtOAc/hexane, 3:2) to give compound **1** (172 mg, 33%) as a pale yellow solid. $^1$H NMR (400 MHz, $CDCl_3$) $\delta$ 8.73 (d, *J* = 6.2 Hz, 2H), 8.15 (d, *J* = 8.8 Hz, 2H), 7.77 (d, *J* = 8.8 Hz, 2H), 7.5 3 (d, *J* = 6.1 Hz, 2H), 4.48 (q, *J* = 7.2 Hz, 2H), 1.45 (t, *J* = 7.1 Hz, 3H); $^{13}$C NMR (101 MHz, $CDCl_3$) $\delta$ 185.6, 163.5, 150.6, 146.7, 144.3, 132.7, 130.9, 127.6, 121.7, 62.5, 14.1; ESI-HRMS *m/z* calcd. for $C_{26}H_{23}N_4O_2$ [M+H]$^+$: 256.0968; found: 256.0979.

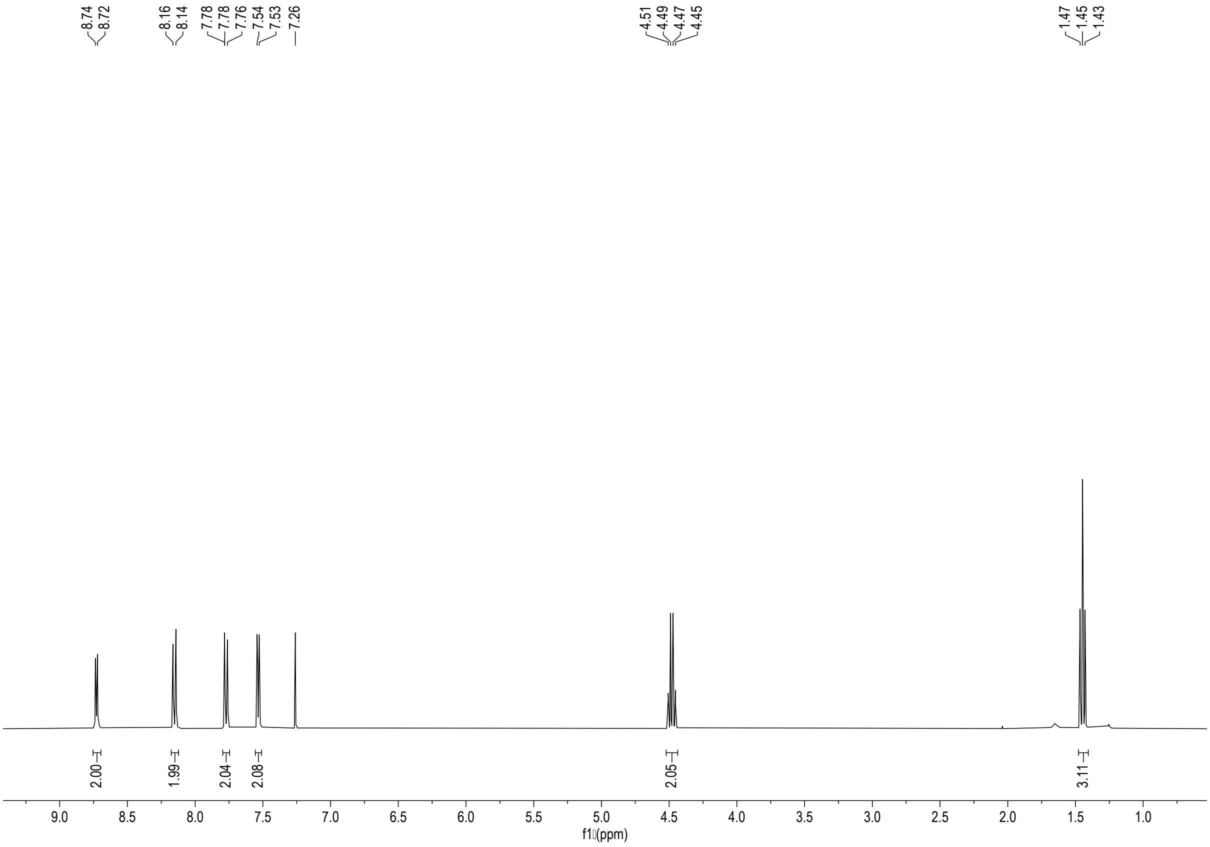


$^1$H-NMR spectrum of **1** (400 MHz, $CDCl_3$)

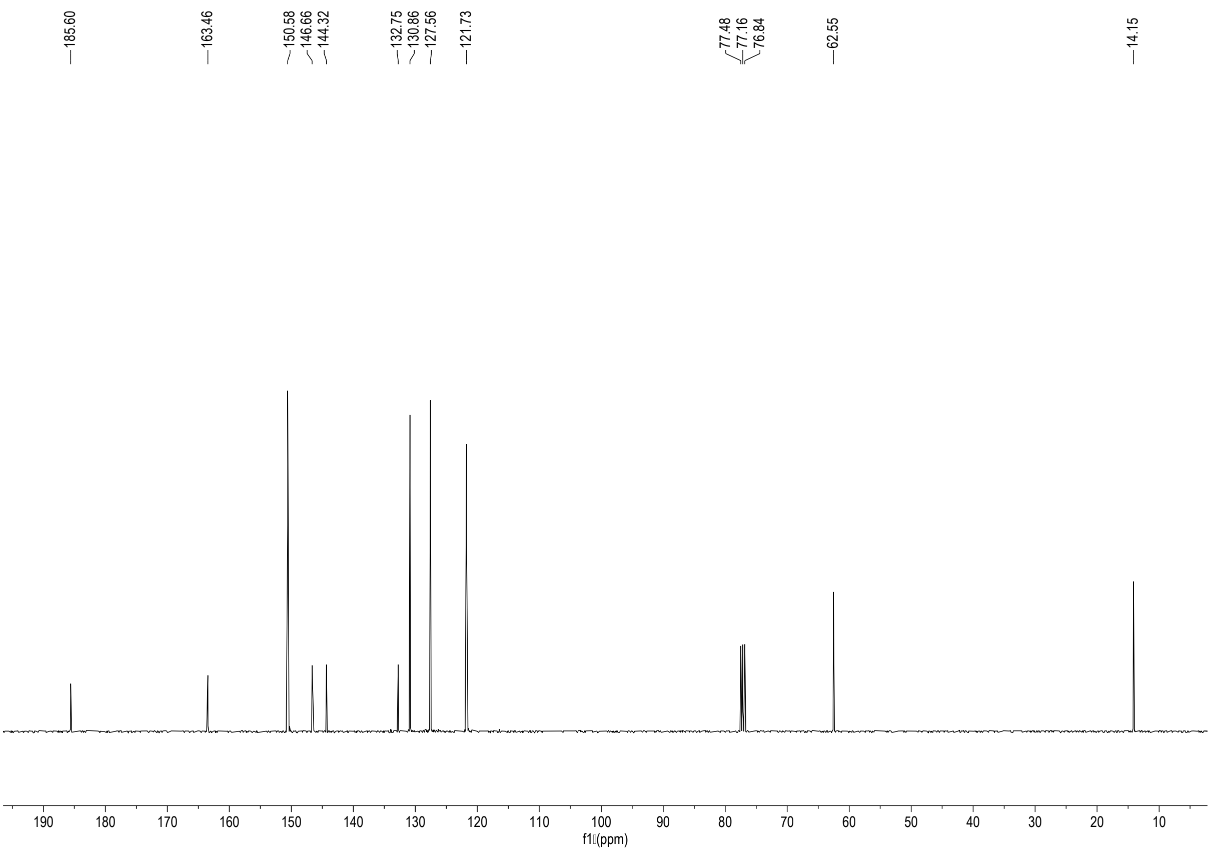


$^{13}$C-NMR spectrum of **1** (101 MHz, $CDCl_3$)

**Compound 2.**

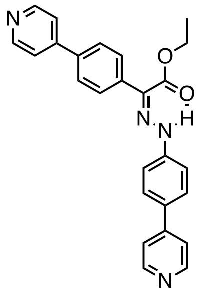

A pre-cooled solution of $NaNO_2$ (46 mg, 0.67 mmol) in water (0.25 mL) was added to a solution of 4-(pyridin-4-yl)aniline (117 mg, 0.67 mmol) in 20% HCl (1.5 mL) over a period of 30 min, followed by 1 h stirring at 0 ºC. Then, a solution of $SnCl_2$ (256 mg, 1.33 mmol) in 37% HCl (1 mL) was added dropwise to the solution. The reaction mixture was warmed up and left to stir at r.t. for 1 h. A solution of compound **1** (170 mg, 0.67 mmol) in DMF (2 mL) was then added to the obtained hydrazine. After 1 h heating at 80 °C, the reaction mixture was cooled to r.t., followed by neutralization using sat. $NaHCO_3$ solution. The suspension mixture was diluted with water, and extracted with DCM. The organic layer was further washed with sat. $NaHCO_3$ and brine and then dried over anh. $MgSO_4$, filtered and concentrated. The crude was purified by silica gel chromatography column (EtOAc/DCM, 10:1) to give compound **2** (188 mg, 67%) as a yellow solid. $^1$H NMR (400 MHz, $CDCl_3$) $\delta$ 12.59 (s, 1H), 8.68 (d, $J$ = 6.1 Hz, 2H), 8.63 (d, $J$ = 6.2 Hz, 2H), 7.83 (d, $J$ = 8.4 Hz, 2H), 7.67 (dd, $J$ = 11.9, 8.6 Hz, 4H), 7.56 (d, $J$ = 6.2 Hz, 2H), 7.50 (d, $J$ = 6.2 Hz, 2H), 7.41 (d, $J$ = 8.7 Hz, 2H), 4.42 (q, $J$ = 7.1 Hz, 2H), 1.42 (t, $J$ = 7.2 Hz, 3H); $^{13}$C NMR (101 MHz, $CDCl_3$) $\delta$ 163.6, 150.6, 150.4, 150.3, 147.9, 147.7, 144.0, 137.3, 137.2, 132.1, 130.0, 129.4, 128.4, 128.2, 128.1, 128.0, 126.6, 121.7, 121.5, 121.0, 121.0, 115.0, 114.8, 61.5, 14.3; ESI-HRMS *m/z* calcd. for $C_{26}H_{23}N_4O_2$ $[M+H]^+$: 423.1816; found: 423.1824.

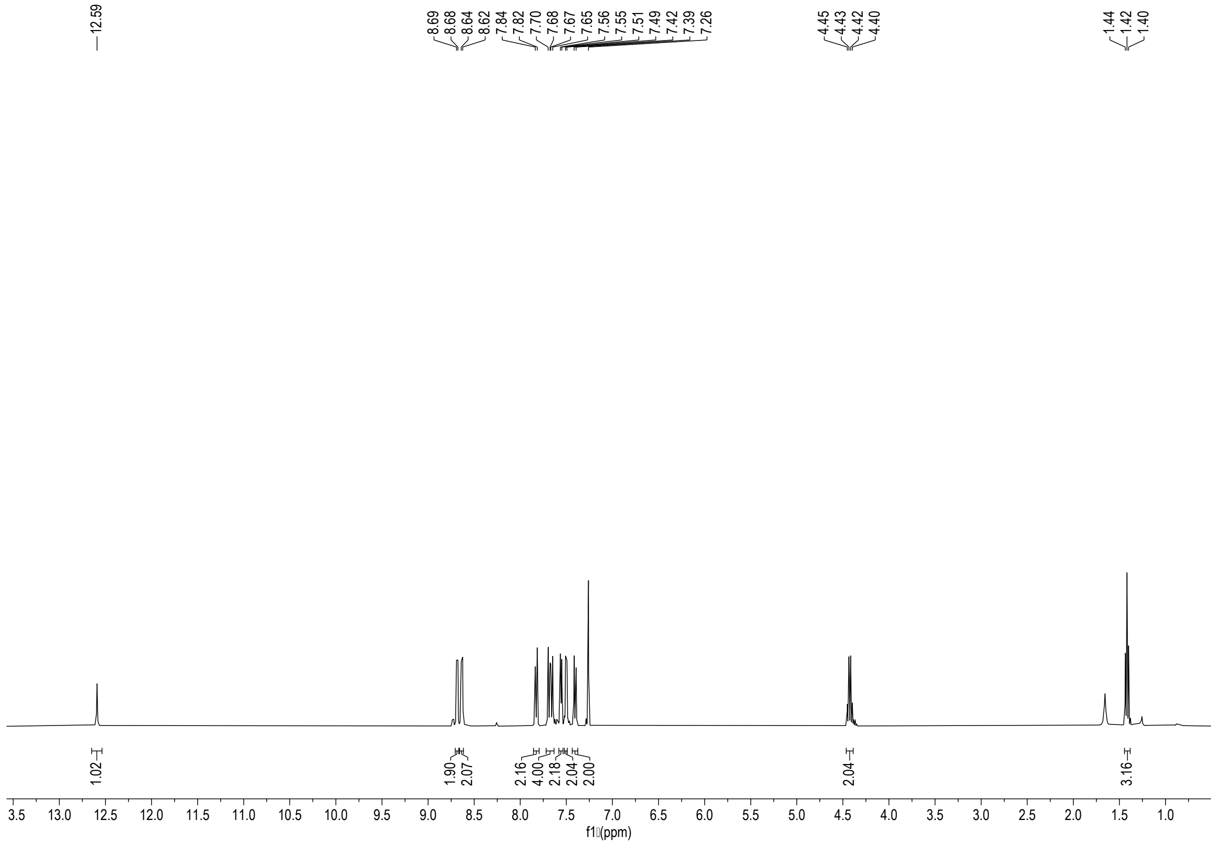


$^1$H-NMR spectrum of **2** (400 MHz, $CDCl_3$)

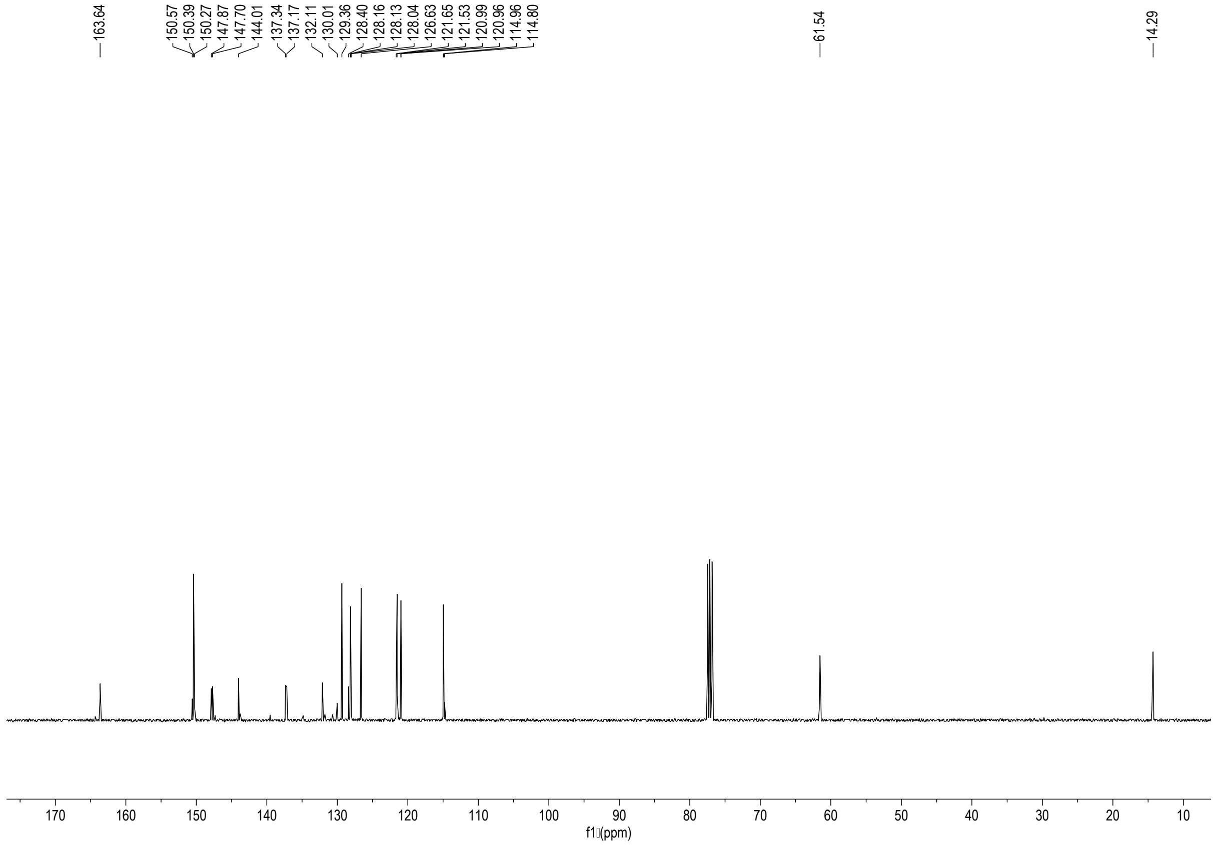


$^{13}$C-NMR spectrum of **2** (101 MHz, $CDCl_3$)

**Compound L6**

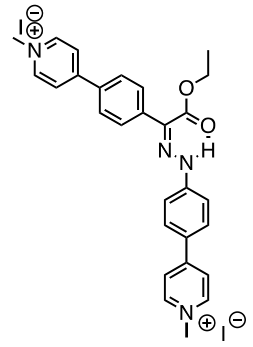

To a solution of compound **2** (90 mg, 0.21 mmol) in $CHCl_3$ (5 mL), methyl iodide (53 µL, 0.85 mmol) was added. The solution was stirred in a sealed vessel at 50 °C overnight. The generated solid was filtered and washed with diethyl ether. Following filtration, compound **L6** (144 mg, 96%) was obtained as a yellow powder. $^1$H NMR (400 MHz, DMSO-$d_6$) $\delta$ 11.93 (s, 1H), 9.02 (d, $J$ = 6.8 Hz, 2H), 8.89 (d, $J$ = 6.8 Hz, 2H), 8.54 (d, $J$ = 7.0 Hz, 2H), 8.44 (d, $J$ = 7.1 Hz, 2H), 8.15 (d, $J$ = 8.7 Hz, 4H), 7.95 (d, $J$ = 8.7 Hz, 2H), 7.63 (d, $J$ = 8.9 Hz, 2H), 4.45 (q, $J$ = 7.1 Hz, 2H), 4.34 (s, 3H), 4.28 (s, 3H), 1.35 (t, $J$ = 7.1 Hz, 3H); $^{13}$C NMR (101 MHz, DMSO-$d_6$) $\delta$ 162.3, 153.4, 153.3, 146.7, 145.5, 145.1, 138.5, 132.8, 131.3, 129.6, 128.5, 128.0, 126.1, 123.9, 122.4, 114.9, 61.8, 47.1, 46.7, 13.9; ESI-HRMS $m/z$ calcd. for $C_{28}H_{28}N_4O_2I$ $[M]^+$: 579.1257; found: 579.1251.

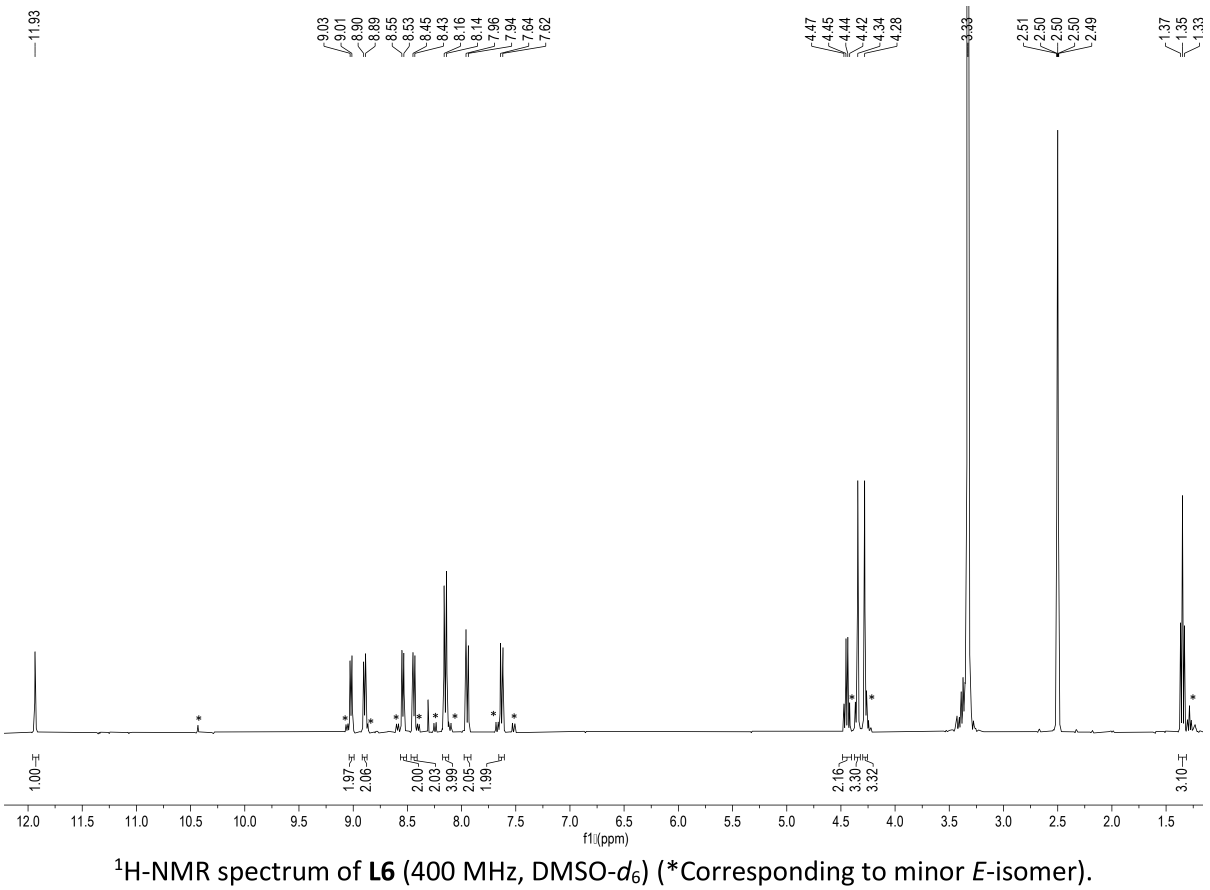


$^1$H-NMR spectrum of **L6** (400 MHz, DMSO-$d_6$) (*Corresponding to minor *E*-isomer).

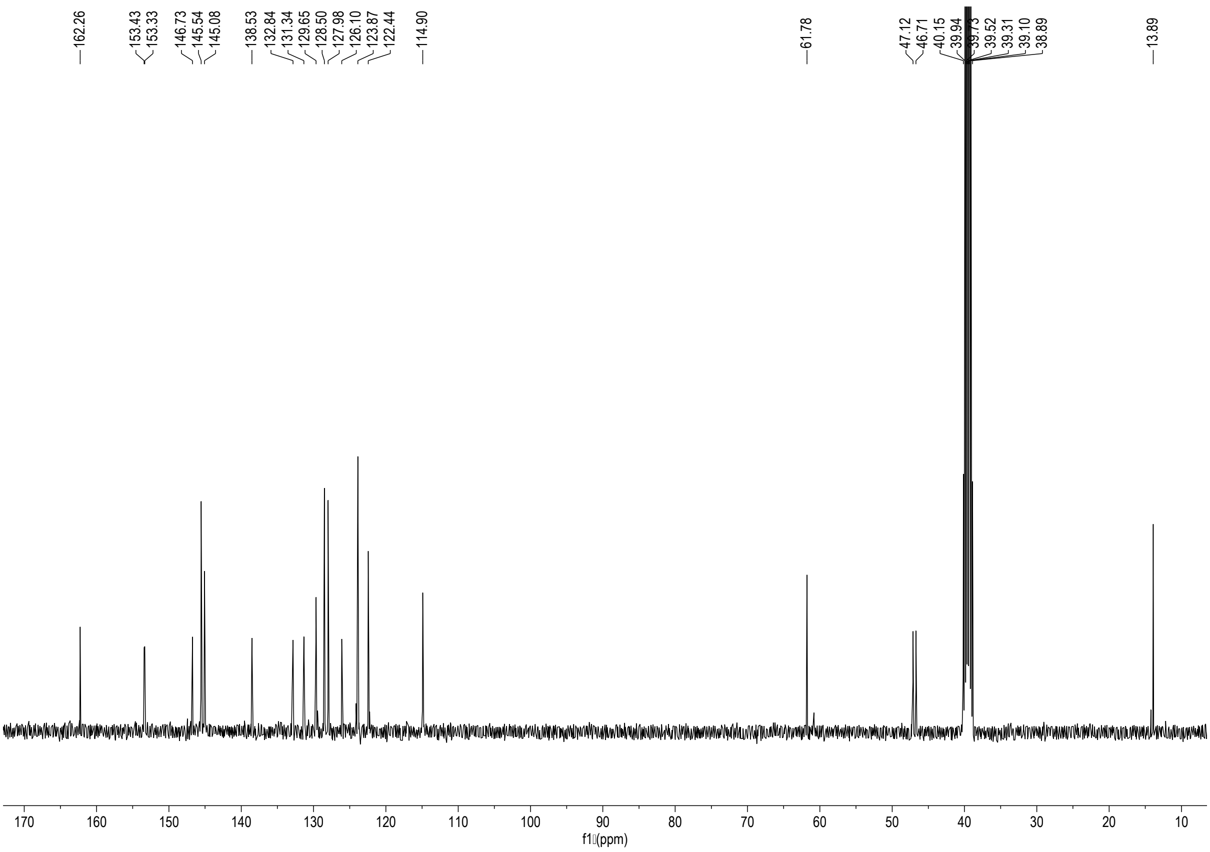

$^{13}$C-NMR spectrum of **L6** (101 MHz, DMSO-$d_6$)

**Synthesis of L9**

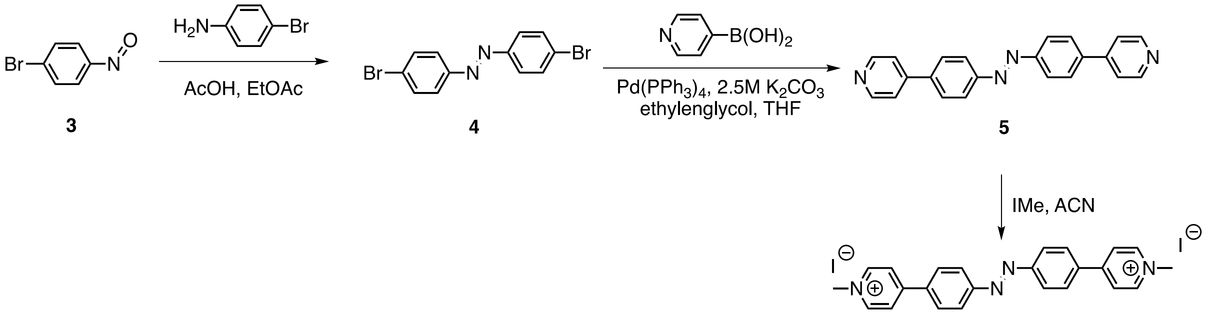

**(*E*)-4,4'-dibromoazobenzene (4)**

Nitrosobenzene derivative **3**[5] (1 g, 5.41 mmol) and 4-bromoaniline (960 mg, 5.41 mmol) were dissolved in AcOH/EtOAc (1:1, 20 mL). The reaction mixture was stirred at 45 ºC overnight. The mixture was cooled down to r.t., diluted with DCM and washed with water (x2). The combined organic layers were dried over anh. $MgSO_4$, filtered and concentrated. The crude was purified by silica gel chromatography column (hexane/DCM 9:1), to give compound **4** (1.57g, 86%) as an orange solid. Experimental data in agreement with reported literature.[6]

**(*E*)-4,4'-bis(pyridin-4-yl)azobenzene (5)**

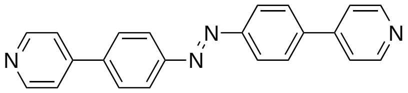

A suspension of (*E*)-4,4'-dibromoazobenzene (**4**) (300 mg, 0.89 mmol), $Pd(PPh_3)_4$ (104 mg, 0.09 mmol), ethyleneglycol (1 drop) and 4-pyridinylboronic acid (364 mg, 2.66 mmol) in a mixture of THF (12 mL) and aq. 2.5 M $K_2CO_3$ (3 mL) was bubbled with $N_2$ for 10 min. The resulting solution was heated at 70 °C overnight. After cooling to room temperature, water was added. The aqueous layer was extracted with DCM (x2) and the combined organic extractions dried over $MgSO_4$, filtered and concentrated *in vacuo*. The residue was purified by flash silica chromatography (DCM/MeOH, 40:1 $\rightarrow$ 30:1), afforded compound **5** (259 mg, 87%) as an orange amorphous solid. $^1H$ NMR (400 MHz, $CDCl_3$) $\delta$ 8.72 (d, $J$ = 6.2 Hz, 4H), 8.08 (d, $J$ = 8.6 Hz, 4H), 7.82 (d, $J$ = 8.5 Hz, 4H), 7.58 (d, $J$ = 6.2 Hz, 4H); $^{13}C$ NMR (101 MHz, $CDCl_3$) $\delta$ 152.9, 150.6, 147.3, 140.9, 128.0, 123.9, 121.7; ESI-HRMS $m/z$ calcd. for $C_{22}H_{17}N_4$ $[M+H]^+$: 337.1448; found: 337.1455.

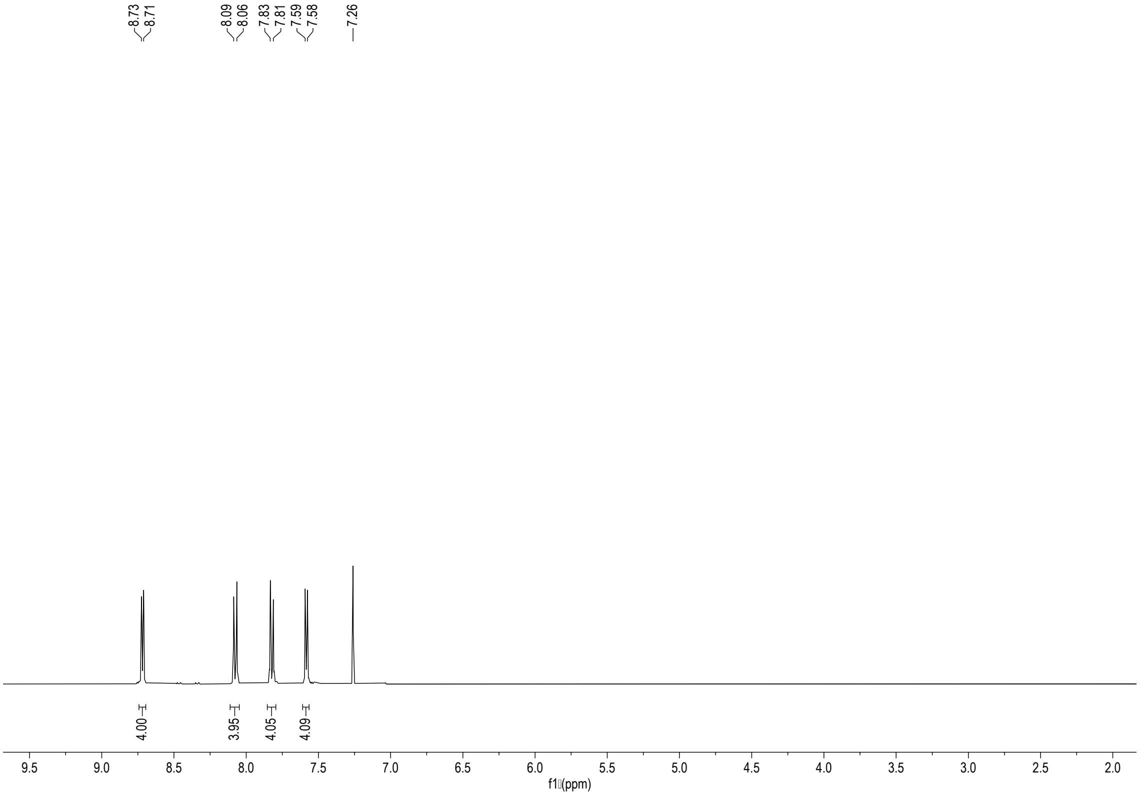


$^1H$ NMR spectrum of compound **5** ($CDCl_3$, 400 MHz).

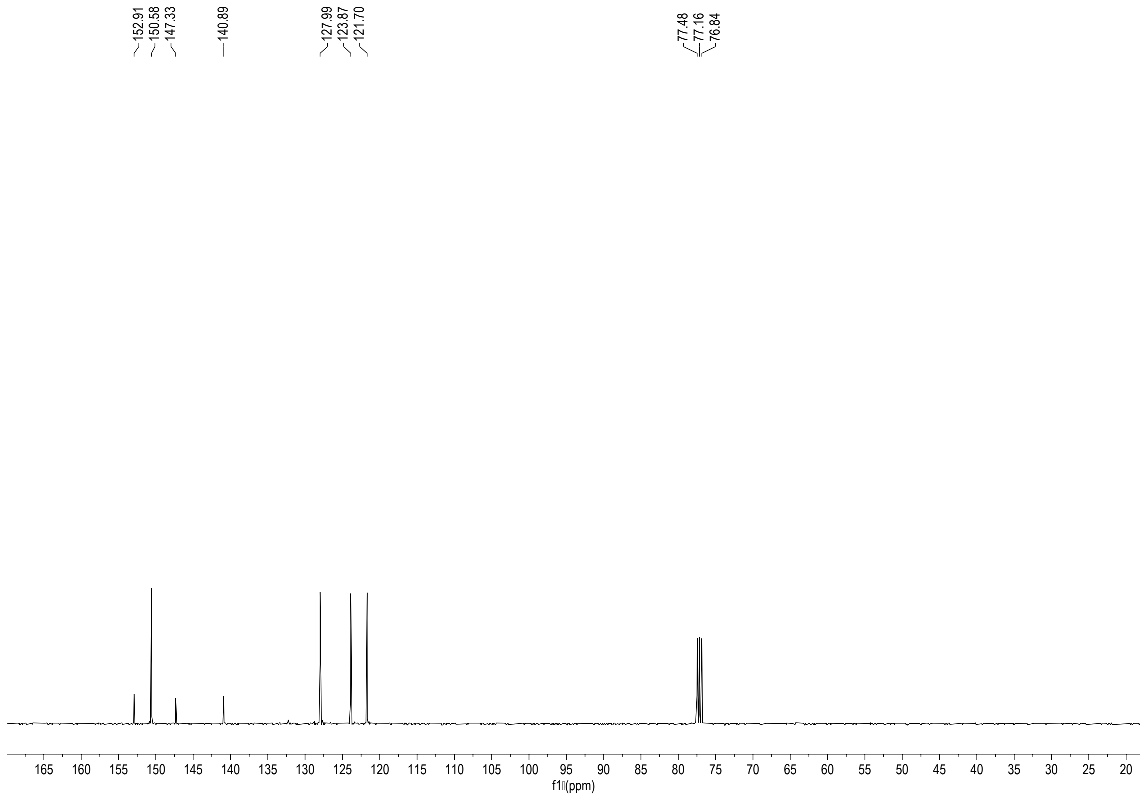


$^{13}C$ NMR spectrum of compound **5** ($CDCl_3$, 101 MHz).

**Compound L9.**

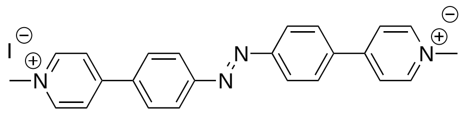

To a solution of compound **5** (100 mg, 0.30 mmol) in acetonitrile (8 mL), methyl iodide (148 µL, 2.38 mmol) was added. The solution was stirred in a sealed vessel at 50 °C overnight. The generated solid was filtered and washed with ether. Following filtration, compound **L9** (156 mg, 85%) was obtained as an orange powder. $^{1}H$ NMR (400 MHz, DMSO-$d_6$) $\delta$ 9.10 (d, $J$ = 6.5 Hz, 4H), 8.63 (d, $J$ = 6.9 Hz, 4H), 8.36 (d, $J$ = 8.5 Hz, 4H), 8.17 (d, $J$ = 8.6 Hz, 4H), 4.38 (s, 6H); $^{13}C$ NMR (101 MHz, DMSO- $d_6$) $\delta$ 153.4, 152.9, 145.8, 136.5, 129.6, 124.5, 123.8, 47.3; ESI-HRMS $m/z$ calcd. for $C_{24}H_{22}IN_4^+$ $[M]^+$: 493.0884; found: 493.0887.

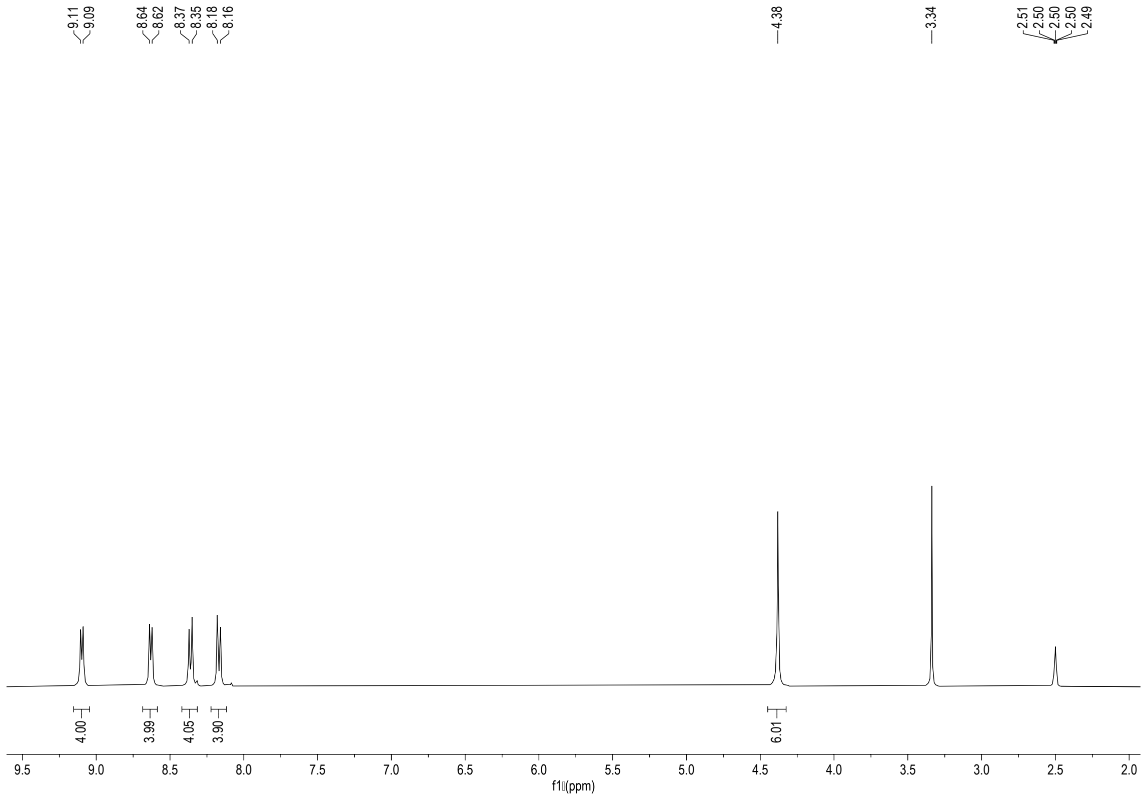


[1]H NMR spectrum of compound **L9** (DMSO-$d_6$, 400 MHz).

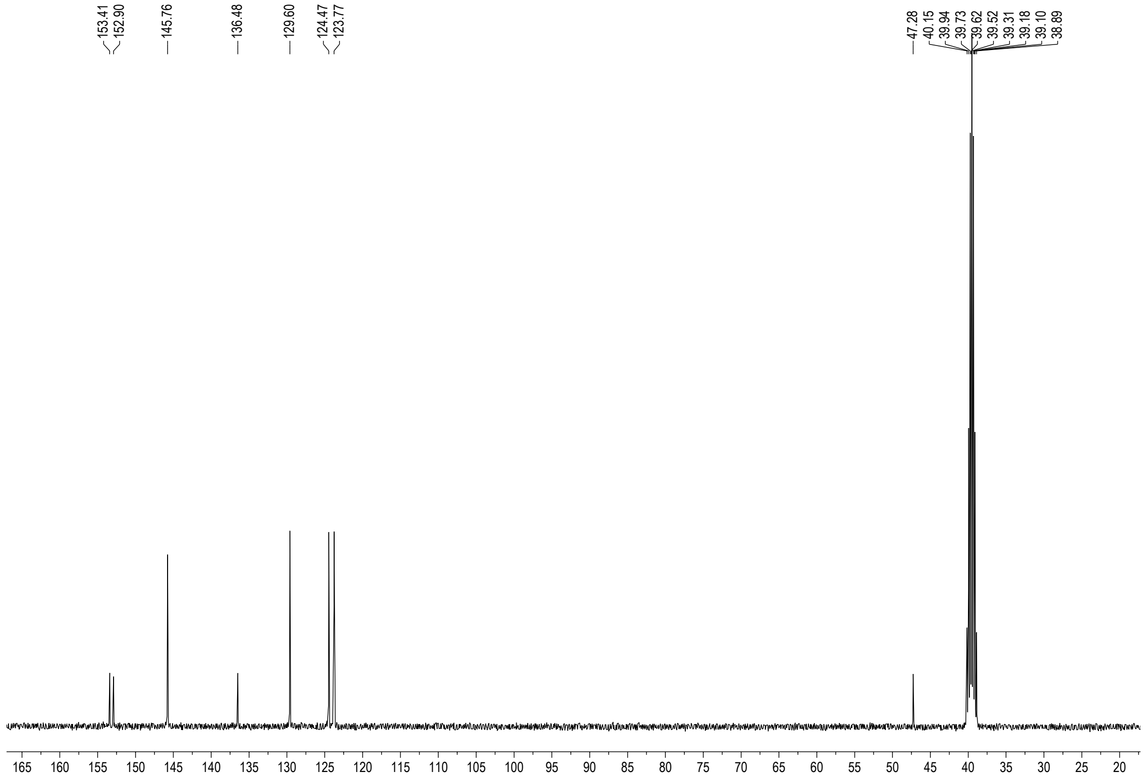


[13]C NMR spectrum of compound **L9** (DMSO-$d_6$, 101 MHz).

**Synthesis of L10.**

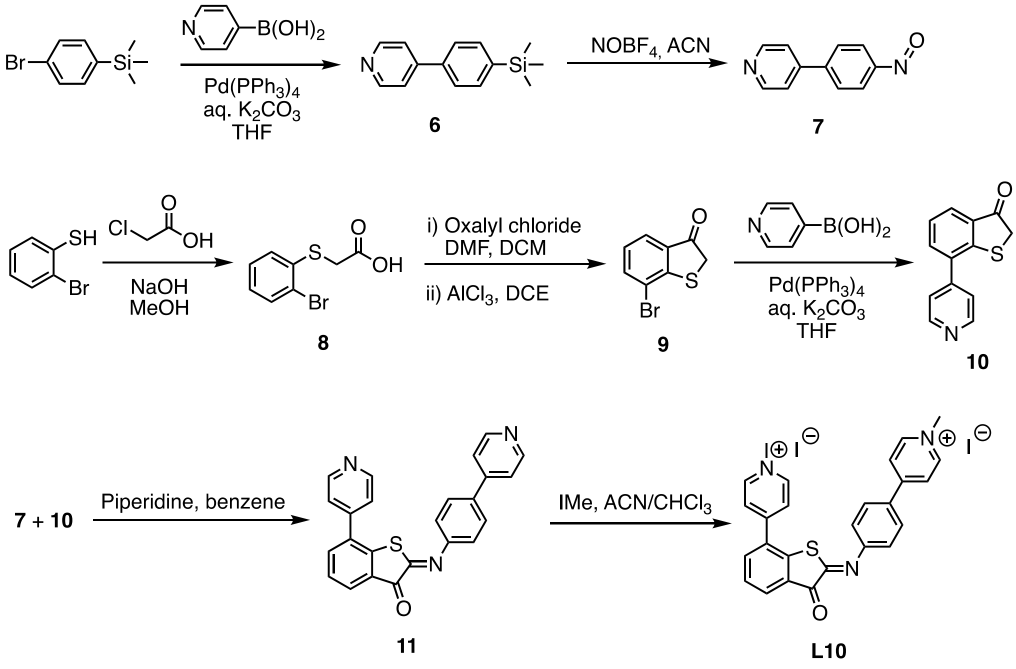


**4-(4-(trimethylsilyl)phenyl)pyridine (6)**

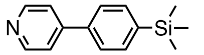

4-pyridinylboronic acid (1.3 g, 9.52 mmol), 1-bromo-4-(trimethyl)benzene (1.5 g, 6.35 mmol), $Pd(PPh_3)_4$ (378 mg, 0.32 mmol) and $K_3PO_4{\cdot}H_2O$ (9.2 g, 38.09 mmol) were dissolved in a mixture of degassed THF/$H_2O$ (10:1, 66 mL). The reaction mixture was heated to 60 °C and stirred overnight. The mixture was cooled and water was added. The aqueous layer was extracted with EtOAc and DCM. The combined organic layers were dried over anh. $MgSO_4$, filtered and concentrated. The crude was purified by silica gel chromatography column ($CHCl_3$/EtOAc, 100:0 ⟶ 10:1) to give compound **6** (1.42 g, 98%) as a yellow oil. $^1H$ NMR (400 MHz, $CDCl_3$) $\delta$ 8.66 (d, $J$ = 6.2 Hz, 2H, H-2), 7.68 – 7.59 (m, 4H, H-6, H-7), 7.51 (d, $J$ = 6.2 Hz, 2H, H-3), 0.31 (s, 9H, H-9); $^{13}C$ NMR (101 MHz, $CDCl_3$) $\delta$ 150.4 (C-2), 148.4 (C-4), 141.9 (C-8), 138.6 (C-5), 134.2 (C-7), 126.4 (C-6), 121.7 (C-3), -1.1 (C-9); ESI-HRMS $m/z$ calcd. for $C_{14}H_{18}NSi$ $[M+H]^+$: 228.1203; found: 228.1207.

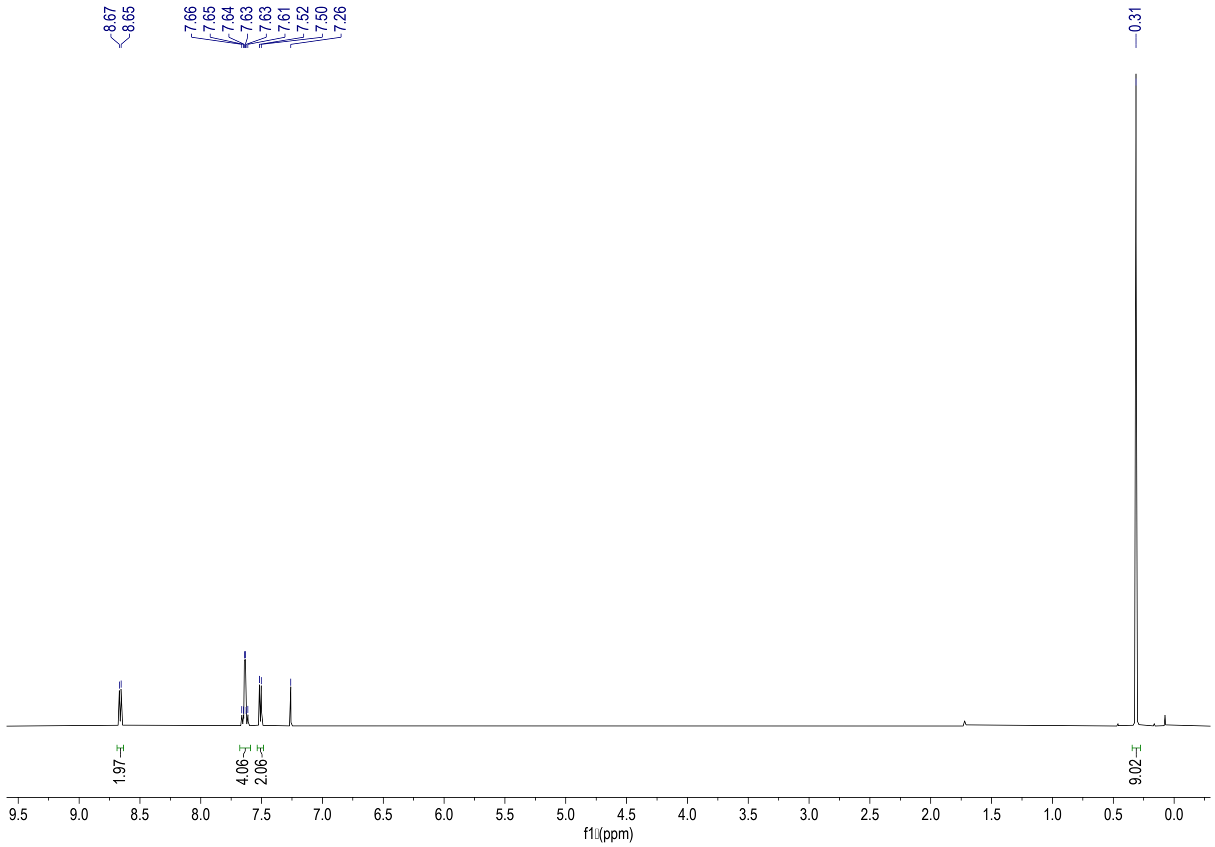


$^{1}$H-NMR spectrum of **6** (400 MHz, $CDCl_3$)

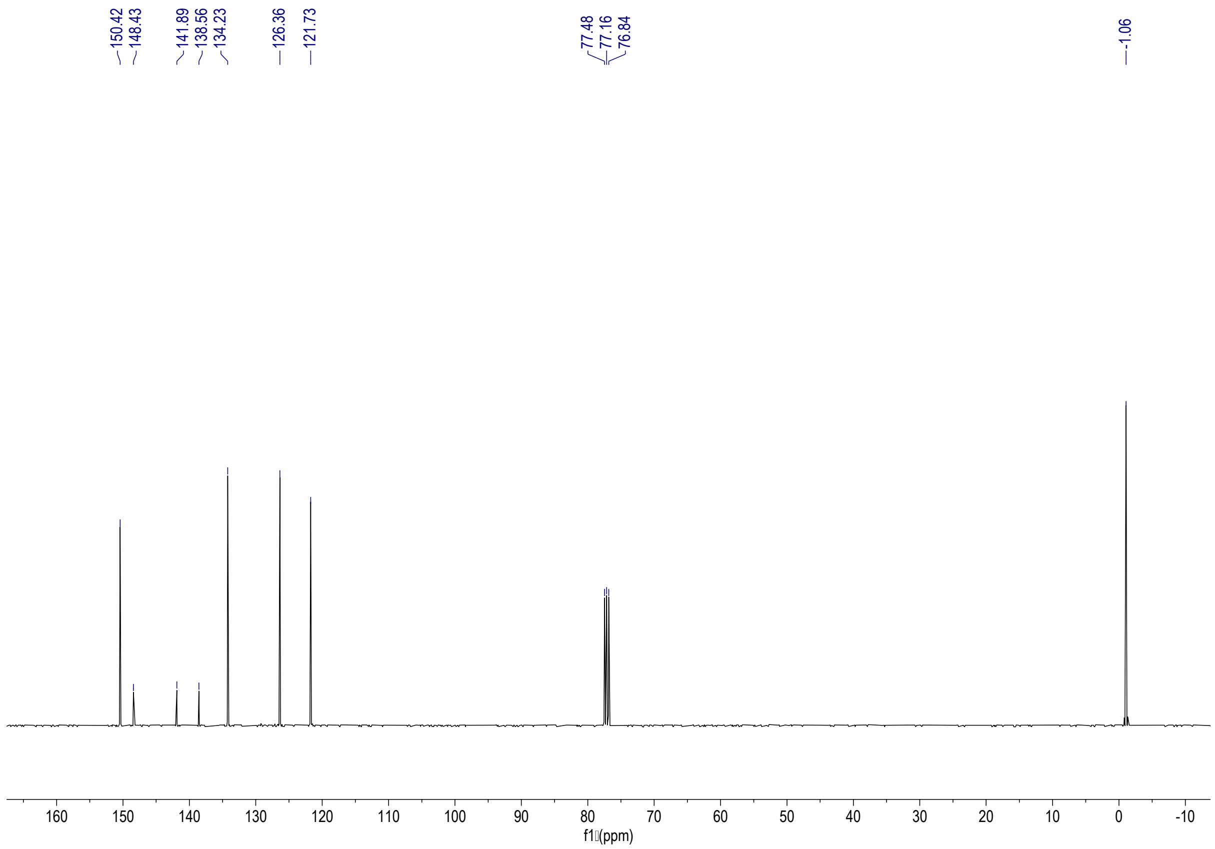


$^{13}$C-NMR spectrum of **6** (101 MHz, $CDCl_3$)

**4-(4-nitrosophenyl)pyridine (7)**

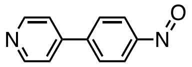


To a solution of compound **6** (900 mg, 3.96 mmol) in ACN, $NOBF_4$ (1.12 g, 9.11 mmol) was added. The reaction mixture was stirred at r.t. for 30 min. The mixture was diluted with DCM (100 mL) and washed with sat. $NaHCO_3$ solution (80 mL). The aqueous layer was extracted with DCM. The combined organic layers were dried over anh. $MgSO_4$, filtered and concentrated. The crude was purified by silica gel chromatography column (EtOAc/hexane/toluene, 3:1:1) to give compound **7** (405 mg, 56%) as a yellow solid. $^1$H NMR (400 MHz, $CDCl_3$) $\delta$ 8.75 (d, $J$ = 6.1 Hz, 2H, H-2), 8.02 (d, $J$ = 8.7 Hz, 2H, H-7), 7.88 (d, $J$ = 8.7 Hz, 2H, H-6), 7.56 (d, $J$ = 6.2 Hz, 2H, H-3); $^{13}$C NMR (101 MHz, $CDCl_3$) $\delta$ 164.6 (C-8), 150.7 (C-2), 146.5 (C-4), 144.8 (C-5), 128.3 (C-6), 121.9 (C-3), 121.7 (C-4); ESI-HRMS *m/z* calcd. for $C_{11}H_9N_2O$ $[M+H]^+$: 185.0709; found: 185.0718.

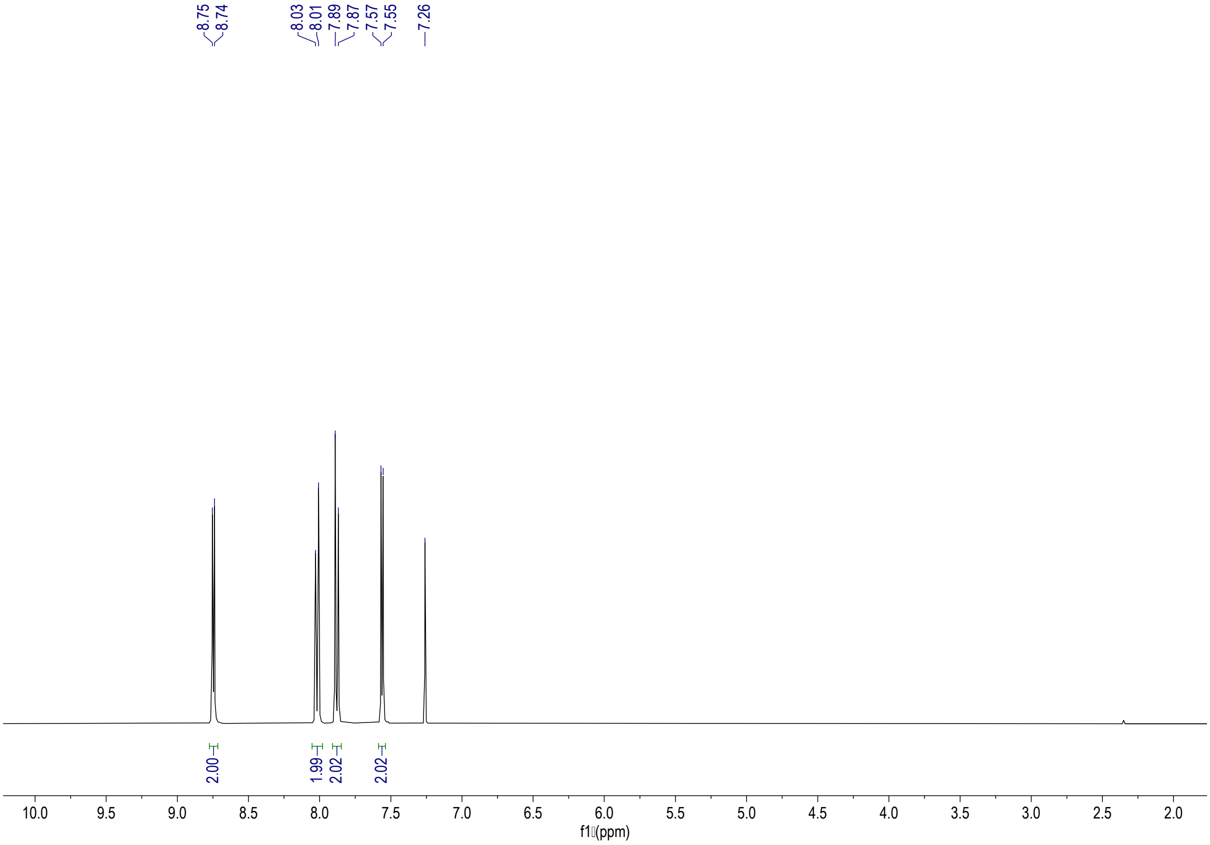


$^1$H-NMR spectrum of **7** (400 MHz, $CDCl_3$)

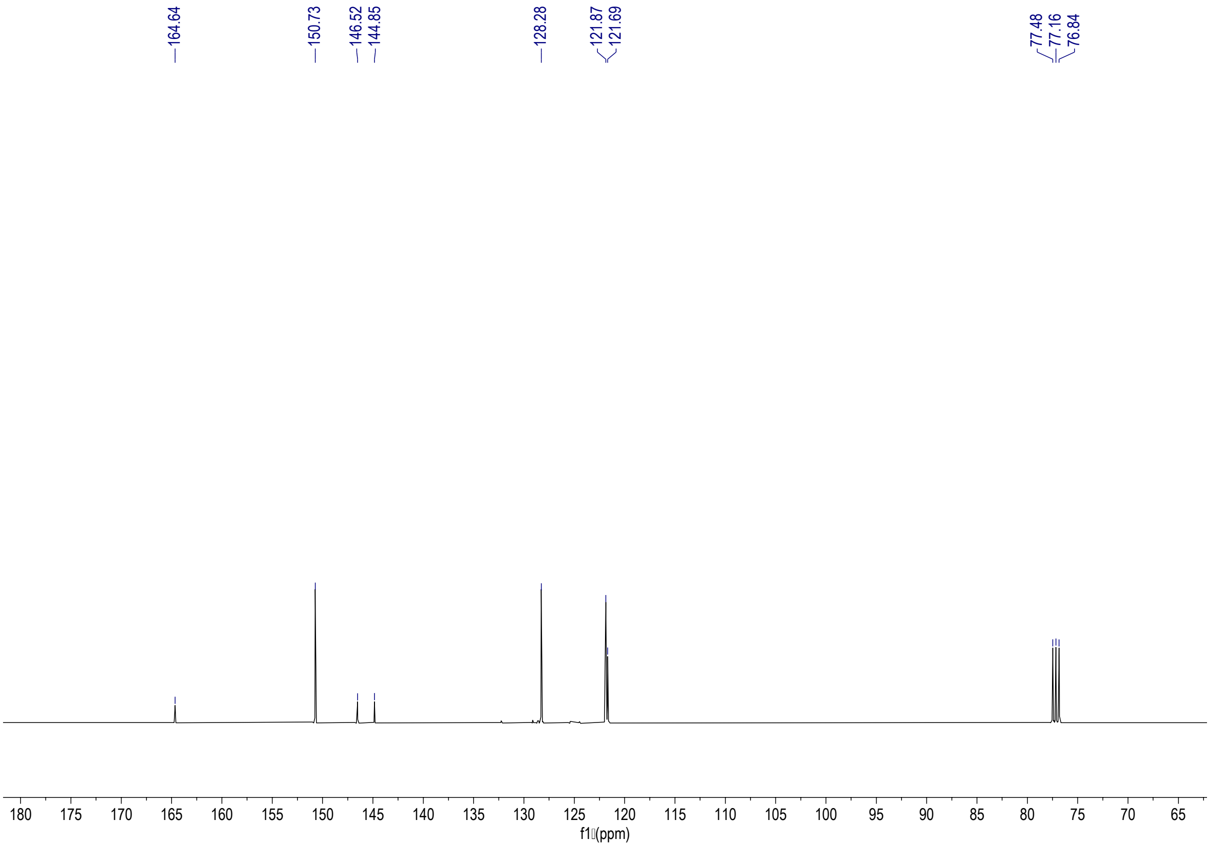


$^{13}$C-NMR spectrum of **7** (101 MHz, $CDCl_3$)

**2-(2-bromophenylthio)acetic acid (8)**

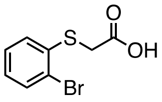


To a stirred solution of NaOH (1.2 g, 30.78 mmol) in MeOH (30 mL) was added dropwise 2-bromothiophenol (3 g, 15, 39 mmol) over a period of 20 min. Once completation of the addition and when no longer the reaction was exothermic, chloroacetic acid (1.76 g, 18.47 mmol) was added portion wise, and the reaction mixture was refluxed overnight. The reaction mixture was then poured into ice-cold HCl (3.5 mL) solution and was stirred for 10 min. The solution was extracted with DCM. The organic phase was washed with water and then dried over anh. $MgSO_4$, filtered and concentrated. The solid was washed with toluene and hexane to give 2-(2-bromophenylthio)acetic acid (**8**) (2.8 g, 74%) as a white solid. $^1$H NMR (400 MHz, DMSO-$d_6$) $\delta$ 7.60 (dd, $J$ = 7.9, 1.3 Hz, 1H, H-3), 7.37 (ddd, $J$ = 7.9, 7.2, 1.3 Hz, 1H, H-5), 7.30 (dd, $J$ = 8.0, 1.6 Hz, 1H, H-6), 7.10 (ddd, $J$ = 7.9, 7.2, 1.7 Hz, 1H, H-4), 3.89 (s, 2H, H-7); $^{13}$C NMR (101 MHz, DMSO-$d_6$) $\delta$ 170.2 (C-8), 137.2 (C-1), 132.7 (C-3), 128.4 (C-5), 126.8 (C-6), 126.7 (C-4), 120.9 (C-2), 34.2 (C-7); ESI-HRMS $m/z$ calcd. for $C_8H_6O_2S^{79}Br$ $[M\text{-}H]^-$: 244.9272, found: 244.9267; $m/z$ calcd. for $C_8H_6O_2S^{81}Br$ $[M\text{-}H]^-$: 246.9251; found: 246.9255.

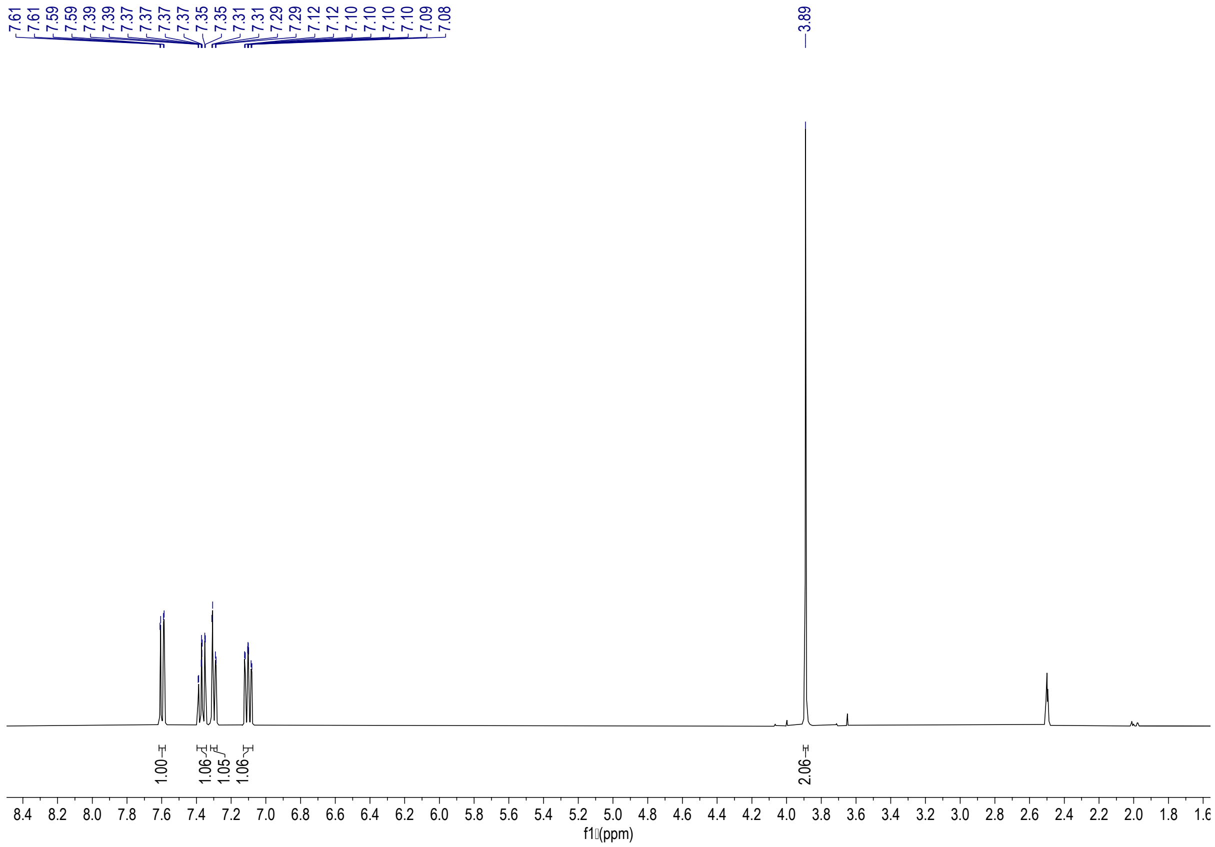


$^{1}$H-NMR spectrum of **8** (400 MHz, DMSO-$d_6$)

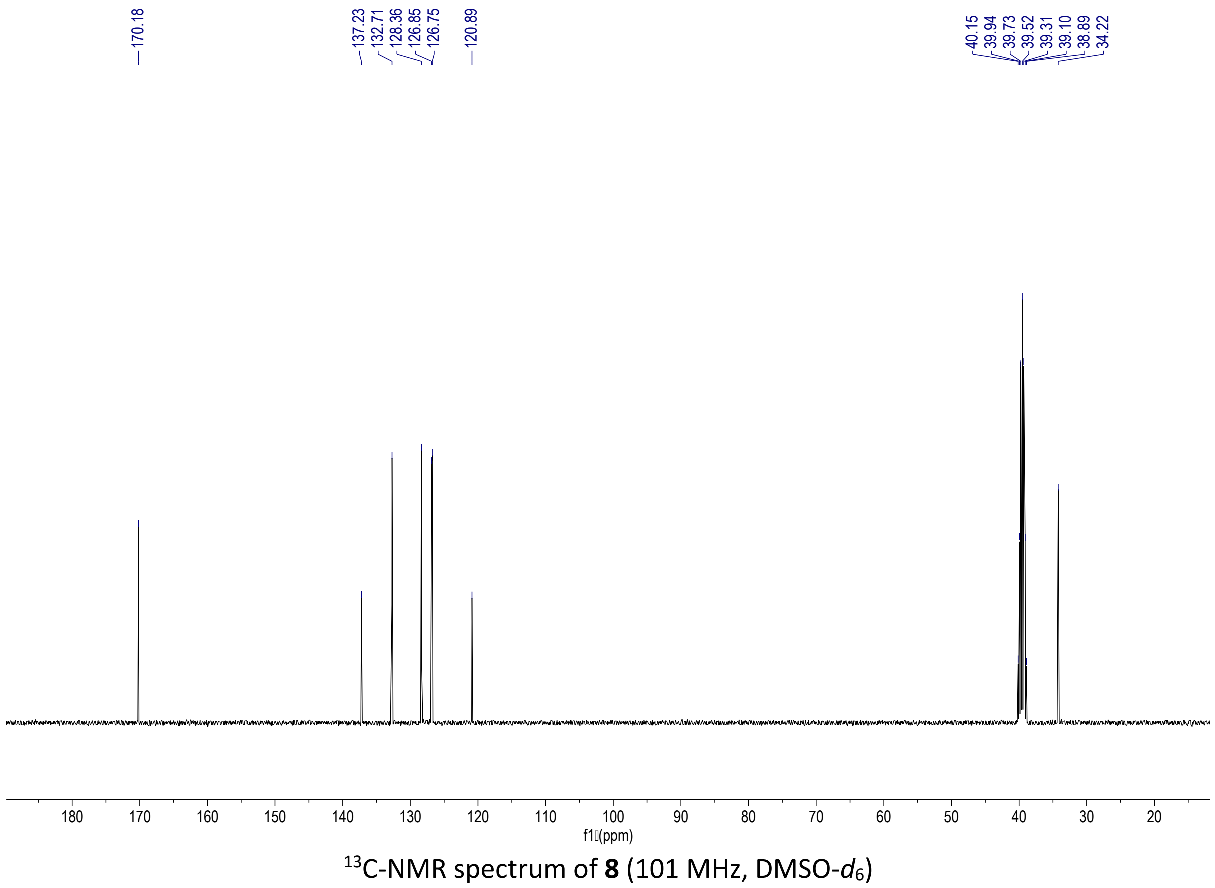


$^{13}$C-NMR spectrum of **8** (101 MHz, DMSO-$d_6$)

**7-bromobenzo[*b*]thiophen-3(2*H*)-one (9)**

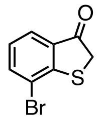


To a solution of compound **8** (1.2 g, 4.88 mmol) in dry DCM (10 mL), oxalylchloride (3.9 mL, 7.81 mmol) and DMF (1 drop) were added. The reaction mixture was stirred at r.t. for 1 h. After no gas formation was observed, the mixture was concentrated in vacuo to remove all solvents and remaining oxalylchloride. To a solution of the resulting crude in DCE (15 mL) cooled to 0 °C, $AlCl_3$ (1.04 g, 7.81 mmol) was added portion wise. The reaction mixture was allowed to reach r.t. and stirred overnight. The mixture was diluted with ice water and extracted twice with DCM. The combined organic layers were washed with water and then dried over anh. $MgSO_4$, filtered and concentrated. The crude was purified by silica gel column chromatography (toluene/hexane, 5/1) to give compound **9** (924 mg, 83%) as a white solid. $^1$H NMR (400 MHz, $CDCl_3$) $\delta$ 7.74 (td, $J$ = 7.5, 1.1 Hz, 2H, H-3, H-5), 7.14 (t, $J$ = 7.7 Hz, 1H, H-4), 3.84 (s, 2H, H-8); $^{13}$C NMR (101 MHz, $CDCl_3$) $\delta$ 199.6 (C-7), 155.5 (C-1), 138.2 (C-3), 133.1 (C-6), 126.3 (C-4), 125.4 (C-5), 118.9 (C-2), 40.3 (C-8); ESI-HRMS $m/z$ calcd for $C_8H_6BrOS$ $[M+H]^+$: 228.9317; found: 228.9312.

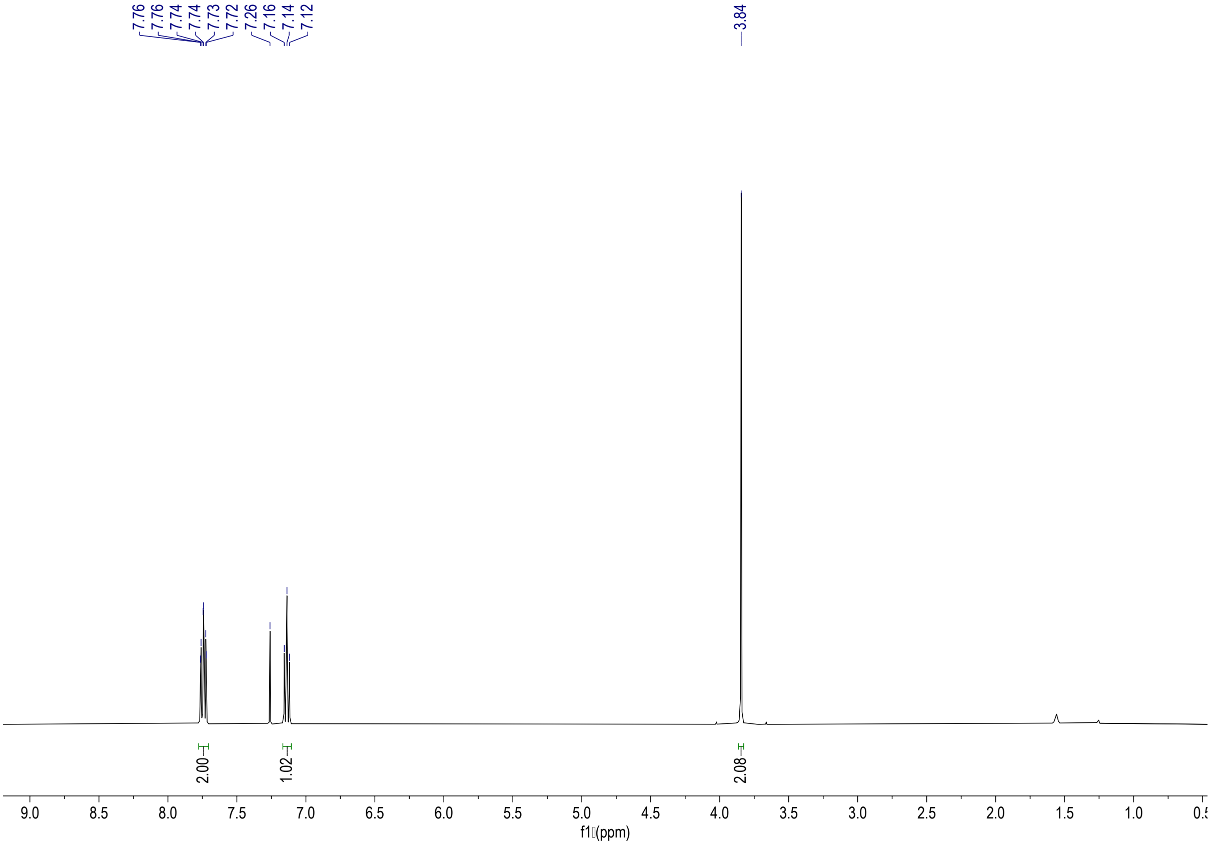


$^1$H-NMR spectrum of **9** (400 MHz, $CDCl_3$)

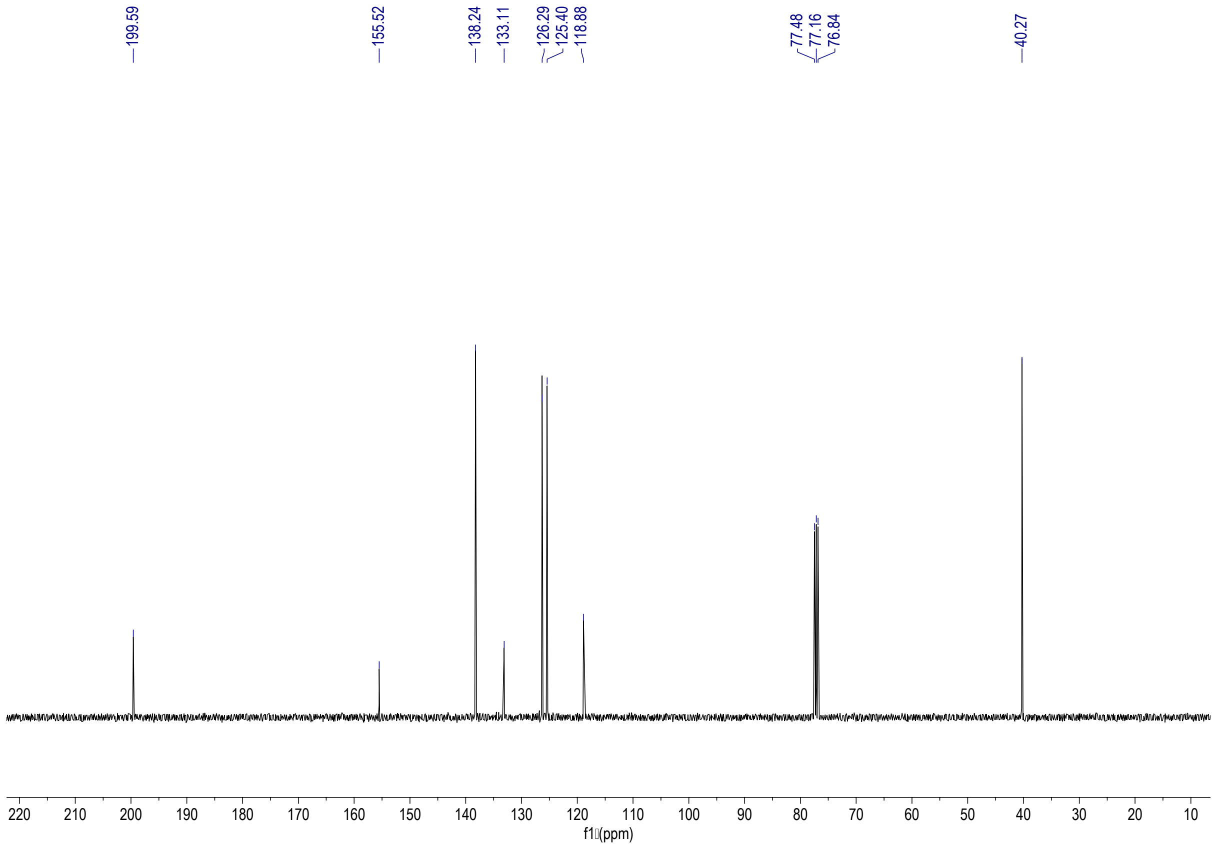


$^{13}$C-NMR spectrum of **9** (101 MHz, $CDCl_3$)

**7-(pyridin-4-yl)benzo[*b*]thiophen-3(2*H*)-one (10)**

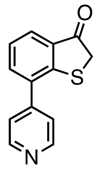


To a solution of compound **9** (800 mg, 3.51 mmol), $Pd(PPh_3)_4$ (410 mg, 0.35 mmol), ethylene glycol (1 drop) and 4-pyridinylboronic acid (719 mg, 5.26 mmol) in dry THF (20 mL), a 2.5 M aq. $K_2CO_3$ solution (4 mL) (previously degassed by evacuation under vacuum and backfilling with nitrogen three times) was added. The reaction mixture was stirred overnight at 70 °C. After thar, the mixture was cooled to r.t. and water was added. The aqueous layer was extracted with DCM and the organic phase was dried over anh. $MgSO_4$, filtered and concentrated. The crude was purified by silica gel column chromatography (EtOAc/DCM, 1:1) to give compound **10** (177 mg, 22%) as a pink solid. $^1$H NMR (400 MHz, $CDCl_3$) $\delta$ 8.74 (d, *J* = 6.1 Hz, 2H, H-11), 7.86 (dd, *J* = 7.7, 1.3 Hz, 1H, H-5), 7.61 (dd, *J* = 7.4, 1.3 Hz, 1H, H-3), 7.52 (d, *J* = 6.1 Hz, 2H, H-10), 7.37 (t, *J* = 7.6 Hz, 1H, H-4), 3.82 (s, 2H, H-8); $^{13}$C NMR (101 MHz, $CDCl_3$) $\delta$ 199.8 (C-7), 152.5 (C-1), 150.4 (C-11), 146.1 (C-9), 135.6 (C-3), 135.4 (C-2), 132.1 (C-6), 127.0 (C-5), 125.8 (C-4), 122.8 (C-10), 39.5 (C-8); ESI-HRMS *m/z* calcd. for $C_{13}H_{10}NOS$ $[M+H]^+$: 228.0478; found: 228.0485.

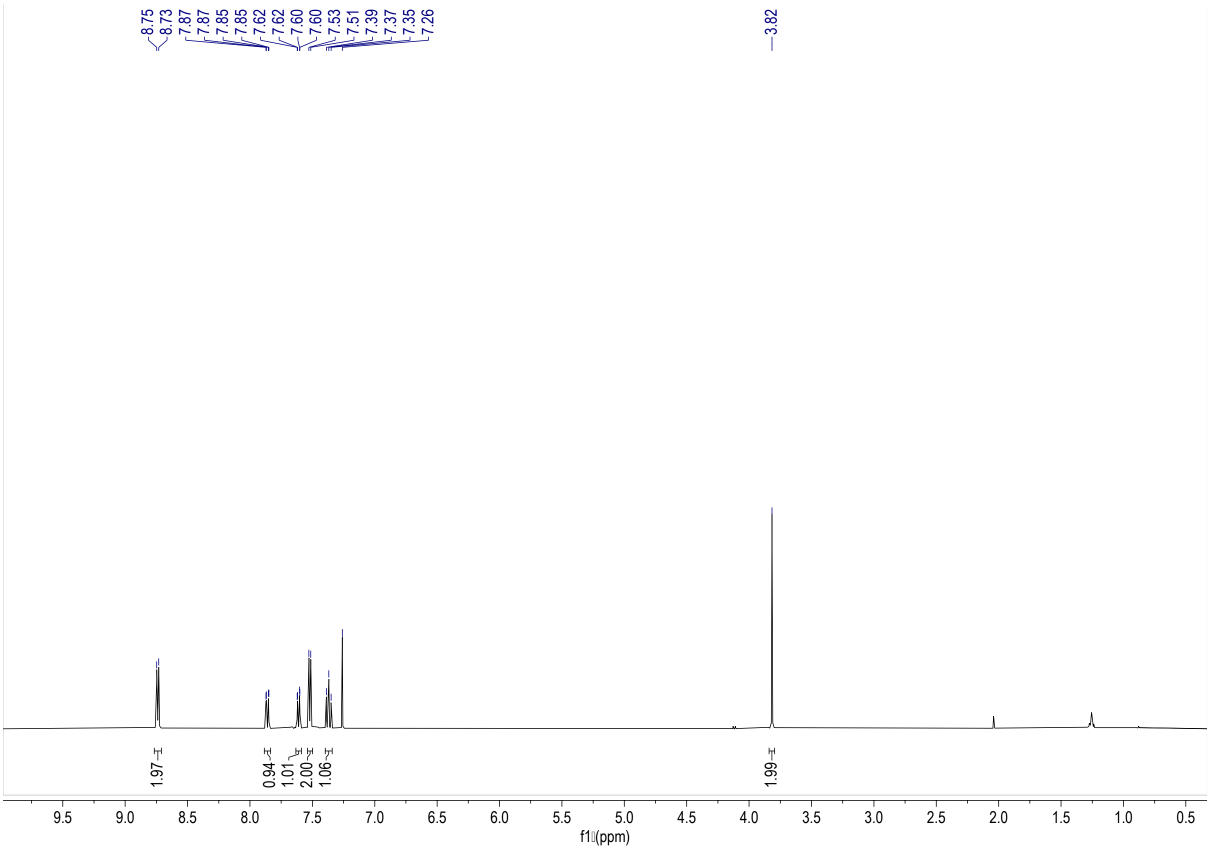


[1]H-NMR spectrum of **10** (400 MHz, $CDCl_3$)

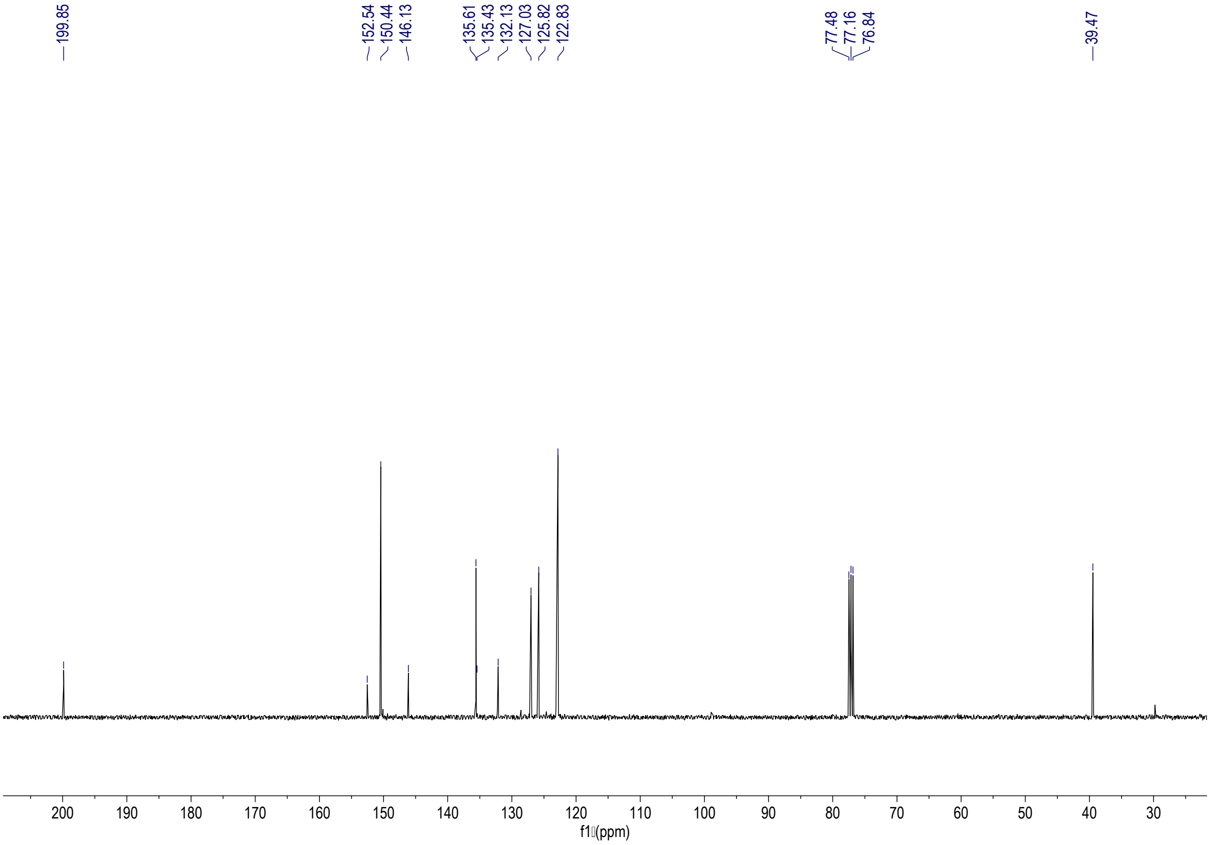


[13]C-NMR spectrum of **10** (101 MHz, $CDCl_3$)

**(*Z*)-7-(pyridin-4-yl)-2-((4-(pyridin-4-yl)phenyl)imino)benzo[*b*]thiophen-3(2*H*)-one (11)**

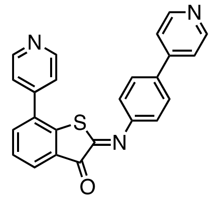

To a solution of compound **10** (135 mg, 0.59 mmol) in benzene (15 mL), compound **7** (197 mg, 1.07 mmol) and piperidine (4 drops) were added. The reaction mixture was stirred at reflux for 1 h. The mixture was diluted with DCM and washed with water. The aqueous layer was extracted twice with DCM. The combined organic layers were dried over anh. $MgSO_4$, filtered and concentrated. The crude was purified by silica gel column chromatography ($CHCl_3$/MeOH, 40:1) to give compound **11** (186 mg, 80%) as a redish solid. $^1H$ NMR (400 MHz, $CDCl_3$) $\delta$ 8.73 (dd, $J$ = 4.4, 1.7 Hz, 2H), 8.65 (dd, $J$ = 4.4, 1.7 Hz, 2H), 8.07 (dd, $J$ = 7.6, 1.3 Hz, 1H), 7.73 (m, 2H), 7.68 (dd, $J$ = 7.6, 1.4 Hz, 1H), 7.55 – 7.48 (m, 3H), 7.40 (dd, $J$ = 4.4, 1.7 Hz, 2H), 7.35 (m, 2H); $^{13}C$ NMR (101 MHz, $CDCl_3$) $\delta$ 185.3, 156.6, 150.7, 150.5, 149.9, 147.1, 145.1, 142.6, 137.2, 137.1, 135.9, 128.6, 128.2, 127.9, 127.5, 122.7, 121.8, 121.4; ESI-HRMS $m/z$ calcd. for $C_{24}H_{16}N_3OS$ $[M+H]^+$: 394.1009; found: 394.1015.

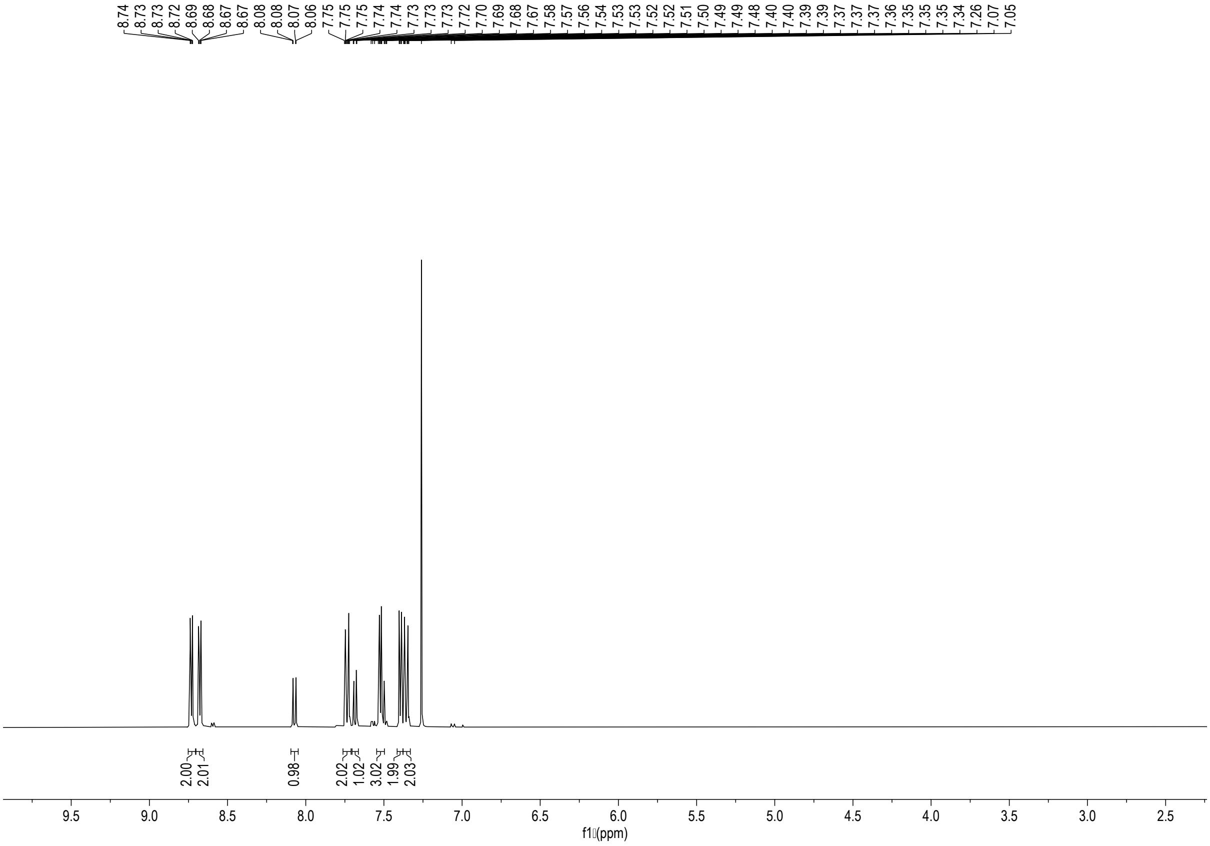

$^1H$-NMR spectrum of **11** (400 MHz, $CDCl_3$)

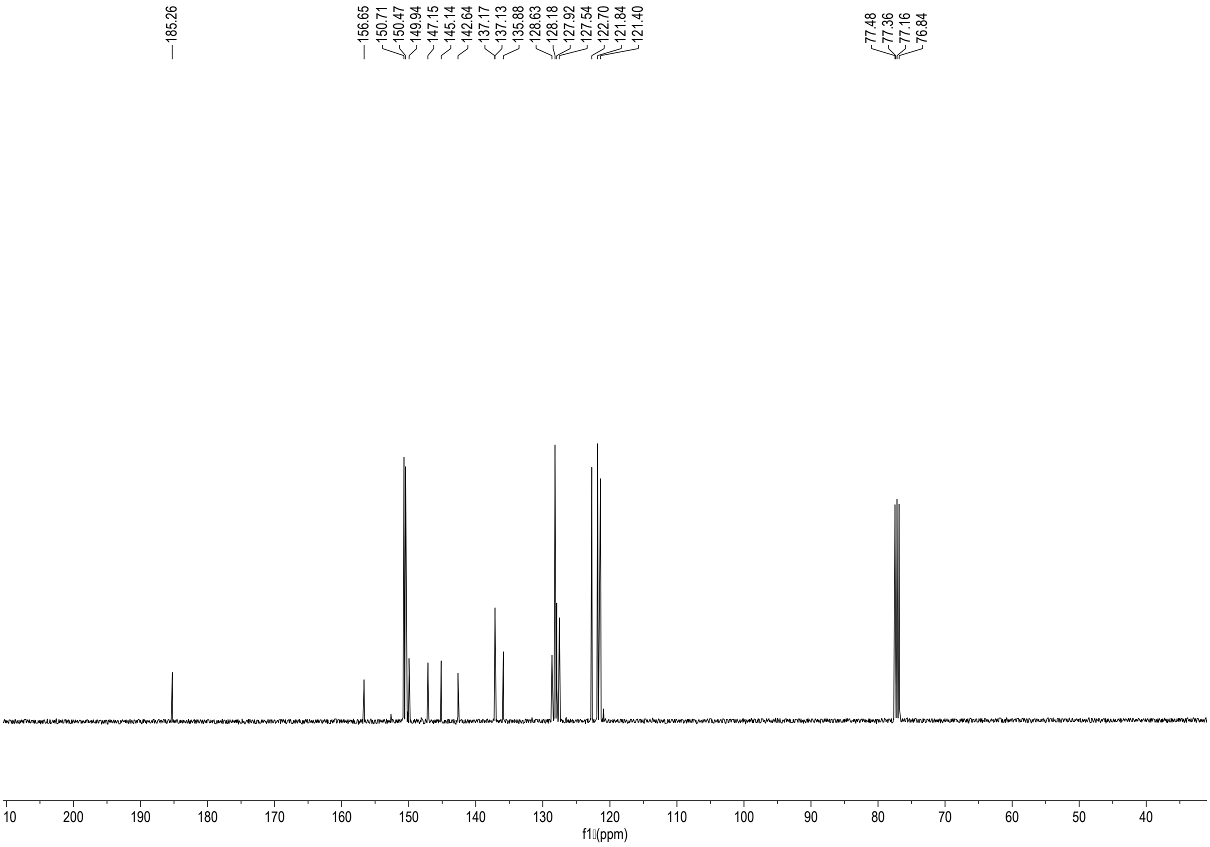


$^{13}$C-NMR spectrum of **11** (101 MHz, $CDCl_3$)

**Compound L10**

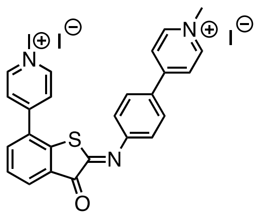


To a solution of compound **11** (130 mg, 0.33 mmol) in ACN/$CHCl_3$ (1:1, 8 mL), methyl iodide (182 µL, 1.32 mmol) was added. The solution was stirred in a sealed vessel at 50 °C overnight. The generated solid was filtered and washed with diethyl ether. Following filtration, compound **L10** (186 mg, 83%) was obtained as an orange powder. $^{1}$H NMR (400 MHz, DMSO-$d_6$) $\delta$ 9.03 (dd, $J$ = 13.1, 6.6 Hz, 4H), 8.52 (d, $J$ = 6.9 Hz, 2H), 8.29 (d, $J$ = 6.8 Hz, 2H), 8.22 (d, $J$ = 8.6 Hz, 2H), 8.16 (dd, $J$ = 7.6, 1.3 Hz, 1H), 8.07 (dd, $J$ = 7.7, 1.3 Hz, 1H), 7.74 (t, $J$ = 7.7 Hz, 1H), 7.40 (d, $J$ = 8.6 Hz, 2H), 4.34 (s, 6H); $^{13}$C NMR (101 MHz, DMSO-$d_6$) $\delta$ 183.5, 157.5, 153.7, 153.5, 153.0, 152.8, 152.0, 146.2, 145.6, 144.3, 141.7, 138.0, 132.1, 131.6, 129.8, 129.7, 129.0, 128.5, 128.2, 126.4, 123.8, 120.8, 120.4, 114.0, 47.8, 47.1; ESI-HRMS *m/z* calcd. for $C_{26}H_{21}N_3OSI$ $[M]^+$: 550.0450; found: 550.0451.

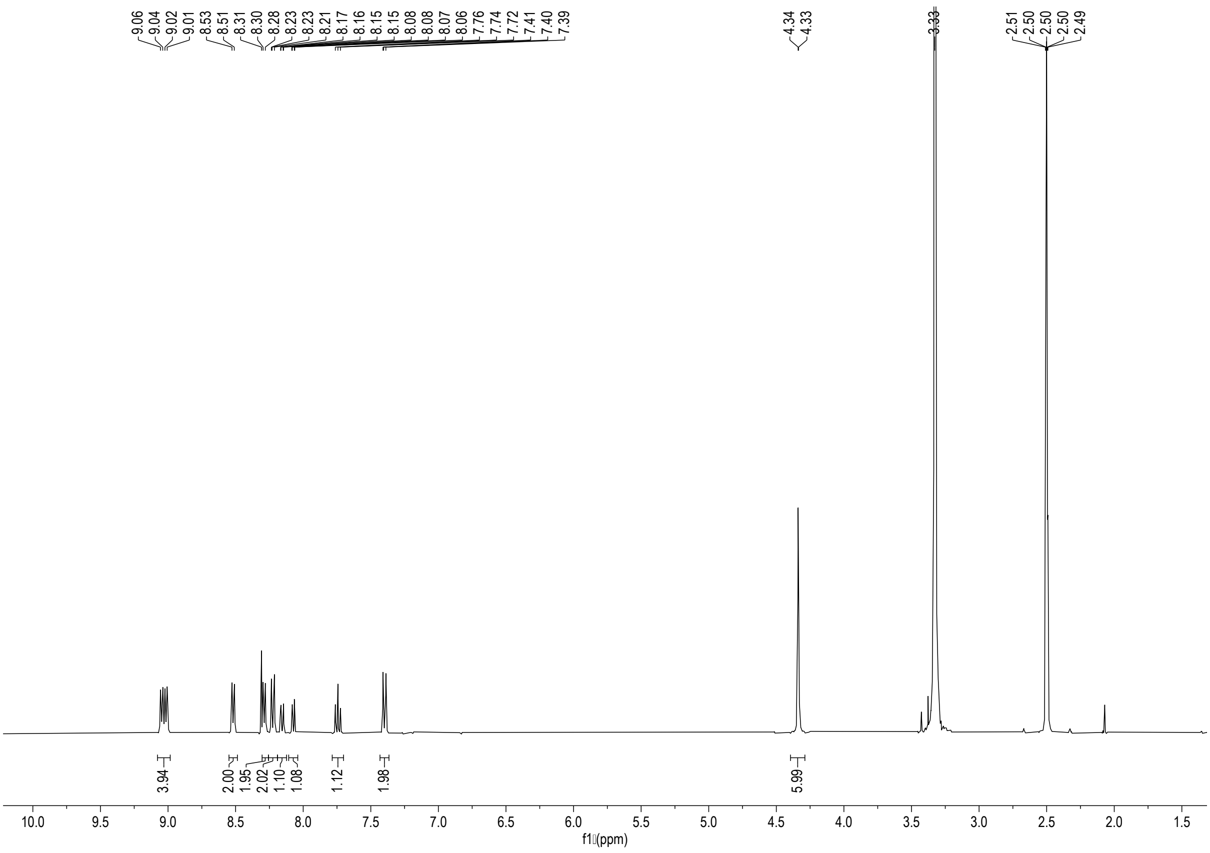


$^{1}H$-NMR spectrum of **L10** (400 MHz, DMSO-$d_6$)

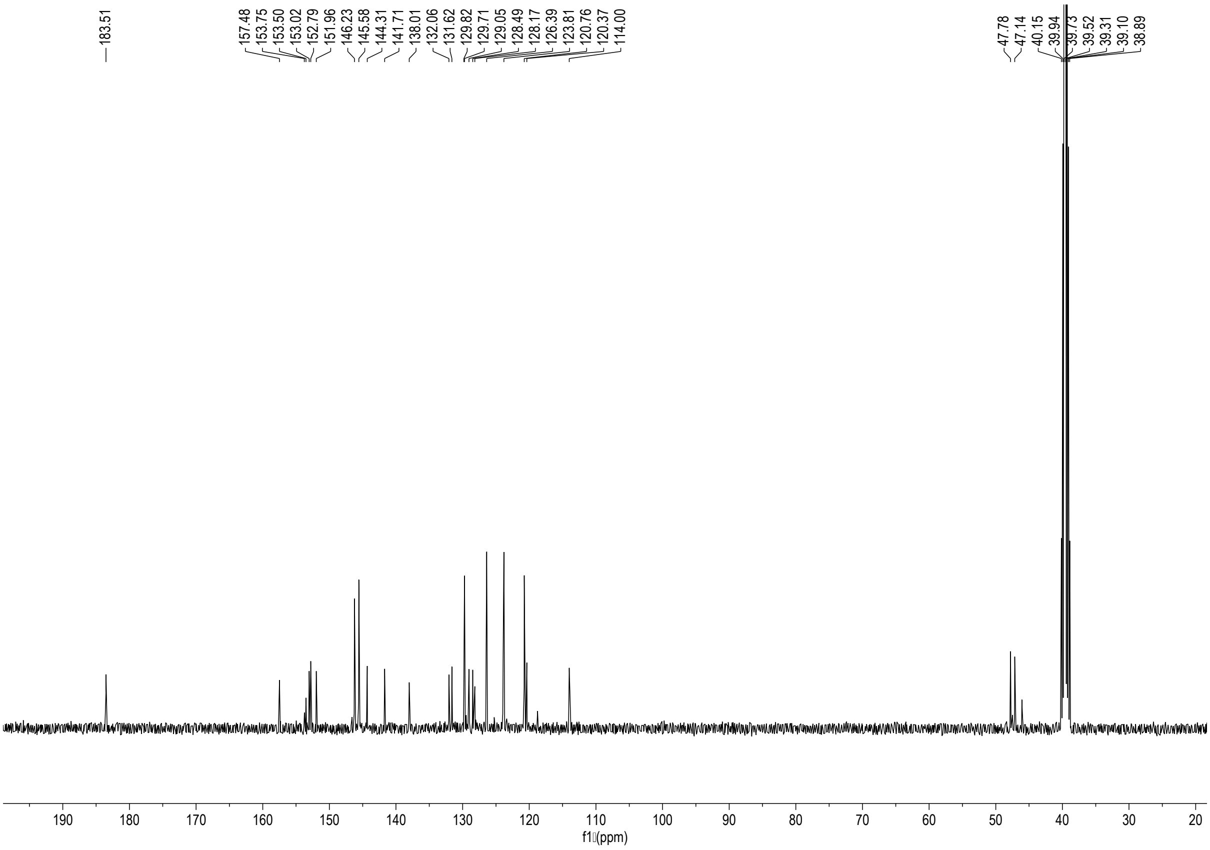


$^{13}C$-NMR spectrum of **L10** (101 MHz, DMSO-$d_6$)

**Synthesis of L11**

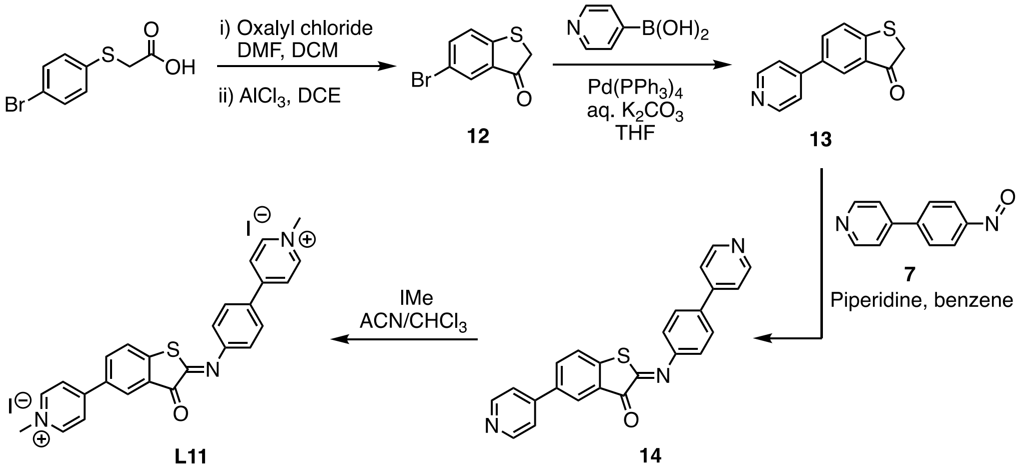

**5-bromobenzo[*b*]thiophen-3(2*H*)-one (12)**

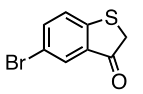

To a solution of 2-(4-bromophenylthio)acetic acid (1.4 g, 5.69 mmol) in dry DCM (10 mL), oxalylchloride (4.6 mL, 9.11 mmol) and DMF (1 drop) were added. The reaction mixture was stirred at r.t. for 1 h. After no gas formation was observed, the mixture was concentrated in vacuo to remove all solvents and remaining oxalylchloride. To a solution of the resulting crude in DCE (15 mL) cooled to 0 °C, $AlCl_3$ (1.2 g, 9.11 mmol) was added portion wise. The reaction mixture was allowed to reach r.t. and stirred overnight. The mixture was diluted with ice water and extracted twice with DCM. The combined organic layers were washed with water and then dried over anh. $MgSO_4$, filtered and concentrated. The crude was purified by silica gel column chromatography (toluene/hexane, 5/1) to give compound **12** (1.02 g, 79%) as a light pink solid. $^1$H NMR (400 MHz, $CDCl_3$) $\delta$ 7.89 (dd, $J$ = 2.1, 0.5 Hz, 1H, H-4), 7.63 (dd, $J$ = 8.4, 2.1 Hz, 1H, H-6), 7.31 (dd, $J$ = 8.4, 0.5 Hz, 1H, H-7), 3.83 (s, 2H, H-2); $^{13}$C NMR (101 MHz, $CDCl_3$) $\delta$ 198.6 (C-3), 153.0 (C-7a), 138.4 (C-6), 132.7 (C-3a), 129.4 (C-4), 126.0 (C-7), 118.5 (C-5), 39.9 (C-2); APCI-HRMS *m/z* calcd. for $C_8H_6BrOS$ $[M+H]^+$: 228.9317; found: 228.9314.

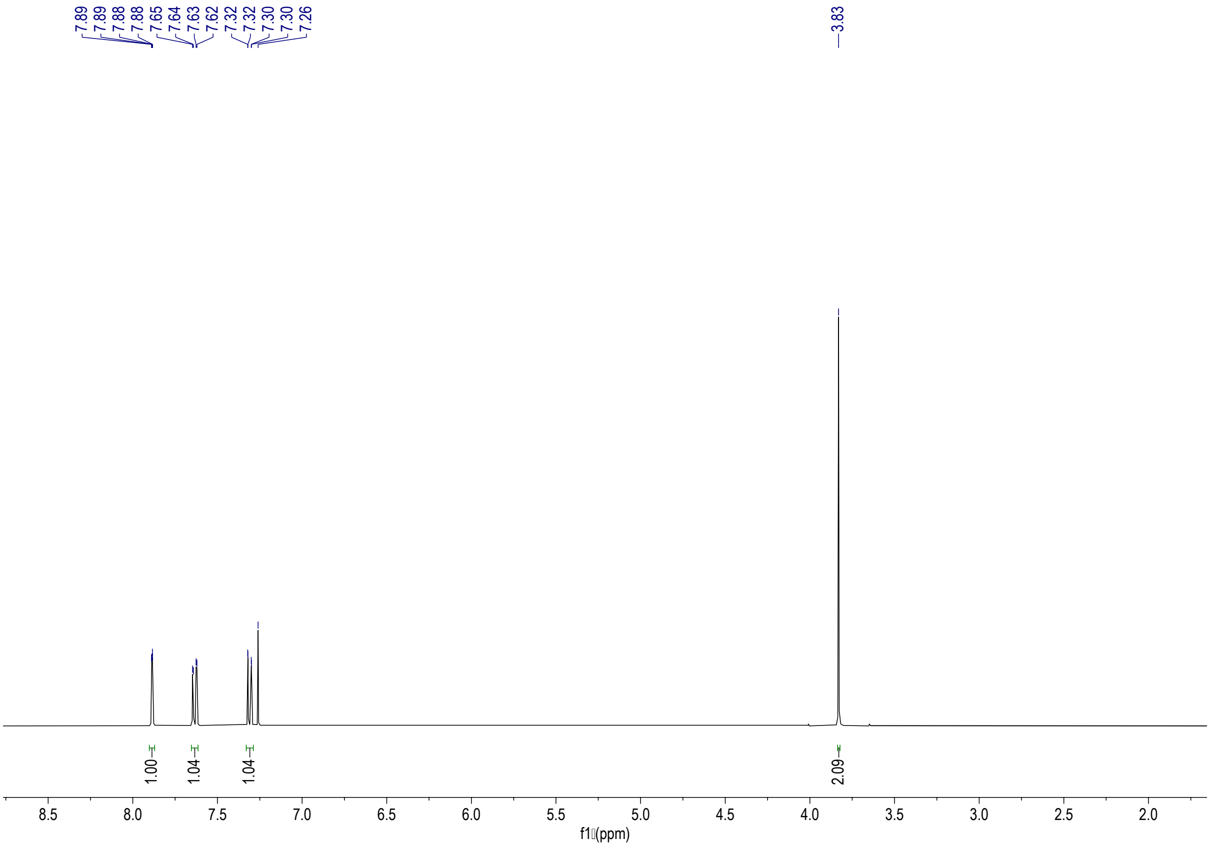


$^{1}$H-NMR spectrum of **12** (400 MHz, $CDCl_3$)

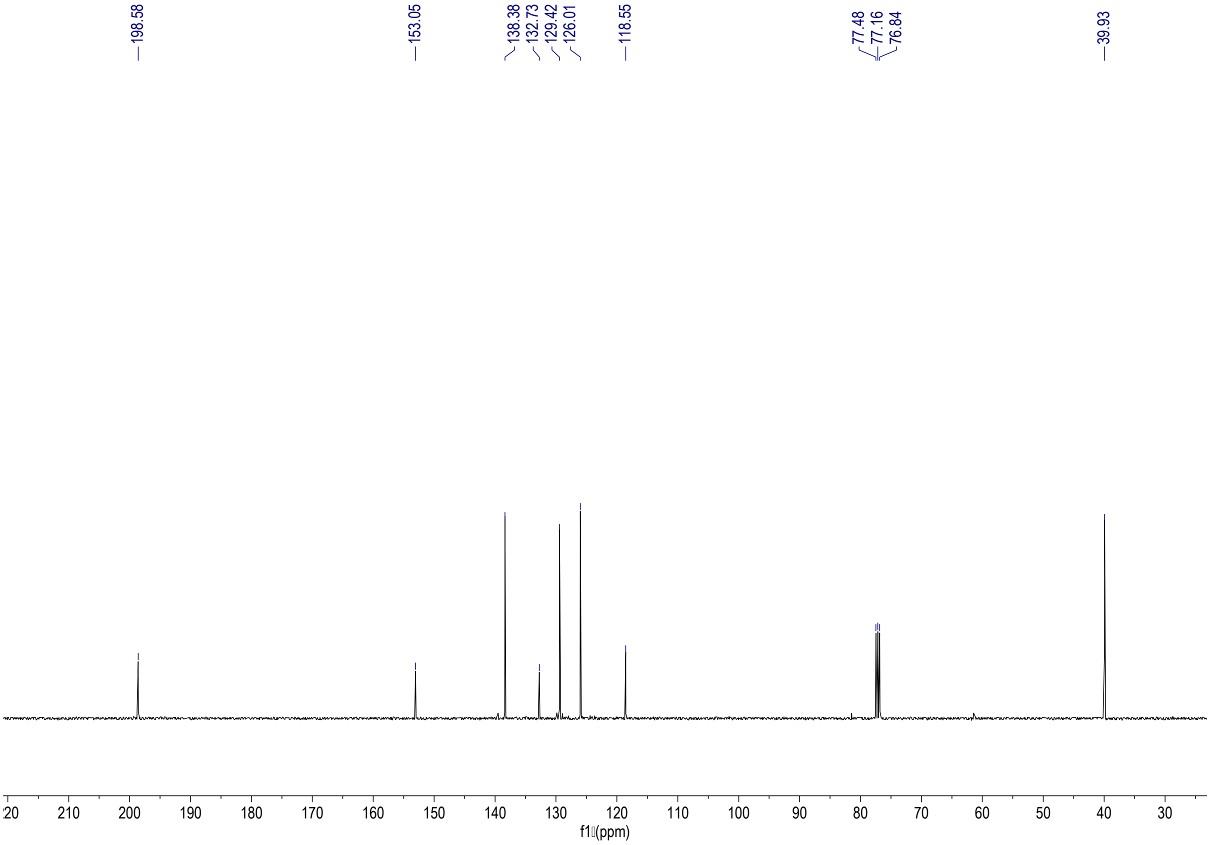


$^{13}$C-NMR spectrum of **12** (101 MHz, $CDCl_3$)

**(*Z*)-5-(pyridin-4-yl)-2-((4-(pyridin-4-yl)phenyl)imino)benzo[*b*]thiophen-3(2*H*)-one (14)**

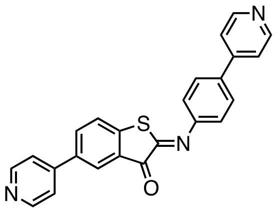


To a solution of compound **12** (400 mg, 1.76 mmol), $Pd(PPh_3)_4$ (205 mg, 0.18 mmol), ethylene glycol (1 drop) and 4-pyridinylboronic acid (360 mg, 2.63 mmol) in dry THF (15 mL), a 2.5 M aq. $K_2CO_3$ solution (2 mL) ( previously degassed by evacuation under vacuum and backfilling with nitrogen three times) was added. The reaction mixture was stirred overnight at 70 °C. After thar, the mixture was cooled to r.t. and water was added. The aqueous layer was extracted with DCM and the organic phase was dried over anh. $MgSO_4$, filtered and concentrated. The crude was purified by silica gel column chromatography (EtOAc/DCM, 1:1) to give compound **13** (225 mg, 57%) as a pink solid. To a solution of compound **13** (80 mg, 0.35 mmol) in benzene (8 mL), compound **7** (117 mg, 0.63 mmol) and piperidine (2 drops) were added. The reaction mixture was stirred at reflux for 1 h. The mixture was diluted with DCM and washed with water. The aqueous layer was extracted twice with DCM. The combined organic layers were dried over anh. $MgSO_4$, filtered and concentrated. The crude was purified by silica gel column chromatography ($CHCl_3$/MeOH, 40:1) to give compound **14** (101 mg, 73%) as a redish solid. $^1$H NMR (400 MHz, $CDCl_3$) $\delta$ 8.73 (d, $J$ = 6.2 Hz, 2H, H-10), 8.69 (d, $J$ = 6.1 Hz, 2H, H-17), 8.26 (dd, $J$ = 2.1, 0.6 Hz, 1H, H-4), 7.93 (dd, $J$ = 8.2, 2.0 Hz, 1H, H-6), 7.77 (d, $J$ = 8.5 Hz, 2H, H-13), 7.57 (dd, $J$ = 8.2, 0.6 Hz, 1H, H-7), 7.54 (ddd, $J$ = 4.5, 3.5, 1.7 Hz, 4H, H-9, H-16), 7.41 (d, $J$ = 8.6 Hz, 2H, H-12); $^{13}$C NMR (101 MHz, $CDCl_3$) $\delta$ 184.9 (C-3), 156.6 (C-2), 150.7 (C-10), 150.5 (C-17), 149.8 (C-11), 147.1 (C-15), 145.8 (C-8), 144.9 (C-3a), 137.3, 137.2 (C-7a, C-14), 135.4 (C-6), 128.4 (C-5), 128.1 (C-13), 126.0 (C-4), 125.8 (C-7), 122.0 (C-12), 121.4 (C-16), 121.2 (C-9); ESI-HRMS *m/z* calcd. for $C_{24}H_{16}N_3OS$ $[M+H]^+$: 394.1009; found: 394.0990.

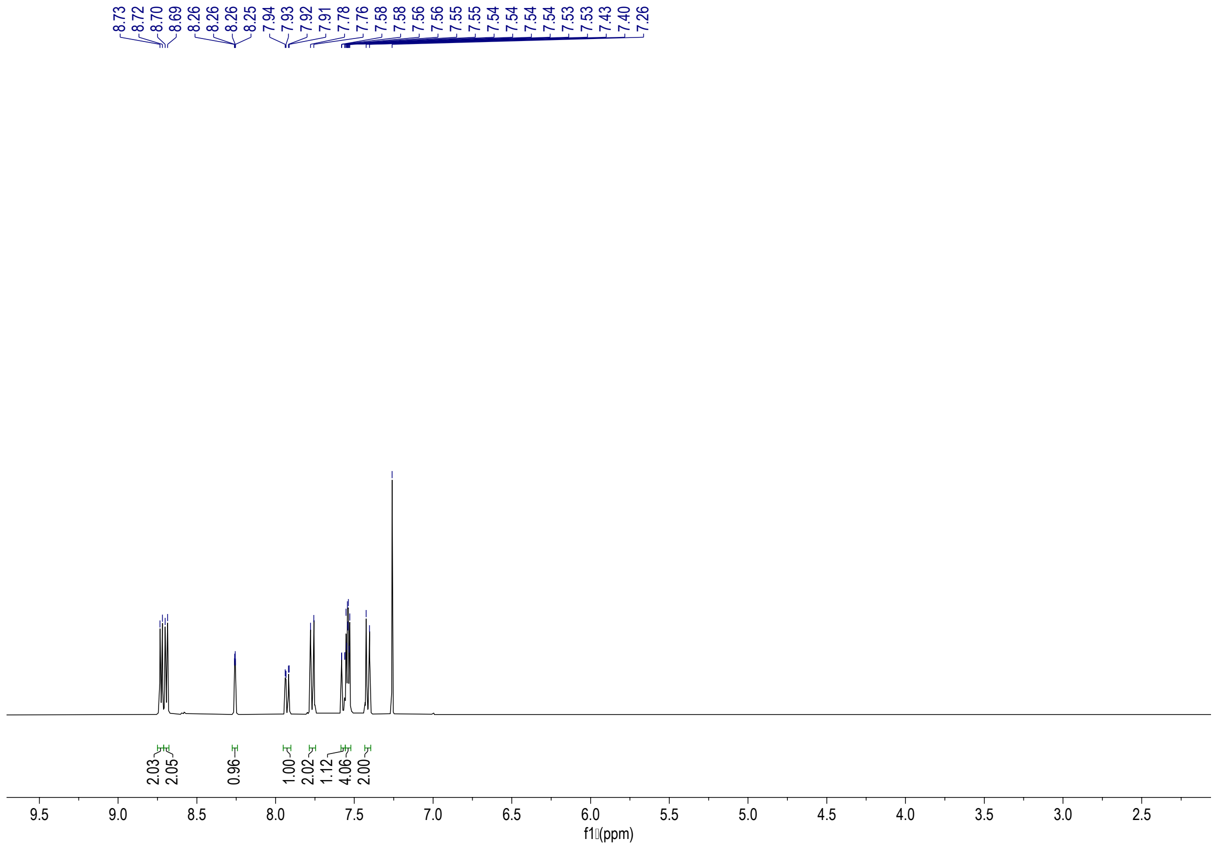


$^{1}$H-NMR spectrum of **14** (400 MHz, $CDCl_3$)

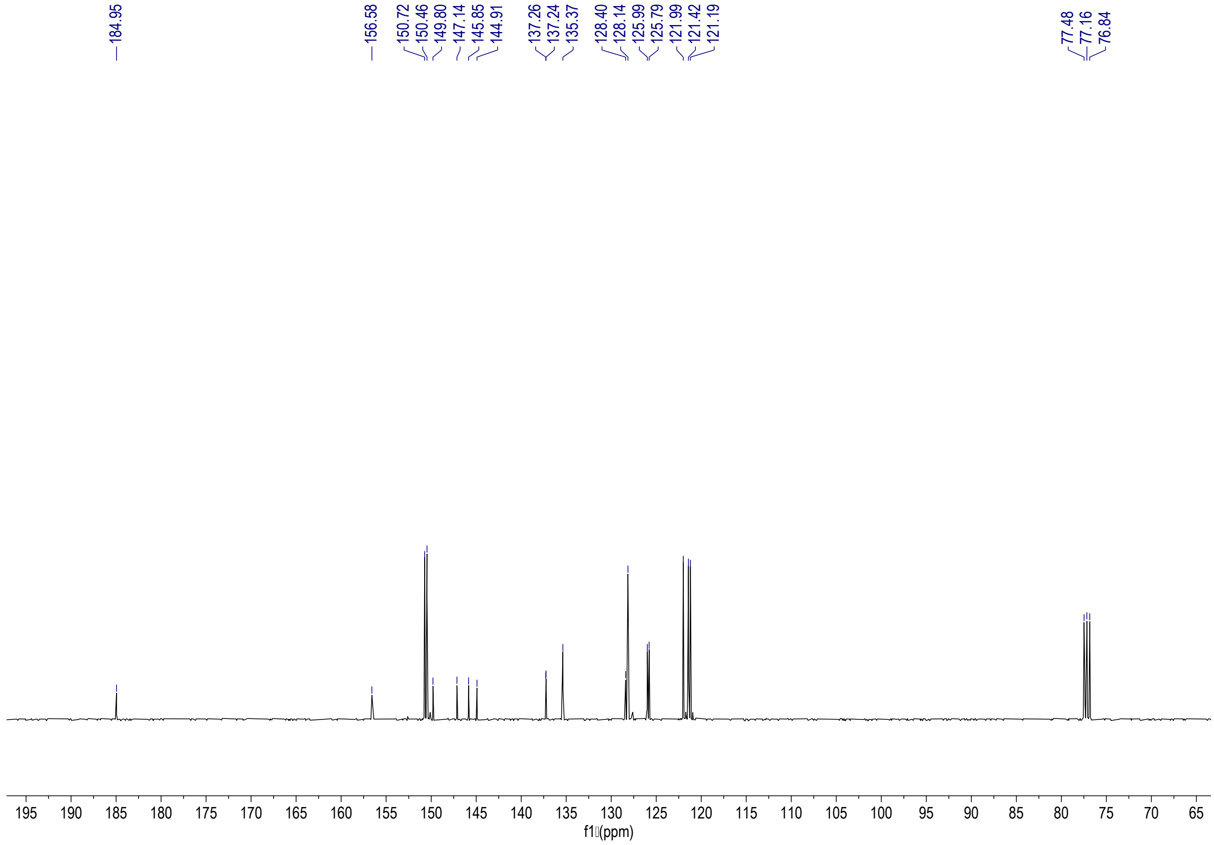


$^{13}$C-NMR spectrum of **14** (101 MHz, $CDCl_3$)

**Compound L11**

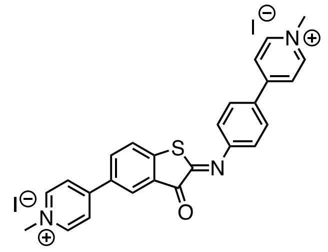

To a solution of compound **14** (70 mg, 0.18 mmol) in ACN/$CHCl_3$ (1:1, 6 mL), methyl iodide (45 µL, 0.71 mmol) was added. The solution was stirred in a sealed vessel at 50 °C overnight. The generated solid was filtered and washed with diethyl ether. Following filtration, compound **L11** (110 mg, 91%) was obtained as an orange powder. $^1$H NMR (400 MHz, DMSO-$d_6$) $\delta$ 9.05 (d, $J$ = 7.1 Hz, 2H), 9.01 (d, $J$ = 7.1 Hz, 2H), 8.65 (d, $J$ = 6.9 Hz, 2H), 8.59 (d, $J$ = 2.2 Hz, 1H), 8.55 (d, $J$ = 7.1 Hz, 2H), 8.47 (dd, $J$ = 8.4, 2.1 Hz, 1H), 8.27 (d, $J$ = 8.4 Hz, 2H), 8.02 (d, $J$ = 8.4 Hz, 1H), 7.47 (d, $J$ = 8.6 Hz, 2H), 4.35 (2s, 6H); $^{13}$C NMR (101 MHz, DMSO-$d_6$) $\delta$ 183.6, 158.6, 153.2, 152.8, 152.3, 146.9, 145.7, 145.6, 136.2, 132.3, 131.7, 129.8, 128.5, 126.7, 126.2, 124.2, 123.9, 121.1, 47.3, 47.2; ESI-HRMS $m/z$ calcd. for $C_{26}H_{21}N_3OSI$ [M]$^+$: 550.0450; found: 550.0458.

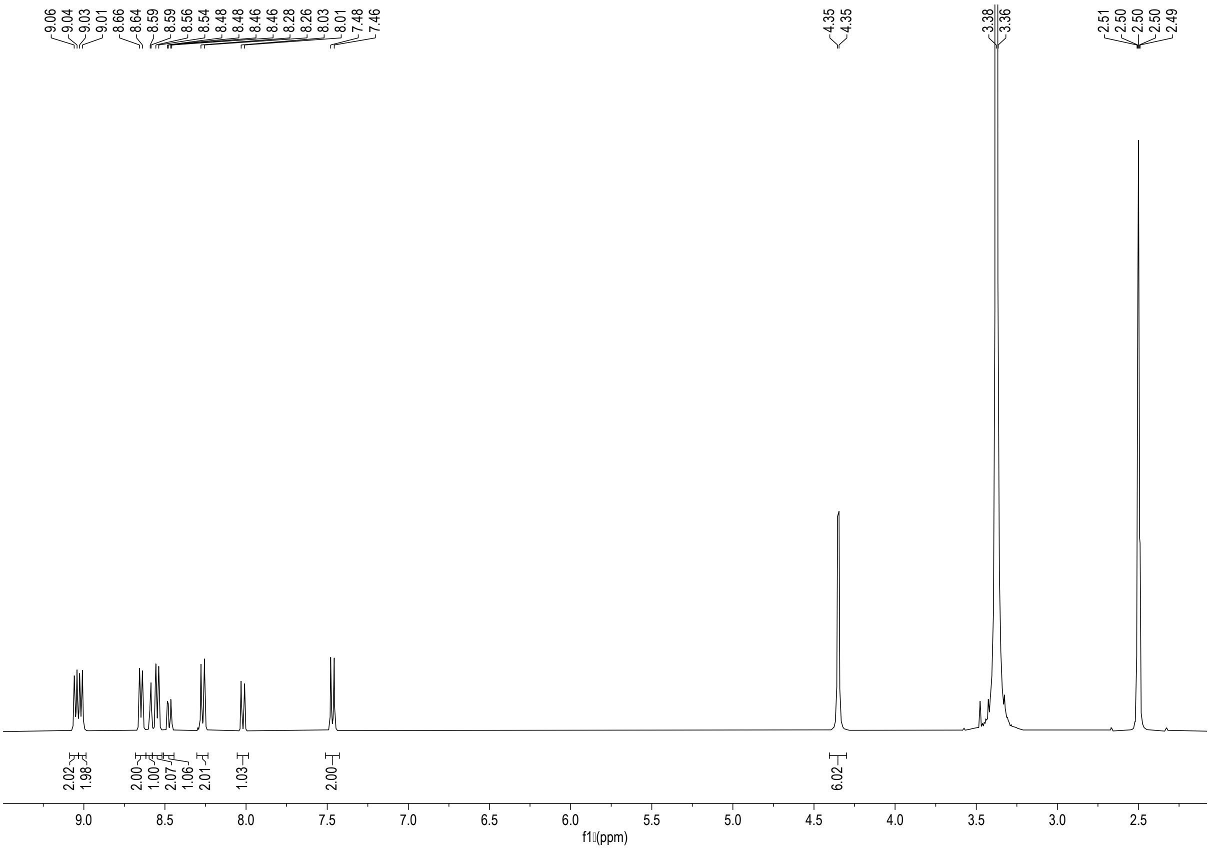

$^1$H-NMR spectrum of **L11** (400 MHz, DMSO-$d_6$)

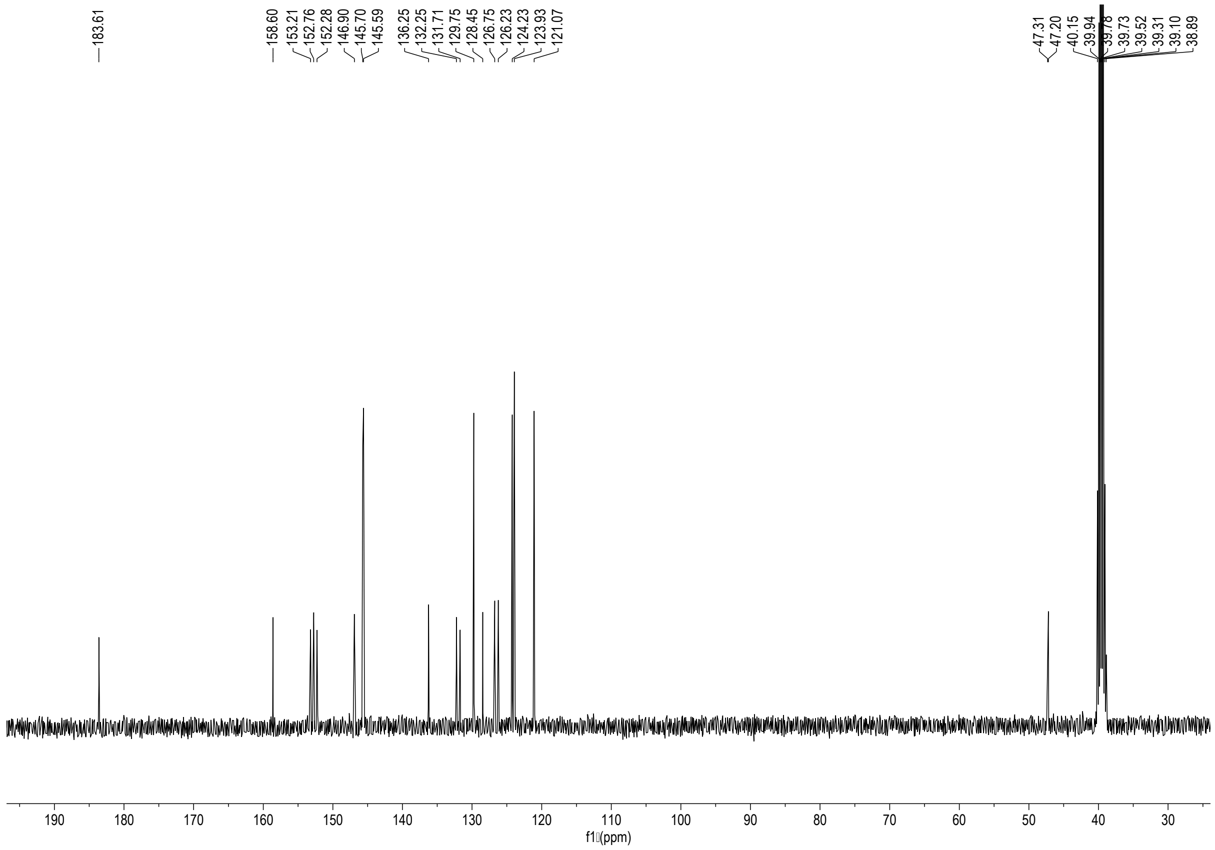


$^{13}$C-NMR spectrum of **L11** (101 MHz, DMSO-$d_6$)

### 2. Antimicrobial screening

#### Bacterial strains, growth conditions

Bacterial strains used in this study are summarised in **Table S1**. Unless otherwise specified, all microorganisms were grown on Nutrient or Luria-Bertani agar and broth (Sigma) for 18-20 h at 37°C static and 180 rpm shaking respectively.

**Table S1. Bacterial strains used in this study.**

| Strain Name | Description | Reference/Source |
|---|---|---|
| *Escherichia coli* ATCC 25922 | CLSI recommended susceptible control strain | American Type Culture Collection |
| *Escherichia coli* UTI 808 | MDR clinical UTI isolate | Gift from Dr Jacqueline Findlay |
| *Escherichia coli* IR60 | MDR clinical bloodstream isolate | Gift from Prof Tim Walsh (Oxford University) |
| *Staphylococcus aureus* strain Newman | Clinical isolate | Lorenz et al.[7] |
| *Klebsiella pneumoniae* NCTC 5055 | Reference strain | Kol et al.[8] |
| *Klebsiella pneumoniae* KP11 | Clinical isolates | Avison et al.[9] |
| *Pseudomonas aeruginosa* PAO1 | Reference strain | Holloway et al.[10] |
| *Pseudomonas aeruginosa* 4623 | Clinical isolates | Gift from Prof Matthew B Avison |
| *Acinetobacter baumannii* ATCC 19606 | Susceptible quality control strain | American Type Culture Collection |
| *Acinetobacter baumannii* 6490A | Clinical isolate | Gift from Prof Matthew B Avison |

MDR, multi-drug resistant

#### Antimicrobial susceptibility assays

Bacterial strains were tested against G4 ligands by disc susceptibility and/or broth microdilution methods as outlined by the CLSI with Mueller-Hinton agar and cation-adjusted Mueller-Hinton broth (Sigma) respectively.[11] or the disc susceptibility assays, 25 µg of each ligand was impregnated onto sterile 5 mm filter discs before placing on inoculated bacterial plates.

#### Growth curve assays

To assess whether **L5**, **L9** and **PDS** are bacteriostatic or bactericidal agents, overnight *E. coli* ATCC 25922 cultures in triplicate were diluted to a final $OD_{600}$ of 0.08-0.1 in Muller Hinton Broth 2 (Sigma-Aldrich) in flat-bottom 96-well plates (Corning) and placed in a plate reader (BMG Omega) to track absorbance readings (600 nm, every 10 min, 37°C) until the cultures entered exponential growth phase (300 min). At 300 min, 64, 4, 64 µg/mL **L5**, **L9** and **PDS** respectively were introduced to the wells and absorbance readings collected until 800 min. For determining proteomic culture conditions, cultures were prepared in 10 mL volumes in 50 mL conical flasks with 0, 32 and 64 µg/mL **PDS** and **L5** and 0, 1, 2 µg/mL **L9** and grown at 37°C, 180 RPM. Absorbance readings were measured manually on the spectrophotometer (Amersham Biosciences Ultrospec 2100 pro) every 1 h for 6-12 h and then left overnight. The last absorbance reading was taken at +24 h.

**Table S2. Minimal inhibitory concentration (MIC) values for selected candidate G4 ligands against *S. aureus* and *E. coli*.**

| | MIC (μg/mL) | | | |
|---|---|---|---|---|
| | *S. aureus* | *E. coli* | | |
| **Ligand** | **Newman** | **ATCC 25922** | **UTI 808** | **IR60** |
| L1 | 256 | >256 | >256 | >256 |
| L2 | >512 | >512 | >512 | >512 |
| L3 | 256 | 256 | 512 | 512 |
| L4 | 128 | 128 | 256 | 256 |
| L5 | 1 | 64 | 128 | 128 |
| L6 | 1 | 64 | 64 | 64 |
| L7 | 128 | >512 | >512 | >512 |
| L8 | 64 | 512 | 512 | 512 |
| L9 | 4 | 4 | 4 | 2 |
| L10 | 512 | 512 | 512 | 512 |
| L11 | 512 | 256 | 512 | 256 |
| PDS | 64 | 64 | 64 | 64 |
| TMPyP4 | 256 | 256 | 512 | 512 |
| BRACO-19 | 128 | >512 | >512 | >512 |

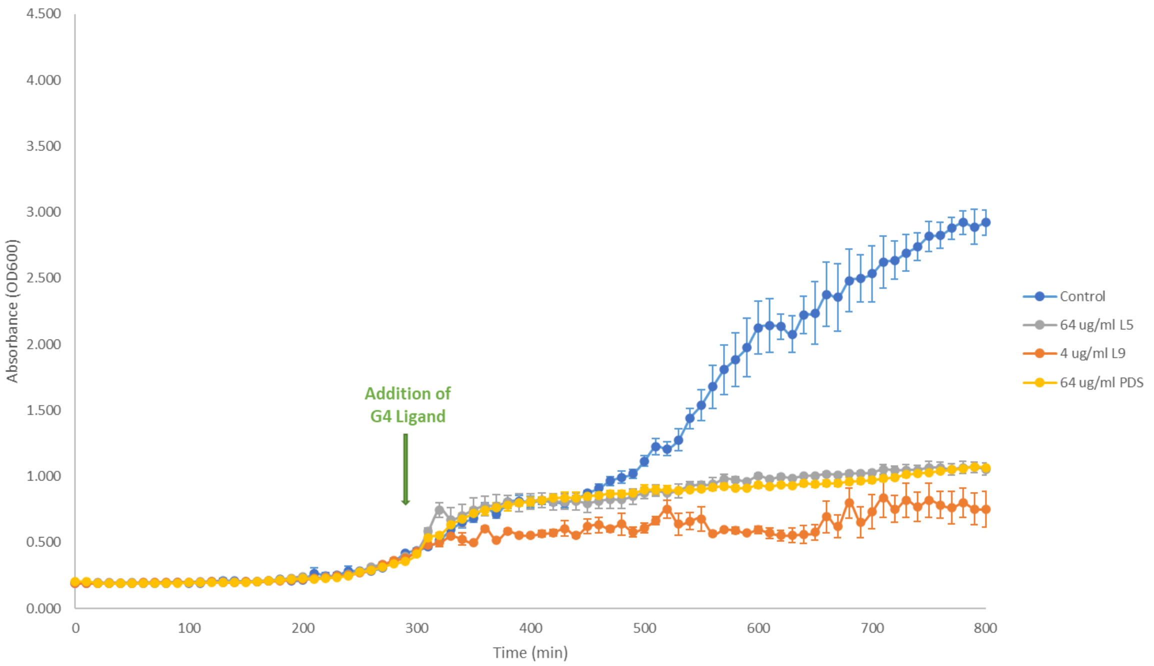


**Figure S2. Growth curves of *E. coli* ATCC25922 treated with L5, L9 and PDS at 1x MIC.** Compounds were administered 300 min after growth was initiated, data points shown are means of three replicates, error bars represent standard errors.

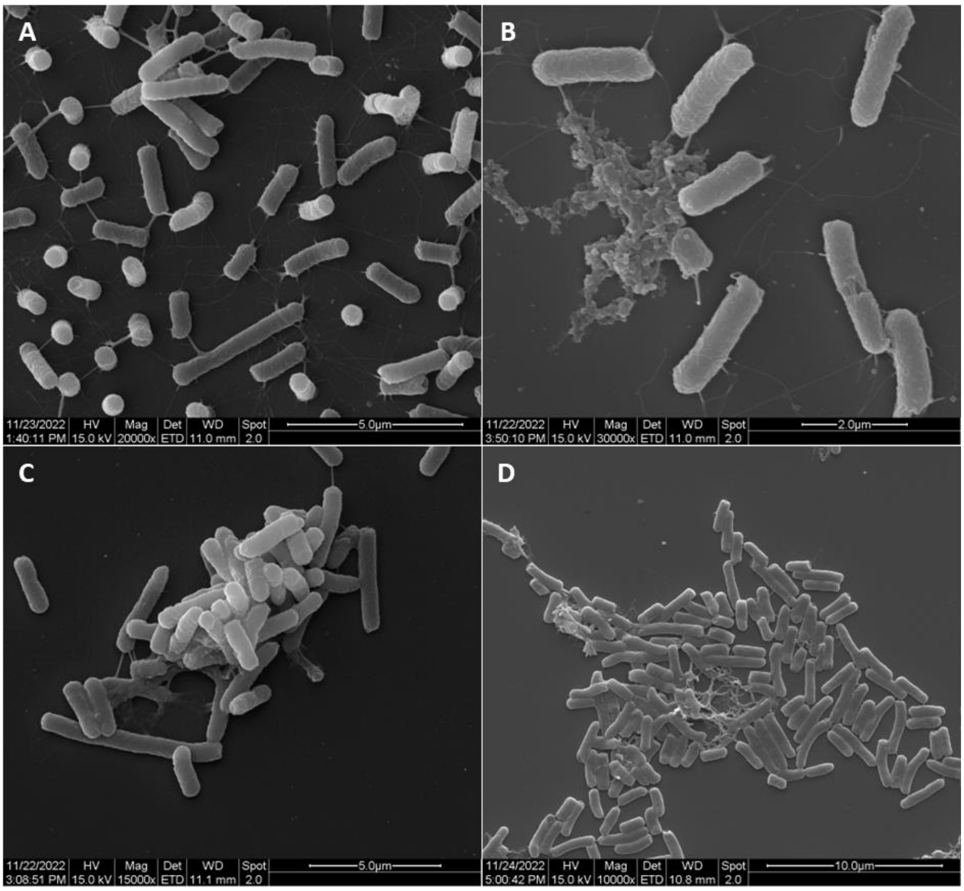


**Figure S3. Scanning Electron Micrographs of *E. coli* ATCC 25922 (A) untreated, and after 2h treatment with 1x MIC (B) L5, (C) L9 and (D) PDS for 2 hours.**

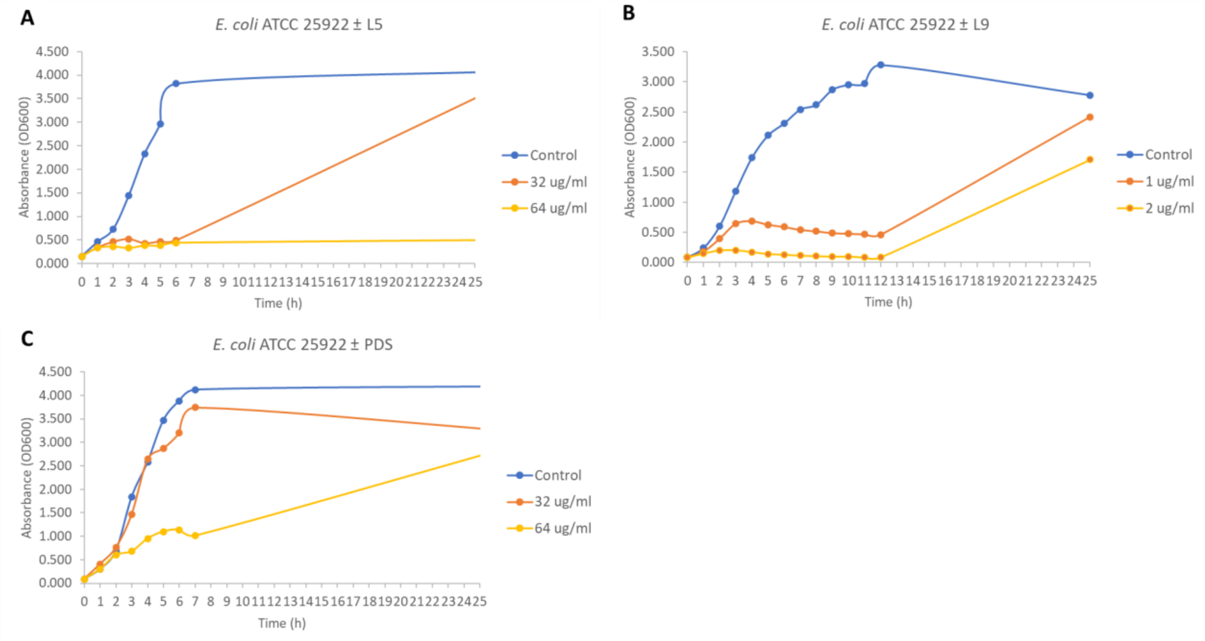


**Figure S4. Growth curves of E. coli ATCC 25922 control (non-treated in blue) and treated with various concentrations of (A) L5, (B) L9 and (C) PDS.**

### 3. Proteomic analysis of *E. coli* treated with L5, L9 and PDS

#### Culture preparation, 6-plex TMT-labelling and LC-MS/MS

Whole cell lysates were obtained from *E. coli* ATCC 25922 cultures grown in Mueller Hinton Broth 2 in triplicate for 7 h, with or without 64, 1 and 64 µg/mL **L5**, **L9** and **PDS** respectively. Aliquots of 50 µg of each sample were digested with trypsin (2.5 µg trypsin per 100 µg protein; 37°C, overnight), labelled with Tandem Mass Tag (TMT) 6-plex reagents according to the manufacturer's protocol (Thermo Fisher Scientific) and the labelled samples pooled.

The pooled sample was evaporated to dryness, resuspended in 5% formic acid and then desalted using a SepPak cartridge according to the manufacturer's instructions (Waters, Milford, Massachusetts, USA). Eluate from the SepPak cartridge was again evaporated to dryness and resuspended in buffer A (20 mM ammonium hydroxide, pH 10) prior to fractionation by high pH reversed-phase chromatography using an Ultimate 3000 liquid chromatography system (Thermo Scientific). In brief, the sample was loaded onto an XBridge BEH C18 Column (130Å, 3.5 µm, 2.1 mm X 150 mm, Waters, UK) in buffer A and peptides eluted with an increasing gradient of buffer B (20 mM Ammonium Hydroxide in acetonitrile, pH 10) from 0-95% over 60 minutes. The resulting fractions were evaporated to dryness and resuspended in 1% formic acid prior to analysis by nano-LC MSMS using an Orbitrap Fusion Lumos mass spectrometer (Thermo Scientific).

#### Nano-LC Mass Spectrometry

High pH RP fractions were further fractionated using an Ultimate 3000 nano-LC system in line with an Orbitrap Fusion Lumos mass spectrometer (Thermo Scientific). In brief, peptides in 1% (vol/vol) formic acid were injected onto an Acclaim PepMap C18 nano-trap column (Thermo Scientific). After washing with 0.5% (vol/vol) acetonitrile 0.1% (vol/vol) formic acid peptides were resolved on a 250 mm × 75 µm Acclaim PepMap C18 reverse phase analytical column (Thermo Scientific) over a 150 min acetonitrile gradient, divided into 7 gradient segments (1-6% solvent B over 1 min., 6 - 15% B over 58 min., 15 - 32% B over 58 min., 32 - 40% B over 5 min., 40 - 90% B over 1 min., held at 90% B for 6 min and then reduced to 1% B over 1 min.) with a flow rate of 300 nL $min^{-1}$. Solvent A was 0.1% formic acid and Solvent B was aqueous 80% acetonitrile in 0.1% formic acid. Peptides were ionized by nano-electrospray ionization at 2.0kV using a stainless steel emitter with an internal diameter of 30 µm (Thermo Scientific) and a capillary temperature of 275°C.

All spectra were acquired using an Orbitrap Fusion Lumos mass spectrometer controlled by Xcalibur 3.0 software (Thermo Scientific) and operated in data-dependent acquisition mode using an SPS-MS3 workflow. FTMS1 spectra were collected at a resolution of 120 000, with an automatic gain control (AGC) target of 200 000 and a max injection time of 50ms. Precursors were filtered with an intensity threshold of 5000, according to charge state (to include charge states 2-7) and with monoisotopic peak determination set to Peptide. Previously interrogated precursors were excluded using a dynamic window (60 s +/- 10 ppm). The MS2 precursors were isolated with a quadrupole isolation window of 0.7m/z. ITMS2 spectra were collected with an AGC target of 10 000, max injection time of 70ms and CID collision energy of 35%.

For FTMS3 analysis, the Orbitrap was operated at 50 000 resolution with an AGC target of 50 000 and a max injection time of 105 ms. Precursors were fragmented by high energy collision dissociation (HCD) at a normalised collision energy of 60% to ensure maximal TMT reporter ion yield. Synchronous Precursor Selection (SPS) was enabled to include up to 5 MS2 fragment ions in the FTMS3 scan.

#### Proteomic analysis of G4-associated and essential proteins for cell growth

Genomic coordinates from the Marsico *et al.*[12] study were used to identify the genes carrying observed G4 sequences (OQs) in the sense and antisense DNA strands of *E. coli* ATCC 25922. An R script was written to extract the OQ sequences and associated genes from NCBI GenBank. G4 sites that were > 3,000 bp upstream any ORFs were classified as non-consequential to gene expression and removed from further analysis. A list of the genes associated with RNA G4 sequences identified by Shao *et al*.[13] was taken from their Table S1A. All G4-associated genes (hereby referred to as G4 genes) identified by these two studies were collated and duplicates removed. The final list consisted of 404 G4 genes for further analysis. Analysis on essential proteins for cell growth was conducted based on the essential genes listed in the Profiling of *E. coli* Chromosome (PEC) database (as of 24 Feb 2021) (https://shigen.nig.ac.jp/ecoli/pec/index.jsp).

**PANTHER, STRING and G4Hunter analysis**

PANTHER 18.0 was used to classify differentially expressed proteins into protein classes.[14] STRING Version 11.5 (https://string-db.org/) was used to study the protein-protein interaction networks of the proteins with changes in abundance due to G4 ligand treatment against *E. coli* CFT073.[15] The highest confidence score of 0.900 was applied to the analysis. G4Hunter (https://bioinformatics.cruk.cam.ac.uk/G4Hunter/) was used to score the propensity of G4-structures to form from the sequences of interest.[16]

**Table S3. Summary of 43 essential genes listed in the Profiling of *E. coli* Chromosome (PEC) database and associated with G4 motifs.**

| | | LogFC (p<0.05) | | |
|---|---|---|---|---|
| **PEC** | **Protein Description** | **L5** | **L9** | **PDS** |
| *nadD* | nicotinic acid mononucleotide adenylyltransferase, NAD(P)-dependent | | | 1.575 |
| *ftsW* | lipid II flippase; integral membrane protein involved in stabilizing FstZ ring during cell division | | | 0.937 |
| *ftsZ* | GTP-binding tubulin-like cell division protein | | | -0.276 |
| *mukE* | Chromosome condensin MukBEF, MukE localization factor | | 1.582 | 0.232 |
| *gyrA* | DNA gyrase (type II topoisomerase), subunit A | | 1.063 | 0.690 |
| *hemB* | 5-aminolevulinate dehydratase (porphobilinogen synthase) | | 1.038 | 0.494 |
| *pth* | peptidyl-tRNA hydrolase | | 1.038 | 0.561 |
| *glyS* | glycine tRNA synthetase, beta subunit | | 0.877 | 0.474 |
| *topA* | DNA topoisomerase I, omega subunit | | 0.863 | 0.312 |
| *mreB* | cell wall structural complex MreBCD, actin-like component MreB | | 0.641 | 1.673 |
| *zipA* | FtsZ stabilizer | | 0.466 | 0.745 |
| *mrdB* | cell wall shape-determining protein | | 0.255 | |
| *glmU* | fused N-acetyl glucosamine-1-phosphateuridyltransferase/glucosamine-1-phosphate acetyl transferase | | 0.195 | 0.531 |
| *ligA* | DNA ligase, NAD(+)-dependent | | 0.143 | |
| *efp* | polyproline-specific translation elongation factor EF-P | | -0.152 | 1.037 |
| *prfB* | peptide chain release factor RF-2 | 1.037 | 1.200 | 1.115 |
| *thrS* | threonyl-tRNA synthetase | 0.962 | 0.265 | 0.628 |
| *rho* | transcription termination factor | 0.899 | 0.362 | 0.899 |
| *nusA* | transcription termination/antitermination L factor | 0.727 | 0.681 | |
| *lnt* | apolipoprotein N-acyltransferase | 0.696 | 0.712 | |
| *mnmA* | tRNA(Gln,Lys,Glu) U34 2-thiouridylase, first step in mnm(5)-s(2)U34-tRNA synthesis | 0.638 | 1.798 | 0.719 |
| *suhB* | inositol monophosphatase | 0.564 | 0.989 | 0.910 |
| *yidC* | membrane protein insertase | 0.342 | 1.551 | 0.696 |
| *murC* | UDP-N-acetylmuramate:L-alanine ligase | 0.299 | 0.419 | 0.386 |
| *hemL* | glutamate-1-semialdehyde aminotransferase (aminomutase) | 0.283 | 0.443 | 1.744 |
| *rpoB* | RNA polymerase, beta subunit | 0.277 | 0.380 | 0.727 |
| *valS* | valyl-tRNA synthetase | -0.171 | 0.113 | 0.687 |
| *asnS* | asparaginyl tRNA synthetase | -0.188 | 1.622 | |
| *leuS* | leucyl-tRNA synthetase | -0.226 | 1.033 | 0.593 |
| *groL* | Cpn60 chaperonin GroEL, large subunit of GroESL | -0.325 | -0.705 | -0.631 |
| *bamA* | outer membrane protein assembly factor, forms pores; required for OM biogenesis; in BamABCDE OM protein complex | -0.429 | | -0.395 |
| *gyrB* | DNA gyrase, subunit B | -0.477 | 1.015 | 0.563 |
| *pyrG* | CTP synthetase | -0.685 | 1.089 | 0.892 |
| *murD* | UDP-N-acetylmuramoyl-L-alanine:D-glutamate ligase | -0.937 | -0.223 | |
| *tyrS* | tyrosyl-tRNA synthetase | -1.021 | 0.900 | |
| *accC* | acetyl-CoA carboxylase, biotin carboxylase subunit | -1.202 | 0.835 | 0.558 |
| *prmC* | N5-glutamine methyltransferase, modifies release factors RF-1 and RF-2 | -1.242 | | |
| *ispH* | 4-hydroxy-3-methylbut-2-enyl diphosphate reductase, 4Fe-4S protein | -1.429 | 0.557 | -0.179 |
| *folC* | bifunctional folylpolyglutamate synthase/ dihydrofolate synthase | -1.508 | 0.586 | |
| *alaS* | alanyl-tRNA synthetase | | | |
| *dnaA* | chromosomal replication initiator protein DnaA, DNA-binding transcriptional dual regulator | | | |
| *murF* | UDP-N-acetylmuramoyl-tripeptide:D-alanyl-D- alanine ligase | | | |
| *rnpB* | RNase P, M1 RNA component | | | |

Heatmap shading: red indicate increases in protein abundance, blue indicate decreases compared to non-treated control, white indicate no proteins detected.

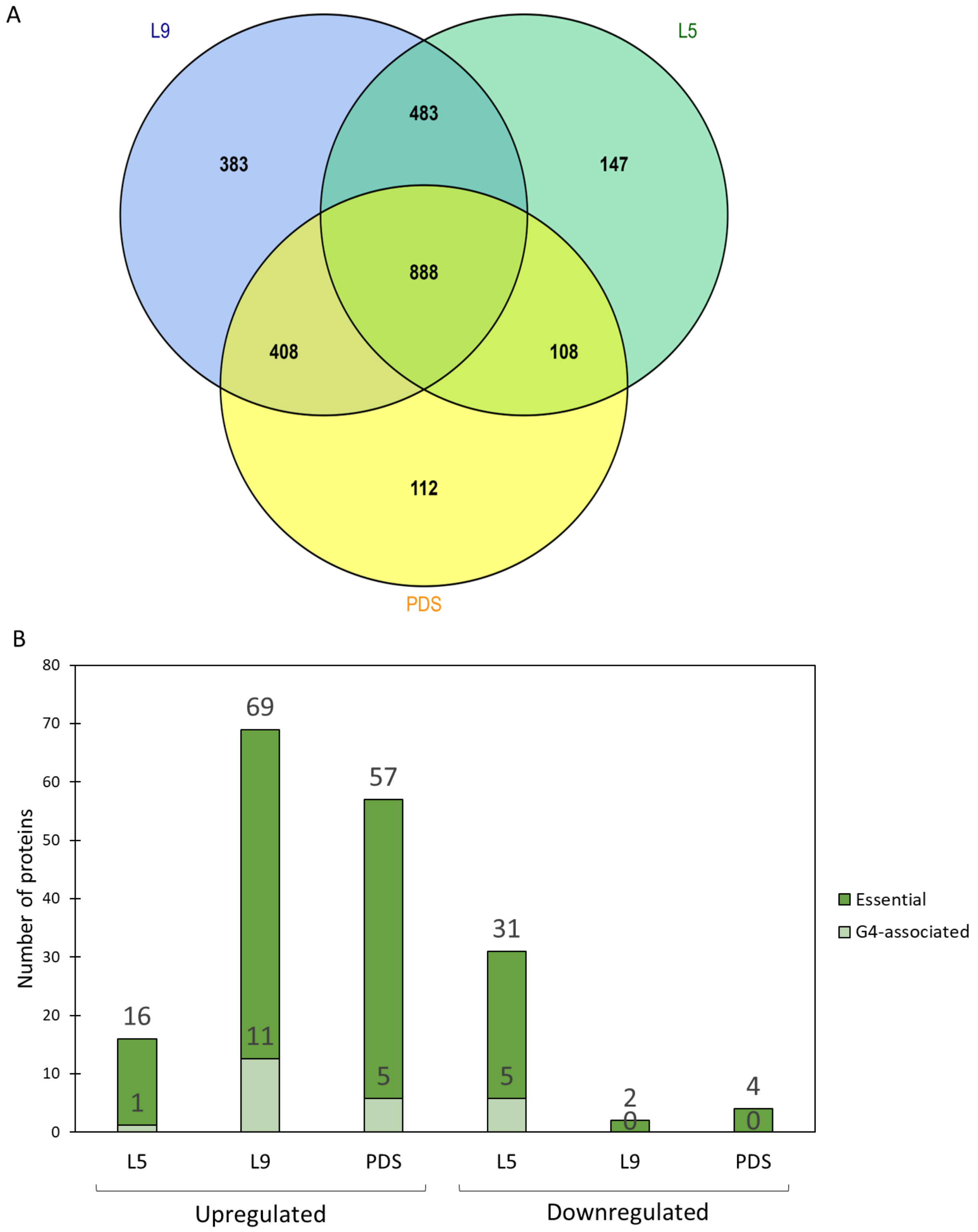


**Figure S5. Summary of (A) total number of proteins with changes in expression levels compared to untreated controls (p ≤ 0.05) and (B) significant essential PEC-associated proteins of which are G4-associated (p ≤ 0.05, (-1 ≤ Log2FC ≥ 1) from total cell proteomes of *E. coli* cultures treated with L5, L9 and PDS.**

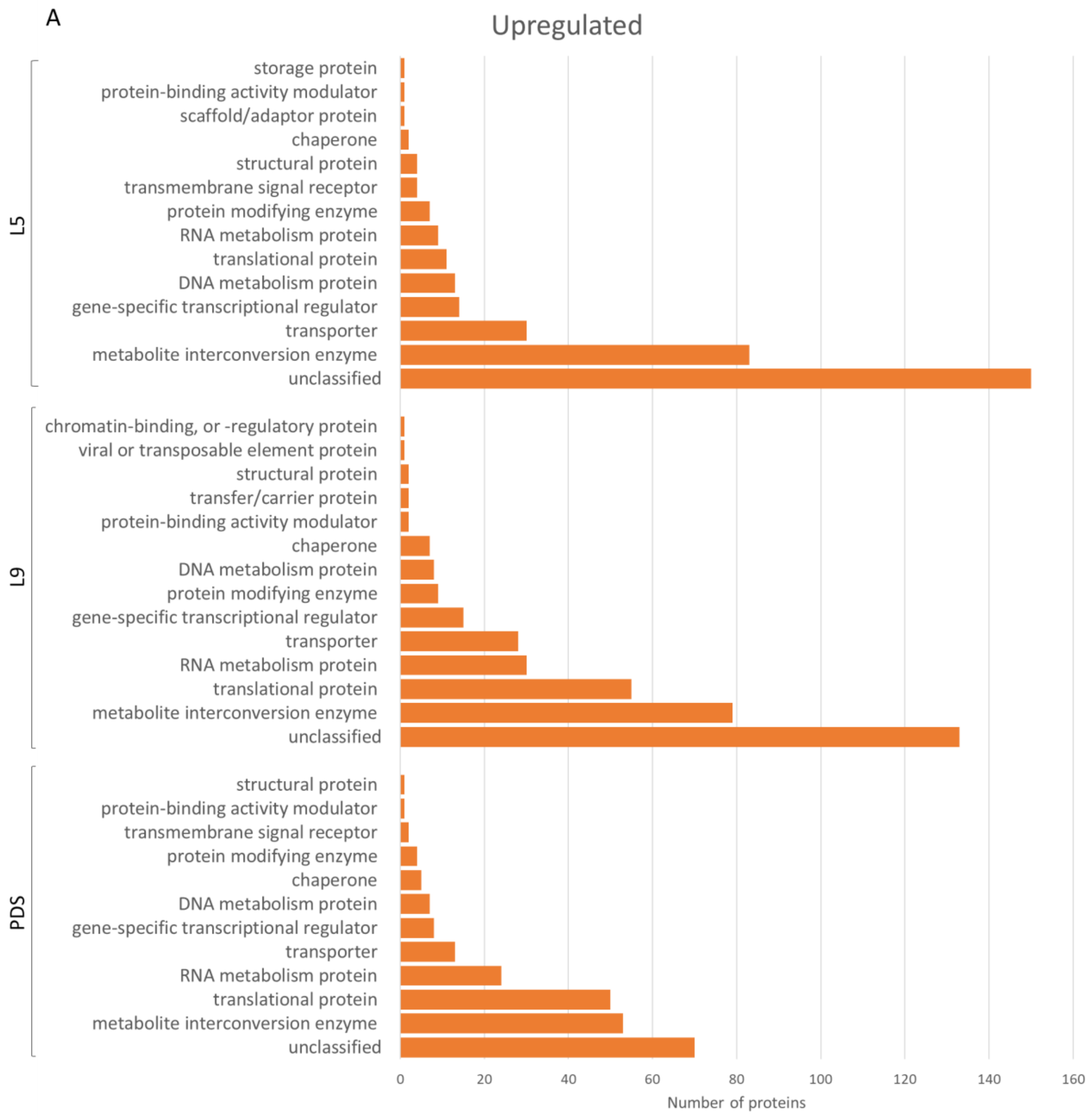
A
Upregulated
L5
storage protein
protein-binding activity modulator
scaffold/adaptor protein
chaperone
structural protein
transmembrane signal receptor
protein modifying enzyme
RNA metabolism protein
translational protein
DNA metabolism protein
gene-specific transcriptional regulator
transporter
metabolite interconversion enzyme
unclassified
L9
chromatin-binding, or -regulatory protein
viral or transposable element protein
structural protein
transfer/carrier protein
protein-binding activity modulator
chaperone
DNA metabolism protein
protein modifying enzyme
gene-specific transcriptional regulator
transporter
RNA metabolism protein
translational protein
metabolite interconversion enzyme
unclassified
PDS
structural protein
protein-binding activity modulator
transmembrane signal receptor
protein modifying enzyme
chaperone
DNA metabolism protein
gene-specific transcriptional regulator
transporter
RNA metabolism protein
translational protein
metabolite interconversion enzyme
unclassified
0
20
40
60
80
100
120
140
160
Number of proteins

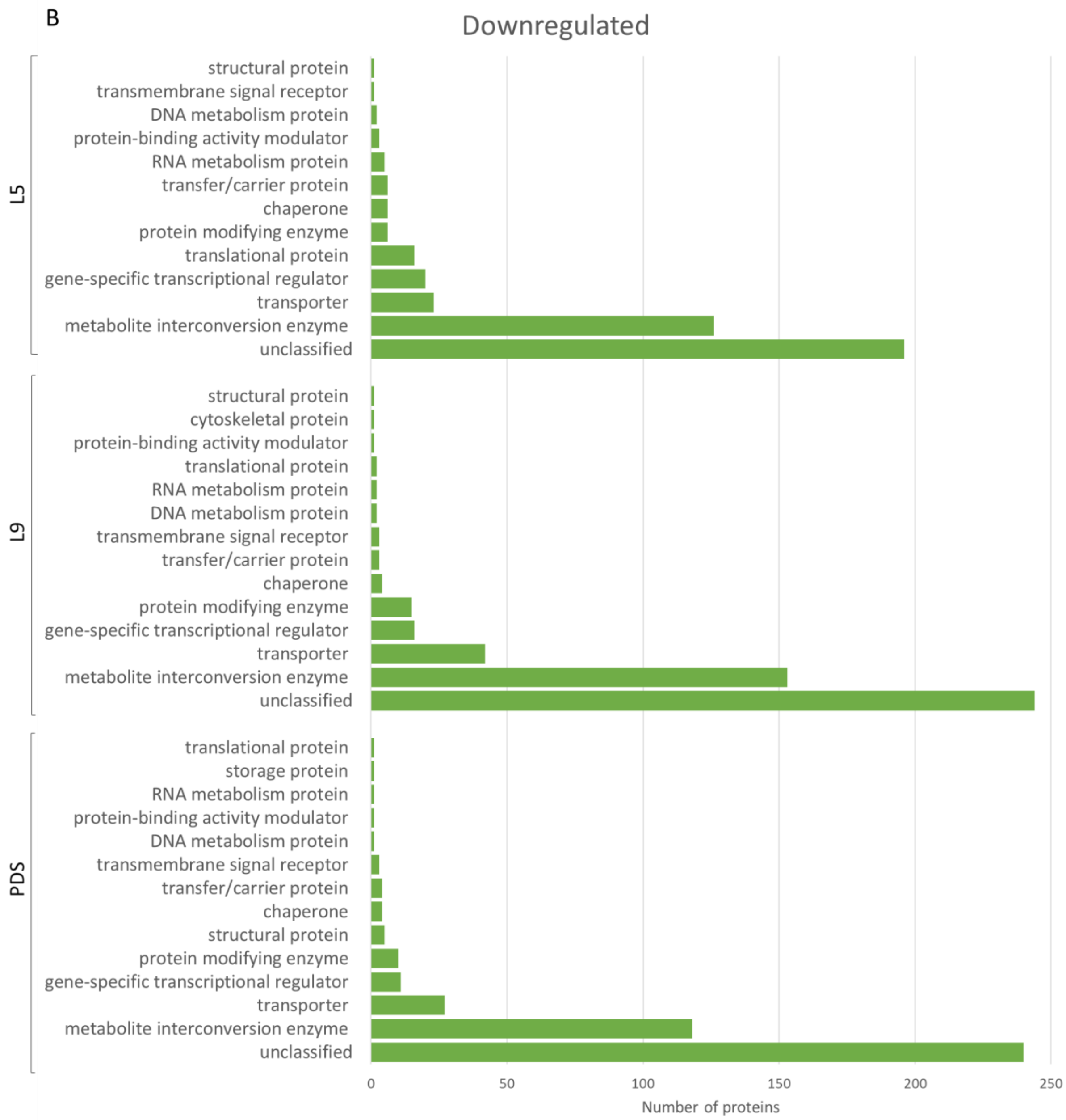


**Figure S6. Summary of differentially expressed (A) upregulated and (B) downregulated protein classes classified by PANTHER.**

### 4. Bioinformatic analysis of OQ-associated proteins of interest to shortlist candidate G4 sequences for biophysical investigation

*Escherichia coli* K-12 MG1655 genome (NC_000913.3 /GCF_000005845.2) was downloaded from the NCBI webpage on July 2022 and analyzed with G4-iM Grinder to locate and characterize Potential G-Quadruplex Sequences (PQSs)[17].G4-iM Grinder is an R-based algorithm that locates, quantifies, and qualifies PQS, potential i-Motifs and their higher-order versions in RNA and DNA genomes. The definitions of a G-quadruplex were set to a LAX parameter configuration as described in Belmonte-Reche *et al.*[18]; the difference with the package's predefined parameters being that it will accept G-runs of size 2 (compared to a minimum size of 3 for the predefined configuration), loops lengths of maximum 20 (compared to a maximum of 10 for the predefined configuration), a maximum total PQS length of 50 (compared to a maximum of 33 for the predefined configuration) and a total maximum of 1 bulge per PQS (compared to 3 total acceptable bulges for the predefined configuration). The "size-restricted overlapping search and frequency count" method (Method 2, M2A and M2B) was then used to locate all the potential candidates. To determine PQS G-quadruplex probability formation, the average of G4Hunter[19] and PQSfinder[20] algorithms were calculated and used as threshold. G-quadruplex probability scores equal or higher than 40 were considered to have a HIGH probability of being able to form G-quadruplexes *in vitro.* PQS with scores equal or higher than 20 but lower than 40 were considered to have a MEDIUM probability, whilst PQSs with scores lower than 20 were considered to have a LOW probability of forming G-quadruplexes. PQS densities were calculated as [($10^5$ × PQS counts)/(genome or gene length)] to compare between different-sized genes and genomes.

**Method**

We performed a broad *in silico* analysis on the genome of Escherichia coli K-12 MG1655 to locate, identify and characterize PQSs with the R-package G4-iM Grinder[17]. We employed a LAX configuration ruleset as G-quadruplex definitions[18] to detect G-runs which associate with at least 3 others and that can ultimately form a PQS. This parameter configuration allowed for the identification of sequences with G-runs of size 2, longer loops, longer overall PQS size and maximum one bulge per candidate. Accepting these features is usually considered to lower the stability of G-quadruplexes compared to the predefined parameters, but not necessarily prevent their formation. Furthermore, G-quadruplexes composed of sized-2 G-runs[21] and with bulges[22] have been mainly studied in microbiological genomes due to their overall smaller genomic sizes and frequently lower G content when compared to their human counterpart (as these genomic features lower the probability of finding traditional PQSs that abide the $[GGG(X_{1-7}GGG)_3]$ folding rule[23]. We evaluated all our results with G4Hunter[19]) and PQSfinder algorithms to discriminate PQSs by their probability of forming G-quadruplexes *in vitro.* These algorithms evaluate different PQS features that affect G4-formation and return a score value that denotes more likelihood of G-quadruplex formation as it increases. On the one side, PQSfinder considers the size of G-runs, the bulges in the G-runs and the length of loops in its calculus, whilst on the other side, G4Hunter evaluates the PQSs' G richness and C skewness. With this information at hand, our attention for further *in vitro* evaluation and testing was set on the candidates with the highest scores located on genes of our interest.

**Table S4. Bioinformatic analysis of OQ-associated proteins of interest.**

| | | length | G % | PQS counts | | | | PQS density (per $10^5$ nt) | | | |
|---|---|---|---|---|---|---|---|---|---|---|---|
| | | | | ALL | Score ≥ 20 | Score ≥ 30 | Score ≥ 40 | ALL | Score ≥ 20 | Score ≥ 30 | Score ≥ 40 |
| | complete genome | 9283304 | 25 | 456955 | 198423 | 48915 | 6285 | 4922 | 2137 | 527 | 68 |
| genes | aroL | 1050 | 28 | 33 | 18 | 9 | 1 | 3143 | 1714 | 857 | 95 |
| genes | asnS | 2802 | 27 | 127 | 64 | 3 | 0 | 4532 | 2284 | 107 | 0 |
| genes | clpB | 5148 | 28 | 260 | 122 | 36 | 1 | 5051 | 2370 | 699 | 19 |
| genes | cobS | 1488 | 31 | 108 | 67 | 15 | 1 | 7258 | 4503 | 1008 | 67 |
| genes | creA | 948 | 27 | 37 | 27 | 20 | 10 | 3903 | 2848 | 2110 | 1055 |
| genes | dppA | 3216 | 26 | 145 | 75 | 23 | 2 | 4509 | 2332 | 715 | 62 |
| genes | folC | 2538 | 31 | 135 | 49 | 15 | 2 | 5319 | 1931 | 591 | 79 |
| genes | glnD | 5346 | 26 | 305 | 162 | 52 | 18 | 5705 | 3030 | 973 | 337 |
| genes | hemL | 2562 | 30 | 211 | 71 | 16 | 1 | 8236 | 2771 | 625 | 39 |
| genes | ispH | 1902 | 31 | 113 | 70 | 12 | 1 | 5941 | 3680 | 631 | 53 |
| genes | maeA | 3396 | 26 | 185 | 65 | 25 | 4 | 5448 | 1914 | 736 | 118 |
| genes | mnma | 2214 | 28 | 124 | 52 | 6 | 0 | 5601 | 2349 | 271 | 0 |
| genes | mtgA | 1458 | 29 | 62 | 34 | 15 | 3 | 4252 | 2332 | 1029 | 206 |
| genes | mukE | 1410 | 27 | 44 | 15 | 6 | 1 | 3121 | 1064 | 426 | 71 |
| genes | pdxA | 1980 | 28 | 151 | 77 | 33 | 15 | 7626 | 3889 | 1667 | 758 |
| genes | thrA | 4926 | 28 | 319 | 176 | 64 | 26 | 6476 | 3573 | 1299 | 528 |
| genes | tyrS | 2550 | 28 | 126 | 65 | 7 | 0 | 4941 | 2549 | 275 | 0 |
| genes | ybiO | 4452 | 27 | 265 | 136 | 39 | 1 | 5952 | 3055 | 876 | 22 |
| genes | yfiC | 1476 | 30 | 95 | 66 | 30 | 2 | 6436 | 4472 | 2033 | 136 |
| genes | yhaJ | 1794 | 31 | 121 | 87 | 48 | 24 | 6745 | 4849 | 2676 | 1338 |
| genes | yidC | 3294 | 26 | 184 | 67 | 6 | 0 | 5586 | 2034 | 182 | 0 |
| genes | yjbD | 546 | 26 | 8 | 3 | 0 | 0 | 1465 | 549 | 0 | 0 |

**Table S5. G4 DNA sequences selected from bioinformatic and proteomics analysis.**

| Oligo Name | Sequence |
|---|---|
| hemL | 5'-FAM-GGT-CCG-GTC-TAT-CAG-GCG-GGT-TAMRA-3' |
| dppA | 5'-FAM-GGG-CTG-GAC-TGG-CGA-TAA-CGG-GG-TAMRA-3' |
| clpB | 5'-FAM-GGC-GCG-TTG-GAC-GGG-G-TAMRA-3' |
| yjbD | 5'-FAM-GGG-AAA-AGG-GTT-AGG-GTG-AGG-G-TAMRA-3' |
| thrA | 5'-FAM-GGG-CGA-TGG-GGG-TAA-TGG-TGC-GGG-GG-TAMRA-3' |
| pdxA | 5'-FAM-GGG-GAG-TTG-GGG-GAA-TAA-GGG-CGG-AGG-G-TAMRA-3' |
| glnD | 5'-FAM-GCG-GTT-GAC-CGG-GCA-GGG-TGG-G-TAMRA-3' |
| asnS | 5'-FAM-GGG-TAA-AGT-CGT-GGC-GTC-GCC-GGG-CCA-GGG-G-TAMRA-3' |
| yhaJ | 5'-FAM-GGG-GGC-GTG-GGA-ACG-GCT-GGA-GCA-GGG-G-TAMRA-3' |
| ybiO | 5'-FAM-GGG-TAC-GCG-TGC-GGG-CGC-TGG-GTA-GCG-GG-TAMRA-3' |
| aroL | 5'-FAM-GGA-GAT-CGT-CGA-AAG-GGA-AGA-GTG-GGC-GGG-TAMRA-3' |
| cobS | 5'-FAM-GGG-CGG-GCA-AAC-GGG-CGA-TAC-GCT-GGG-TAMRA-3' |
| mtgA | 5'-FAM-GGG-ATG-GGC-GTA-GCT-GGG-TTC-GAA-AAG-GG-TAMRA-3' |
| maeA | 5'-FAM-GGG-GCT-TGG-TGA-CCA-GGG-CAT-CGG-CGG-G-TAMRA-3' |
| yfiC | 5'-FAM-GGG-CGC-ATG-GGC-ACC-GGT-GGC-TGG-GG-TAMRA-3' |
| creA | 5'-FAM-GGT-GGT-ATT-AAA-GGG-GGA-TTG-GGT-CTG-GCG-TAMRA-3' |
| mukE | 5'-FAM-GGA-ACG-GCT-GGC-GAA-TGA-GGG-G-TAMRA-3' |

### 5. Circular Dichroism (CD) analysis of G4 DNA candidate sequences identified by bioinformatics

To determine whether the DNA sequence selected from the bioinformatics and proteomic analysis, Circular dichroism (CD) experiments with candidate G4 DNA samples were conducted using a spectropolarimeter (J-810, Jasco, Japan) fitted with a Peltier temperature controller. Measurements were taken in a quartz cuvette with a path length of 5 mm at 20 °C, at a 1000 nm/min scanning speed at 1 nm intervals with a 1 nm bandwidth. The DNA concentration was 5 µM in 0-100 mM potassium phosphate buffer, pH 7.4. The CD spectra were recorded over the range 550 – 200 nm and baseline corrected for the buffer used. During the titrations, aliquots of ligand were added from a 1 mM stock solution in buffer (containing 10% DMSO for solubility). The sample was mixed thoroughly, and the CD spectrum acquired immediately. The reported spectrum for each titration point represents the average of 3 scans. Data processing was carried out using Prism 7. Observed ellipticities were converted to molar ellipticity.

**Parallel**

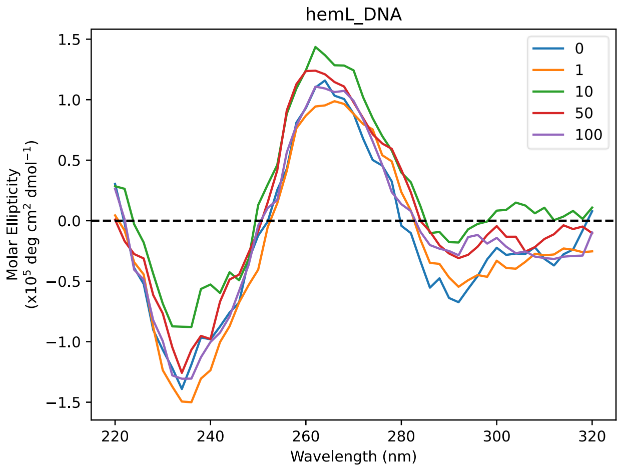


**Hybrid → Parallel**

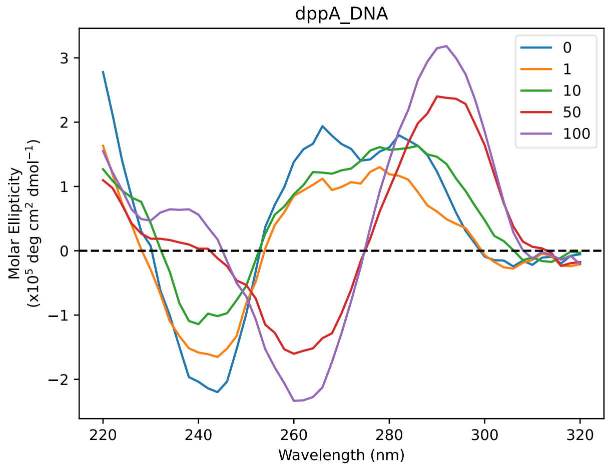


**Parallel**

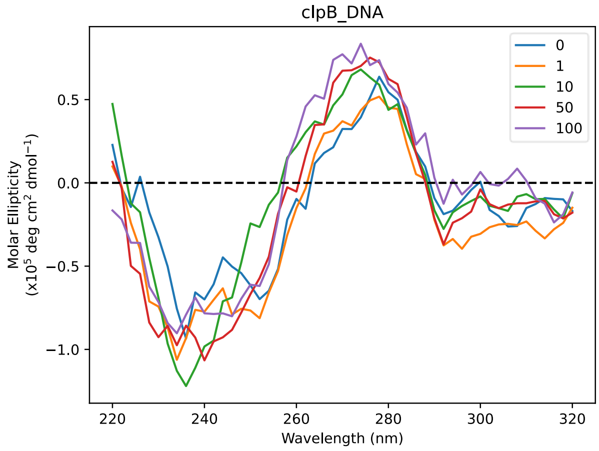


**Parallel → Hybrid**

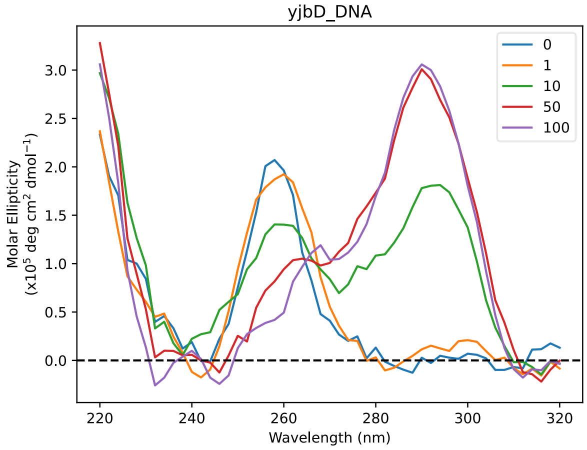

**Parallel + shoulder**

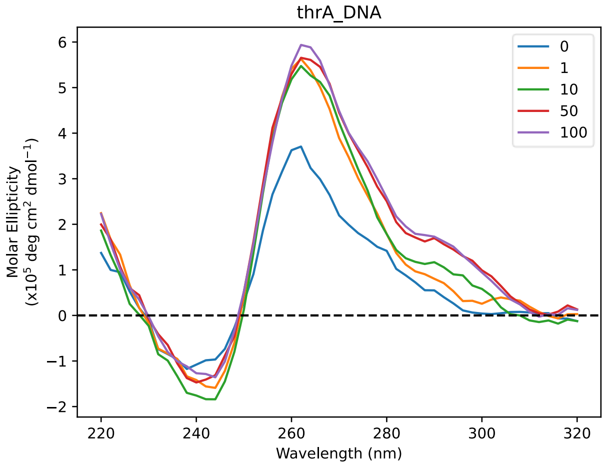


**Parallel + shoulder**

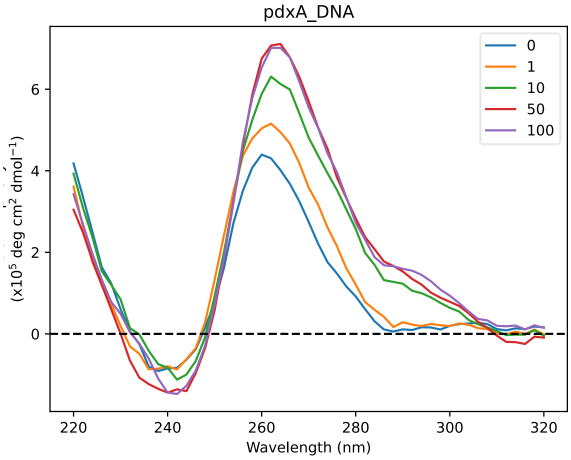


**Parallel + shoulder**

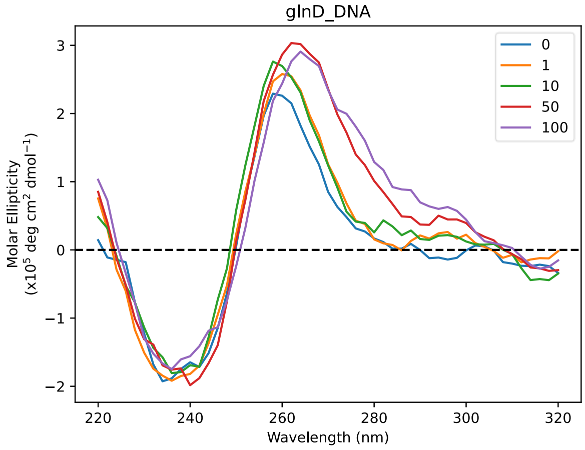


**Parallel + shoulder**

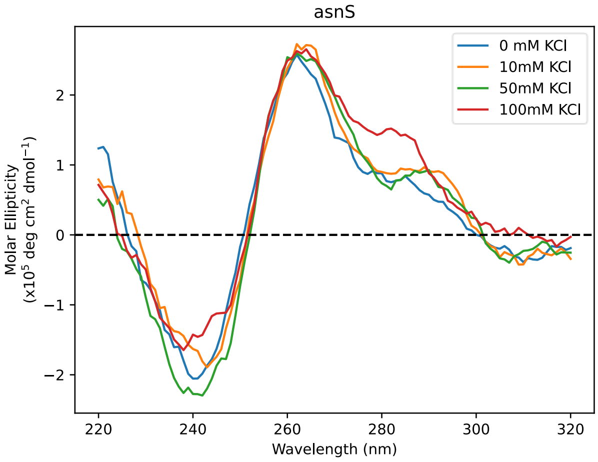


**Hybrid**

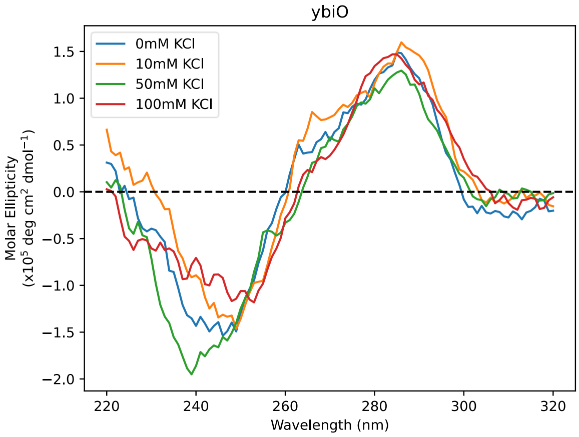


**Parallel**

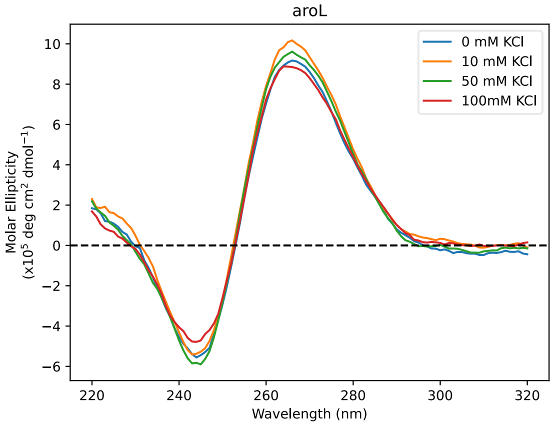

### Parallel + shoulder

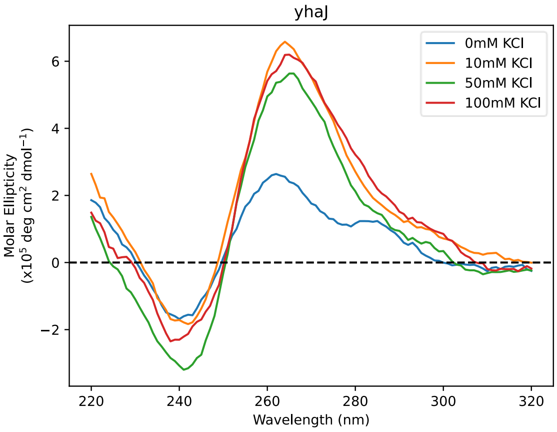


### Mix hybrid - Parallel

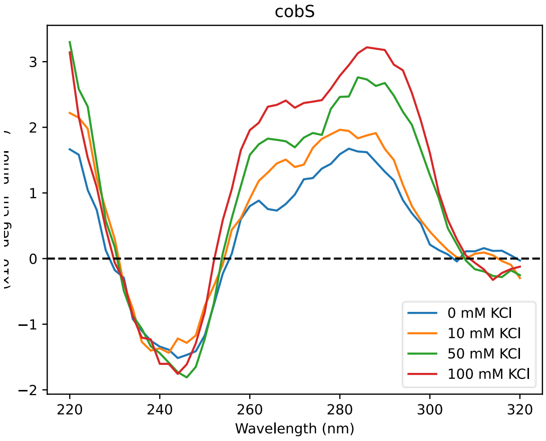


### Mix Hybrid - Parallel

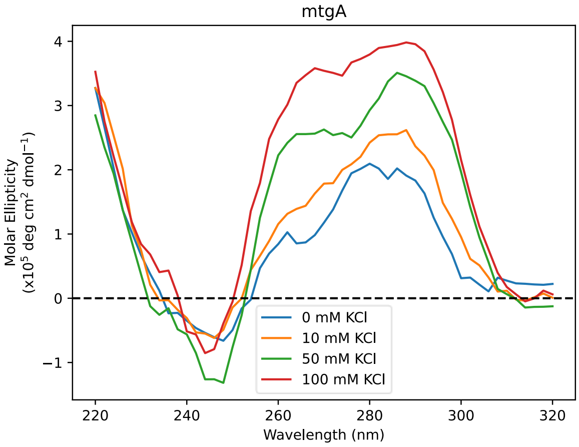


### Mix Hybrid - Antiparallel

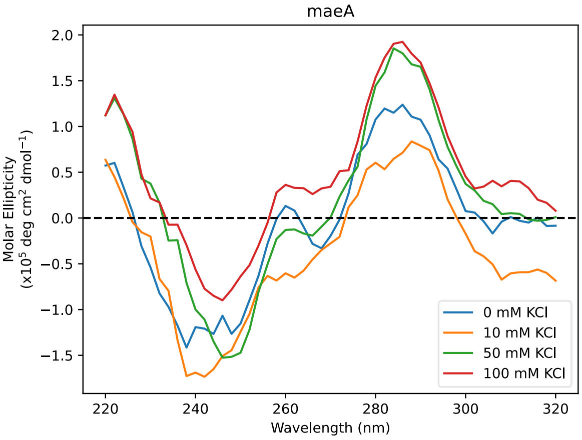


### Parallel → Antiparallel

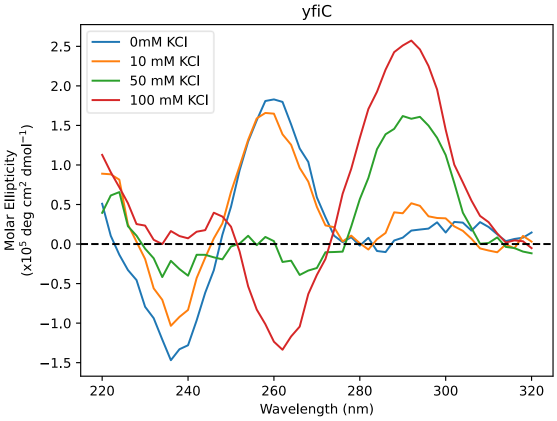


### Mix Hybrid - Parallel

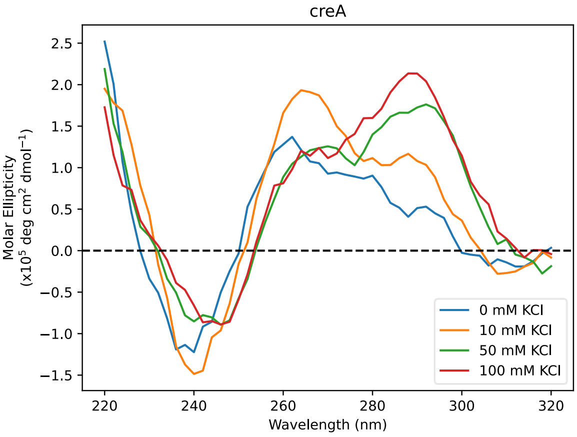


**Figure S7. CD spectra of candidate G4 DNA sequences at different KCl concentrations (0-100 mM).**

### 6. FRET melting assays

FRET melting assays were performed to assess ligand affinity for duplex and G-quadruplex DNA. Briefly, oligonucleotides of interest were obtained labelled at the 5' and 3' ends with FAM (a fluorescence donor) and TAMRA (a fluorescence quencher), respectively. In the folded state, proximity of the donor and quencher result in no observed fluorescence from FAM, since energy is transferred non-radiatively to TAMRA by FRET. As the temperature is raised and the secondary structure denatures, the fluorophores move further apart and the fluorescence signal increases. From the resulting curve, the characteristic melting temperature ($T_{max}$, also referred to as $T_m$) is defined as the temperature which corresponds to the maxima of the first derivative of the normalised fluorescence signal. The change in melting temperature ($\Delta T_m$) induced by the presence of a small molecule ligand provides an indication of the ligand's ability to stabilise the DNA structure. FRET experiments were performed according to the procedure reported by De Cian and co-workers[24] on a Stratagene MX3005P qPCR instrument. The method consisted of holding at 25 °C for 5 min, before heating at 1 °C/min to 96 °C in 1 °C increments, followed by monitoring the fluorescence output at each increment for 1 min. The fluorescence emission of FAM was followed at 516 nm, with a 10 nm full width at half-maximum filter and an 8-fold gain, after excitation at 492 nm with a 9 nm full width at half-maximum filter.

All oligonucleotides used (see **Table S5**) were purchased from Eurogentec (Belgium), purified by HPLC and delivered dry. Oligonucleotide concentrations were determined by UV-absorbance using a NanoDrop 2000 Spectrophotometer from Thermo Scientific. All sequences were annealed before use by heating for 2 minutes at 90°C and then placed immediately into ice. The final concentration of oligonucleotide was 200 nM in 100 mM KCl in all cases. Ligand concentrations were 5 µM. Each sample was tested in duplicate on the same plate, and each plate was repeated in at least duplicate to assess the reproducibility of all results.

Appropriate control experiments were also carried out for each sample set. Data processing was carried out using Origin 9, with $\Delta T_{1/2}$ used to represent $\Delta T_m$.

**Table S6. Melting Temperature ($T_m$) for G4 oligonucleotides sequences identified in Table S5 at 100 mM KCl.**

| **$T_m$ (°C) of nucleotide sequences in the absence of ligands at (100 mM KCl)** | | | | | |
|---|---|---|---|---|---|
| **clpB** | 63.6 | **asnS** | 54.1 | **yfiC** | 45.1 |
| **dppA** | 50.6 | **yhaJ** | 58.5 | **creA** | 58.7 |
| **glnD** | 57.0 | **ybiO** | 62.7 | **mukE** | 53.2 |
| **hemL** | 59.0 | **aroL** | 39.5 | | |
| **pdxA** | 55.6 | **cobS** | 57.5 | | |
| **thrA** | 56.1 | **mtgA** | 54.3 | | |
| **yjbD** | 53.1 | **maeA** | 49.0 | | |

**Table S7. Thermal stabilization ($\Delta T_m$, °C) induced in G4 and duplex DNA at 5 μM concentrations of ligands L5, L9 and pyridostatin (PDS).**

| **Sequence** | **L5** | **L9** | **PDS** |
|---|---|---|---|
| **F10T** | 3.0 | 0.4 | N/A |
| **clpB** | 16.6 | 2.3 | 19.2 |
| **dppA** | 27.6 | 7.0 | 28.2 |
| **glnD** | 25.8 | 10.9 | 31.5 |
| **hemL** | 20.8 | 2.1 | 19.7 |
| **pdxA** | 16.2 | 9.4 | 28.1 |
| **thrA** | 25.2 | 12.2 | 29.5 |
| **yjbD** | 34.6 | 15.1 | 35.8 |
| **asnS** | 31.3 | 4.3 | 26.2 |
| **yhaJ** | 19.3 | 5.1 | 17.4 |
| **ybiO** | 25.9 | 4.9 | 19.6 |
| **aroL** | 32.8 | 12.7 | 22.8 |
| **cobS** | 31.2 | 16.7 | 31.2 |
| **mtgA** | 39.0 | 13.7 | 38.5 |
| **maeA** | 24.7 | 6.9 | 22.3 |
| **yfiC** | 16.8 | 6.5 | 21.1 |
| **creA** | 29.9 | 8.4 | 23.7 |
| **mukE** | 20.9 | 6.5 | 9.4 |

Comparison analysis of gene sequences between Potential G-Quadruplex Sequences (PQS, as determine by G4-Hunter and G4-iGrinder scores), their log2 fold-change in protein abundance (LogFC) and $\Delta T_m$ (as determined by FRET).

**Table S8. G4-Hunter and G4-iGrinder scores for the selected G4 and their LogFC.**

| | | | | L5 | | L9 | | PDS | |
|---|---|---|---|---|---|---|---|---|---|
| Gene | Protein Description | G4-Hunter | G4-iM Grinder | LogFC | ΔTm (°C) | LogFC | ΔTm (°C) | LogFC | ΔTm (°C) |
| ***clpB*** | Chaperone protein | 1.04 | | 0.06 | 16.6 | -1.13 | 2.3 | -1.03 | 19.2 |
| ***dppA*** | Periplasmic dipeptide transport protein | 1.3 | | -1.23 | 27.6 | -2.95 | 7.0 | -1.93 | 28.2 |
| ***glnD*** | Bifunctional uridylyltransferase | 1.08 | | 0.14 | 25.8 | 0.20 | 10.9 | 0.25 | 31.5 |
| ***hemL*** | Glutamate-1-semialdehyde 2,1-aminomutase | NA | | 0.28 | 20.8 | 0.44 | 2.1 | 1.74 | 19.7 |
| ***pdxA*** | 4-hydroxythreonine-4-phosphate dehydrogenase | 1.38 | | ND | 16.2 | -1.98 | 9.4 | -0.59 | 28.1 |
| ***thrA*** | Bifunctional aspartokinase/homoserine dehydrogenase | 1.2 | | -0.36 | 25.2 | 1.36 | 12.2 | 0.03 | 29.5 |
| ***yjbD*** | Uncharacterised protein | 1.44 | | ND | 34.6 | -1.32 | 15.1 | -1.14 | 35.8 |
| ***asnS*** | Asparagine-tRNA ligase | | 30 | -0.19 | 31.3 | 1.62 | 4.3 | 0.35 | 26.2 |
| ***yhaJ*** | Hypothetical transcriptional regulator | | 47 | -0.41 | 19.3 | 1.03 | 5.1 | 0.17 | 17.4 |
| ***ybiO*** | Mechanosensitive channel | | 40 | ND | 25.9 | ND | 4.9 | ND | 19.6 |
| ***aroL*** | Shikimate kinase 2 | | 30 | -1.63 | 32.8 | 1.22 | 12.7 | 0.03 | 22.8 |
| ***cobS*** | Adenosylcobinamide-GDP ribazoletransferase | | 41 | ND | 31.2 | 1.74 | 16.7 | ND | 31.2 |
| ***mtgA*** | Biosynthetic peptidoglycan transglycosylase | | 40 | 2.81 | 39.0 | -1.07 | 13.7 | -0.39 | 38.5 |
| ***maeA*** | NAD-dependent malic enzyme | | 34 | 0.74 | 24.7 | -1.06 | 6.9 | 1.46 | 22.3 |
| ***yfiC*** | tRNA1(Val) (adenine(37)-N6)-methyltransferase | | 37 | ND | 16.8 | 1.78 | 6.5 | ND | 21.1 |
| ***creA*** | Catabolite regulation protein A | | 30 | -2.45 | 29.9 | 1.52 | 8.4 | 0.07 | 23.7 |
| ***mukE*** | Chromosome partition protein | | 40 | ND | 20.9 | 1.58 | 6.5 | 0.23 | 9.4 |

LogFC, log2 fold-change in protein abundance (treated vs non-treated); $\Delta T_m$, difference in FRET melting temperatures (treated vs non-treated). NA, Not available; ND, Not Detected. Grey numbers indicates that value is not significant ($p > 0.05$, t-test).

### 7. UV-visible spectroscopy

UV spectra were recorded on an Agilent Cary 60 UV-Vis Spectrophotometer at ambient temperature using a room-light immune fiber optic probe with 10 mm path length at ambient temperature. Measurements were taken in a quartz cuvette with a path length of 10 mm using in slow scanning mode at 1 nm intervals. The UV-visible spectra were recorded between 700 nm and 300 nm and baseline corrected for the buffer used.

Determination of apparent dissociation constants: Apparent dissociation constants were determined through UV-visible spectroscopy titration experiments. The raw spectra were recorded as described above. The concentration of ligand was fixed at 10 µM in a constant volume of 0.5 mL buffer. The buffer used was potassium phosphate (100 mM, pH 7.4). During the titration, aliquots of oligonucleotide were added to give the required titration points (from a 100 µM stock solution in appropriate buffer containing also 10 µM ligand to maintain constant ligand concentration). NB: the oligonucleotide solution was annealed by heating to 90 °C for 2 minutes and then cooling on ice prior to the addition of ligand (to avoid annealing in the presence of ligand). Following addition, the solution was mixed thoroughly and the UV-visible spectrum was acquired immediately. Data were fitted to an independent-and equivalent-sites binding model using Prism 7 software.

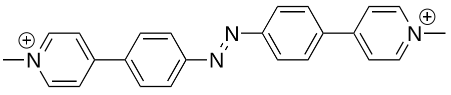

**L9**

### aroL

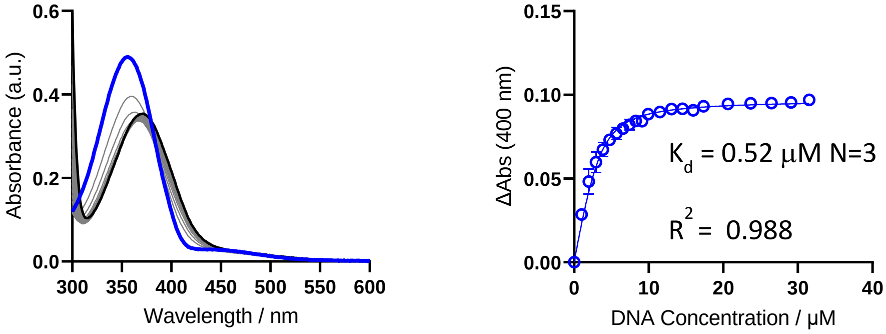


### mtgA

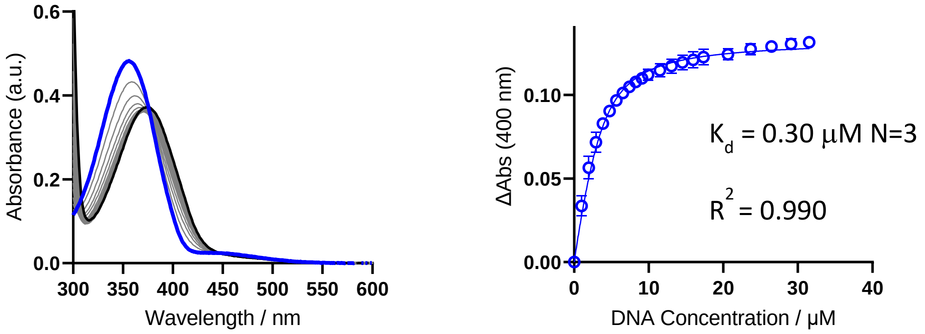


### cobS

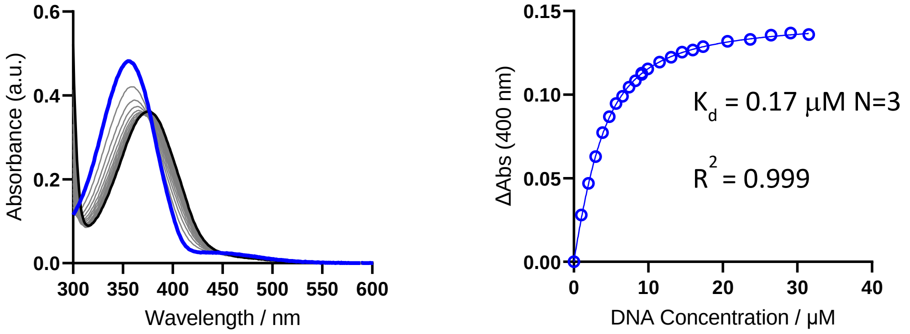


**Figure S8A. UV-Vis titration curves and determination of apparent binding constant ($K_d$) for L9 (0-30 µM) and the different *E. coli* oligonucleotide sequences *aroL*, *mtgA* and *cobS* in 100 mM potassium phosphate buffer.**

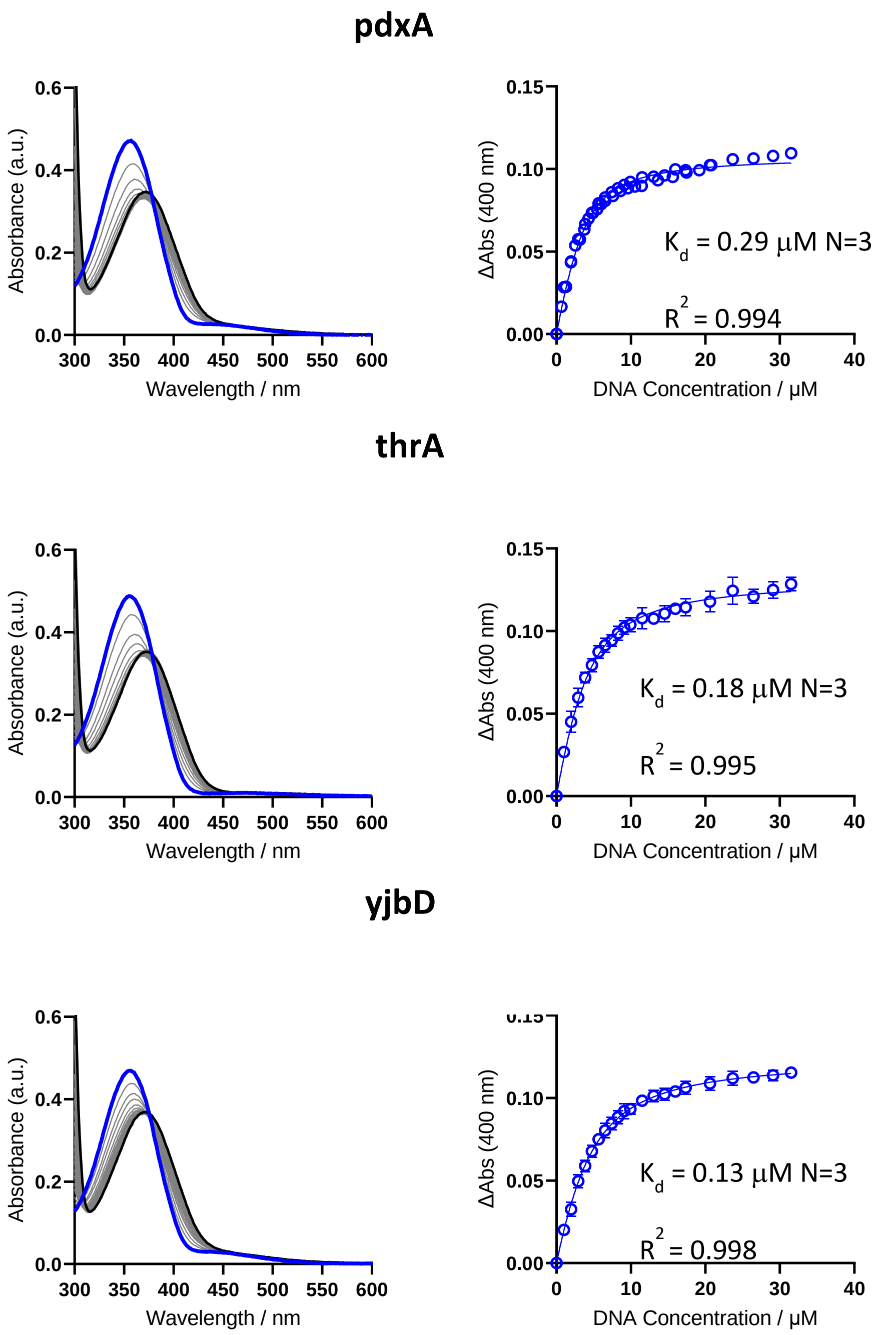


**Figure S8B. UV-Vis titration curves and determination of apparent binding constant ($K_d$) for L9 (0-30 µM) and the different *E. coli* oligonucleotide sequences *thrA*, *yjbD* and *pdxA* in 100 mM potassium phosphate buffer.**

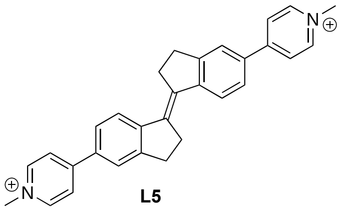

L5

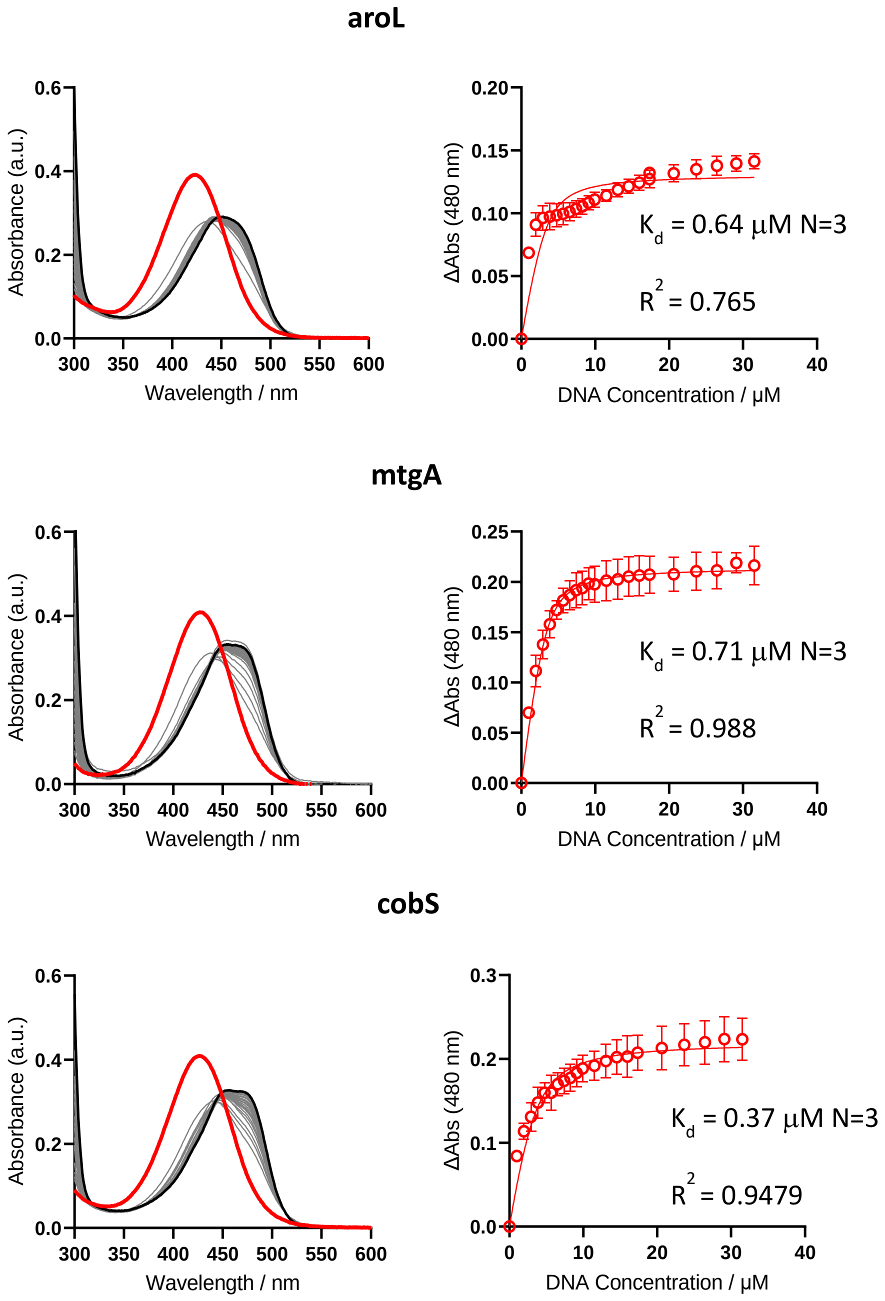


**Figure S9A. UV-Vis titration curves and determination of apparent binding constant ($K_d$) using UV-Vis titration of L5 (0-30 μM) into the different *E. coli* oligonucleotide sequences in 100 mM potassium phosphate buffer.**

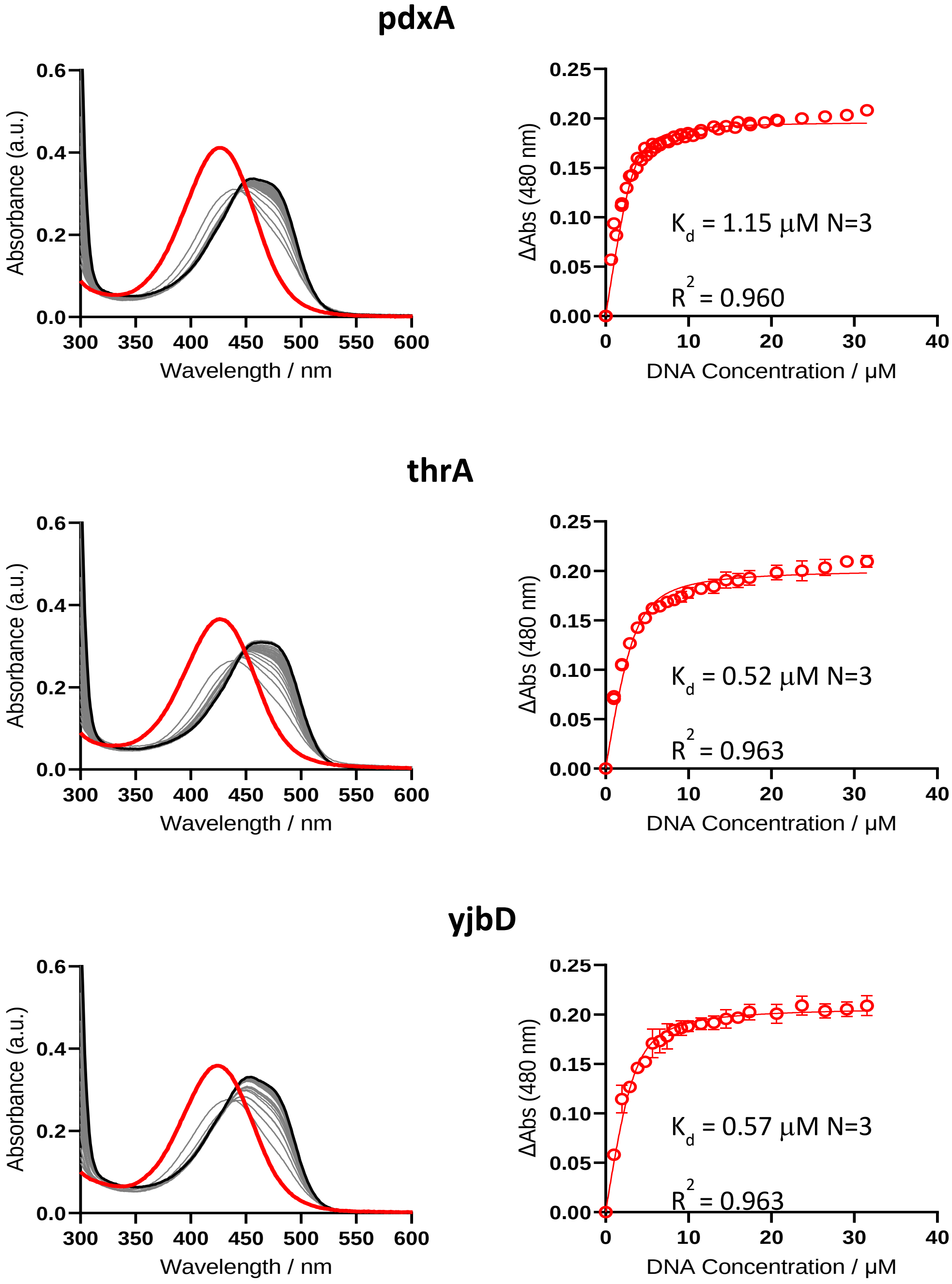


**Figure S9B. UV-Vis titration curves and determination of apparent binding constant ($K_d$) for L5 (0-30 µM) and the different *E. coli* oligonucleotide sequences *thrA*, *yjbD* and *pdxA* in 100 mM potassium phosphate buffer.**

### 8. Circular Dichroism Titrations of L5, L9 and pyridostatin to *pdxA, thrA*, *yjbD, aroL, cobS,* and *mtgA* oligonucleotide sequences

Circular Dichroism (CD) titrations were recorded using a Jasco J-810 spectrometer fitted with a Peltier temperature controller. Measurements were taken in a quartz cuvette with a path length of 5 mm, at 20 °C, at a 1000 nm / min scanning speed at 1 nm intervals, with a 1 nm bandwidth. The CD spectra were recorded between 600 and 220 nm, and baseline corrected for the buffer used. Oligonucleotide concentrations were determined by UV-absorbance using a NanoDrop 2000 Spectrophotometer from Thermo Scientific. The oligonucleotide was annealed before use by heating for 2 minutes at 90°C and then placed immediately into ice. The oligonucleotide was at a concentration of 4.22 µM which gave an OD of 1 and the buffer used was potassium phosphate (100 mM, pH 7.4). Oligonucleotide concentration remained constant throughout, and dilutions were made using a solution containing both oligonucleotide and the ligand, with the amount added determining the relative ratio of the two species. The reported spectrum for each sample represents the average of 3 scans. Data processing was carried out using Prism 7. Observed ellipticities were converted to mean residue ellipticity (θ) = deg $cm^2$ $dmol^{-1}$ (molar ellipticity).

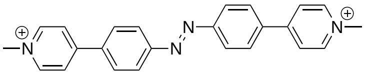

**L9**

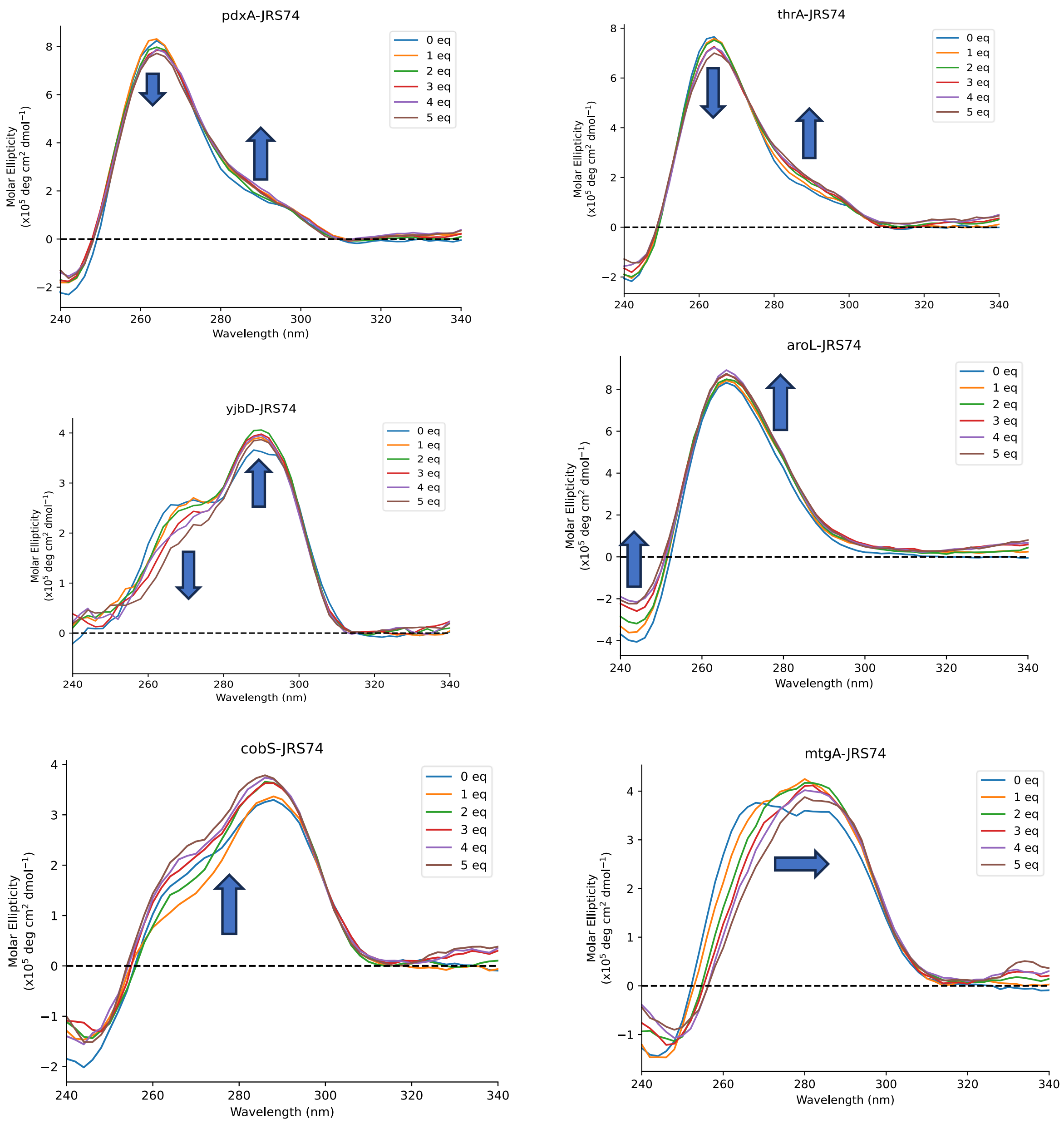


**Figure S9. CD spectra of titration of L9 into the different *E. coli* oligonucleotide sequences in 100 mM potassium phosphate buffer.**

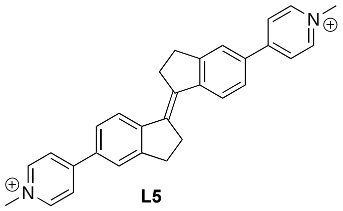

**L5**

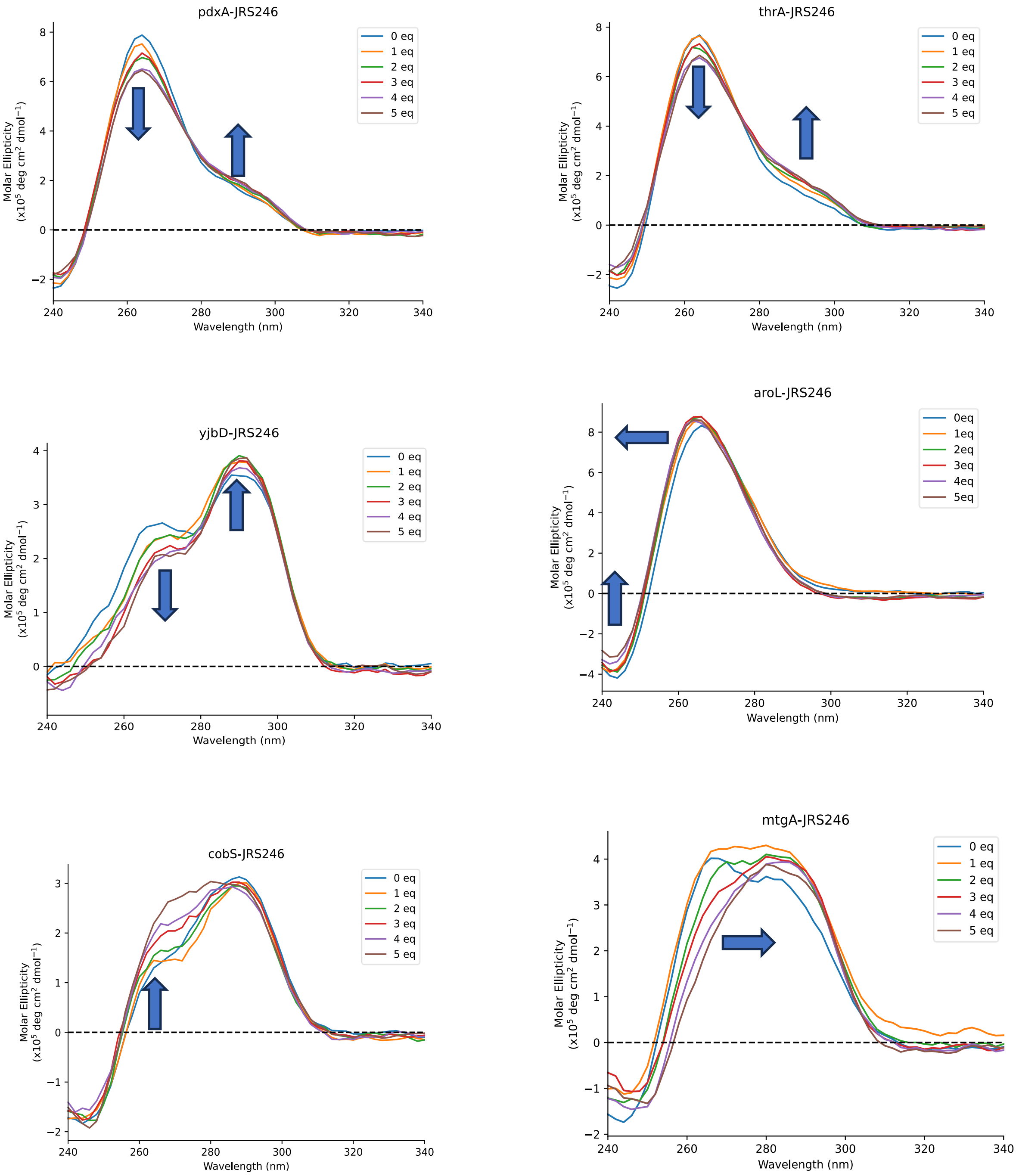


**Figure S10. CD spectra of titration of L5 into the different *E. coli* oligonucleotide sequences in 100 mM potassium phosphate buffer.**

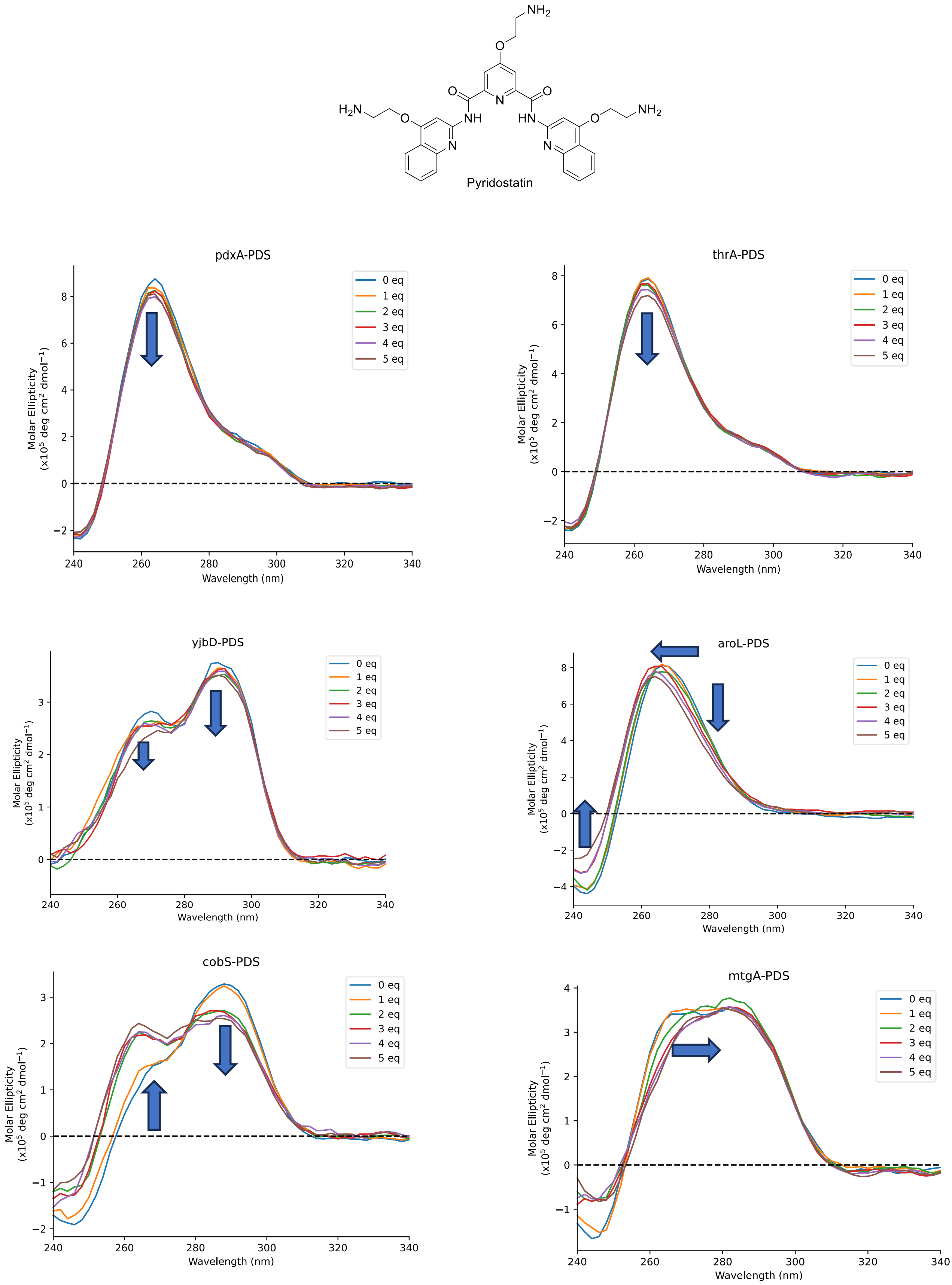


**Figure S11. CD spectra of titration of pyridostatin into the different *E. coli* oligonucleotide sequences in 100 mM KCl buffer.**

### 9. Bacterial Cytological Profiling (BCP) image acquisition and analysis

All samples in biological triplicates were treated with 1x MIC concentration of **L9** or representative antibiotic for 2 hours and prepared as previously described.[25] All images were acquired on the Leica TCS SP8 system attached to a Leica DMi8 inverted microscope (Leica Microsystems) using a 63x/1.4 HC PL APO CS2 oil immersion objective. The samples were excited using a 405 nm diode laser for detecting DAPI (ThermoFisher) fluorescence (over an emission range of 415-470 nm), a DPSS 561 laser for detecting FM4-64FX (ThermoFisher) (emission range 570-650 nm) and an argon laser for SYTOX-Green (ThermoFisher) (emission range 495-550 nm). Confocal Z stacks were taken with the pinhole set to 1AU and a step size of 228 nm and the pixel dwell time was 0.36 us.

Image analysis was performed as previously described.[26, 27] For image data extraction, the raw images from the fluorescent microscope were preprocessed using ImageJ.[28] Subsequently, morphological features such as length, area, perimeter, and fluorescent intensity were measured using CellProfiler 4.0 software[29] (7) followed by image analysis. Briefly, the cytological profiles were transformed using Quantile Transformer, and the morphological features were selected through recursive feature elimination with cross-validation (RFECV-SVM).[30] (8). Lastly, the data dimension reduction and visualization were performed through a non-linear dimensional reduction method called pairwise controlled manifold approximation (PaCMAP).[31]
